# Revealing essential properties for enhanced quantum sensing in engineered *As*LOV2 proteins

**DOI:** 10.64898/2026.02.27.708212

**Authors:** Lewis M. Antill, Joseph K. Baidoo, Luca Gerhards

## Abstract

Genetically encoded flavoproteins have emerged as optically readable radical pair quantum sensors, but the molecular basis of the differing magnetic responses of closely related variants remains unresolved. Here, we analyse parental *As*LOV2 and three evolved MagLOV variants using molecular dynamics, quantum chemical hyperfine calculations, Marcus electron transfer theory, and trajectory-based spin relaxation analysis. The mutations preserve the LOV fold and FMN-binding core but progressively reshape the conformational landscape of the W89 electron donor, with broader rigid-body libration yet increasingly restricted internal torsional motion and predominant occupation of a single rotameric basin in MagLOV2f. These distinct motions have different magnetic consequences: dipolar coupling modulation increases strongly in the later variants, whereas hyperfine modulation rates vary non-monotonically and decrease from MagLOV2 to MagLOV2f. The mutations also alter the radical pair free energy landscape and donor–acceptor orientation, producing variant-dependent back electron transfer rates, with MagLOV2f combining comparatively fast calculated recombination with reduced hyperfine relaxation relative to MagLOV2. Thus, directed evolution redistributes competing recombination and relaxation pathways rather than optimising a single kinetic parameter. This molecular picture rationalises the distinct magnetic-response amplitudes and saturation dynamics of closely related MagLOV proteins and identifies donor conformational control as a practical design variable for protein-based radical pair quantum sensors.

## 1 Introduction

Quantum sensing exploits quantum degrees of freedom to convert weak environmental perturbations into measurable signals. In chemistry, one particularly direct realisation is spin-dependent reactivity: when reaction pathways depend on spin evolution, weak magnetic fields can alter product yields and thereby generate a chemically encoded magnetic response. Photoinduced radical pairs provide a paradigmatic example, in which coherent singlet–triplet interconversion and spin-selective reaction pathways translate spin dynamics into changes in chemical yield [1–3]. Embedding this functionality in genetically encoded proteins could enable spatially targeted and optically readable quantum sensors that operate directly within biological environments [4–7].

Solid-state defects, most prominently nitrogen-vacancy (NV) centres in diamond, have established a powerful platform for magnetic field and temperature sensing, combining high sensitivity with optical initialisation and readout [8, 9]. NV-based probes have enabled thermometry in living cells and *in vivo* [10, 11]. Their broader implementation in complex biological systems, however, requires the delivery and localisation of an inorganic material, surface functionalisation, and coupling of the sensor to the biological target of interest [12, 13]. Genetically encoded probes based on spin-dependent photochemistry, therefore, represent a complementary strategy in which the sensing element can be produced and positioned directly by the cell [4–6].

Flavins provide a natural molecular platform for such protein-based spin chemistry. As ubiquitous redox-active cofactors, they access oxidised, semiquinone, and reduced states and participate in light-driven electron transfer reactions across diverse protein environments [14]. Flavin-centred radical pairs can exhibit magnetic field effects under ambient conditions [15–20], with cryptochromes providing the best-known biological example and a leading candidate mechanism for magnetoreception [21, 22]. Nevertheless, translating flavin radical pair chemistry into a practical intracellular sensor remains challenging because magnetic sensitivity depends not only on radical-pair formation but also on competing reaction kinetics, radical stability, molecular geometry, and spin relaxation within the surrounding protein environment [21, 23–26]. These requirements motivate the development of compact and engineerable flavoprotein scaffolds in which radical pair formation and spin dynamics can be systematically controlled.

Phototropin light–oxygen–voltage (LOV) domains provide such a scaffold. The LOV2 domain of *Avena sativa* phototropin 1 binds flavin mononucleotide (FMN) non-covalently and couples blue light absorption to a conformational response [27]. In the wild type, photoexcitation leads to the formation of a reversible cysteinyl–FMN adduct. Mutation of the reactive cysteine (C48A, Fig. 1a) suppresses this canonical photocycle and promotes flavin radical chemistry while retaining useful photophysical properties [28–32]. LOV-based fluorescent proteins can additionally function under low-oxygen conditions, unlike conventional GFP-based fluorophores, making them attractive genetically encoded reporters for environments in which oxygen availability is limited [33, 34]. More broadly, recent studies using both EYFP- and LOV-derived systems have demonstrated the emerging feasibility of genetically encoded probes whose optical response is coupled to spin-dependent processes [4, 7, 35].

**Fig. 1.**
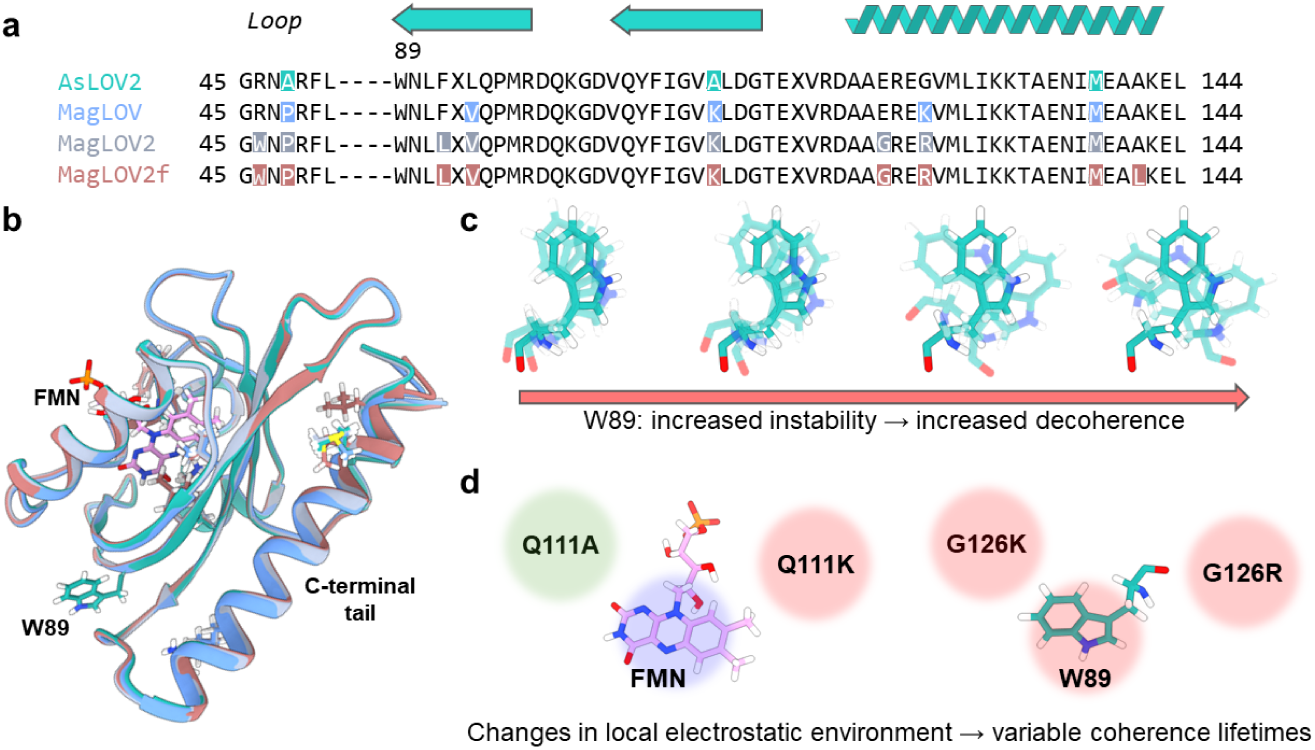
Overview of the MagLOV mutation series and the proposed FMN–W89 radical-pair environment. **a** Sequence alignment with the engineered positions highlighted for the four constructs. **b** Structural overlay showing the conserved LOV fold, FMN cofactor, W89 donor, and C-terminal region. **c** Representative W89 conformations illustrating the mutation-sensitive orientational ensemble. The arrow denotes the experimentally motivated schematic progression used to frame the study; the trajectory analysis below resolves this motion into distinct librational and internal-torsional components rather than a single scalar “instability”. **d** Mutation-dependent changes near FMN and W89 that alter the local electrostatic and conformational environment and can thereby modulate recombination and spin relaxation.

The recently developed MagLOV proteins provide a particularly striking realisation of this concept [5]. Derived from *Avena sativa* LOV2 (Fig. 1a,b), variants obtained through mutagenesis and directed evolution exhibit magnetic field effects (MFEs) and optically detected magnetic resonance (ODMR) under physiological conditions and in living cells [5]. These responses were attributed to a spin-correlated radical pair (SCRP) involving FMN and a nearby tryptophan donor (Fig. 1b–d). Importantly, the different MagLOV variants exhibit markedly different magnetic responses despite sharing the same underlying flavin-based photochemistry. Because these variants arose from largely stochastic mutational trajectories, it remains unclear which molecular changes are responsible for these differences and, more generally, which properties of the protein environment govern efficient radical pair-based quantum sensing.

Resolving this question requires connecting protein sequence and structure to both radical pair kinetics and spin relaxation. Mutations can alter the stability and conformational dynamics of the electron donor, change donor–acceptor geometry and electronic coupling, reshape the local electrostatic environment, and modify fluctuations of the magnetic interactions that drive spin relaxation. These effects act simultaneously and can either enhance or suppress the lifetime and magnetic sensitivity of the SCRP. A molecular description of MagLOV, therefore, requires treating radical pair formation and termination together with the structural fluctuations that determine spin coherence.

Here, we establish a mechanistic description of the four *AsLOV2-derived* constructs characterised experimentally—the parental construct, MagLOV, MagLOV2, and MagLOV2f (MagLOV2 fast)—by combining molecular dynamics simulations, quantum chemical calculations, Marcus electron transfer theory, and spin relaxation analysis. Rather than assuming that directed evolution changes a single notion of donor flexibility, we resolve distinct rigid-body and internal motions of W89 and determine how each is propagated into fluctuations of the electron–electron dipolar and electron–nuclear hyperfine interactions. We further quantify mutation-dependent changes in the radical pair free-energy landscape and donor–acceptor geometry that regulate back electron transfer. The resulting picture is one of kinetic redistribution: the variants occupy different balances of dipolar modulation, hyperfine modulation, and recombination, providing a molecular rationale for why selection for MFE magnitude and selection for MFE saturation rate produce distinct protein dynamics [5].

## 2 Results and Discussion

### 2.1 Global structural differences

To establish whether the introduced mutations alter the LOV scaffold itself or predominantly act through local dynamics, we first compared the global structural ensembles of the four variants. The LOV2 fold remains intact in all simulations, with the largest fluctuations confined to surface-exposed segments: the N terminus (residues 0–10), loop regions around residues 16–20 and 38–41, the solvent-exposed segment around residues 98–101, and the J*α*/C-terminal region (approximately residues 115–125; Fig. 2a). The same pattern is apparent in the per-residue root-mean squared fluctuation (RMSF) profiles (Fig. 2c), while the backbone root-mean squared deviation (RMSD) trajectories remain bounded over the simulated timescale. Replica-resolved RMSD and RMSF data, including the 900 ns extension used to test the MagLOV2f conformational change, are reported in Section S2 of the SI. Thus, the mutations change local conformational populations without producing a systematic loss of the LOV fold or the FMN-binding pocket.

**Fig. 2.**
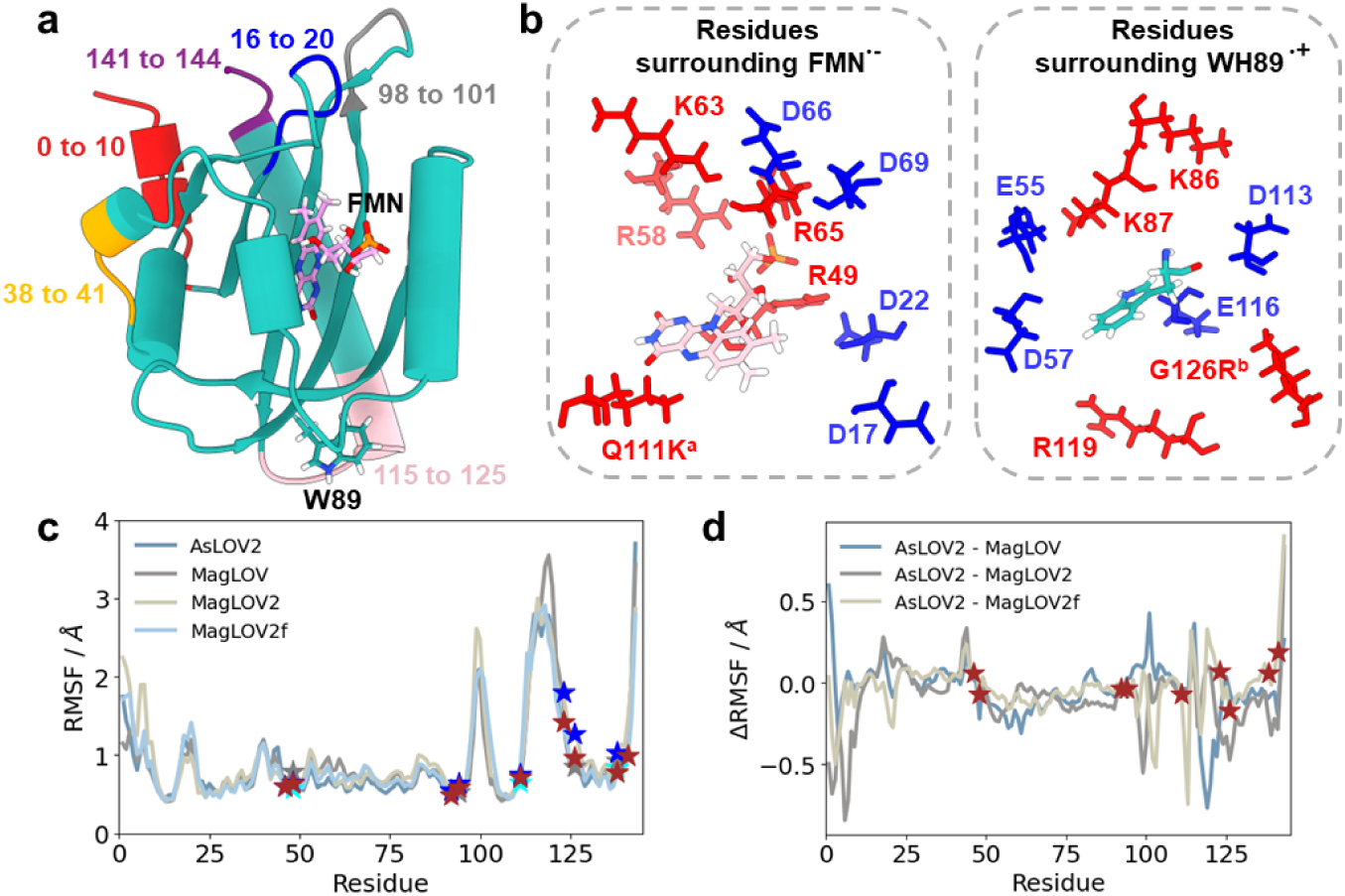
Global structural stability and charged microenvironments surrounding the proposed radical pair. **a** LOV2 structure with the principal flexible surface segments highlighted. **b** Charged residues surrounding FMN*^•−^* (left) and W89H*^•^*^+^ (right). Red and blue denote positively and negatively charged side chains, respectively. The labels Q111K and G126R indicate mutation-dependent changes in the local environment; Q111A in the parental construct is neutral. **c** Per-residue RMSF for all four variants, with engineered positions marked by stars. **d** RMSF differences between *As*LOV2 and MagLOV, MagLOV2, or MagLOV2f.

This distinction is important because the flavin photochemistry is known to be sensitive to its local pocket, even when the overall LOV structure is retained [29, 36]. The engineered Cys-to-Ala substitution (C48A in the construct numbering used in Fig. 1; C450A in full-length phototropin numbering) suppresses canonical cysteinyl–FMN adduct formation while retaining the LOV scaffold [29], and substitutions at the conserved Q111 position can substantially alter LOV recovery kinetics without abolishing photoswitching [36, 37]. In the MagLOV lineage, Q111K additionally changes the electrostatic potential and hydrogen-bonding character near FMN, while G126K/G126R introduces a positively charged side chain close to the W89 donor region (Figs. 1d and 2b). The FMN*^•−^* pocket contains both positively and negatively charged residues, whereas W89H*^•^*^+^ is surrounded by a distinct, similarly heterogeneous charge environment (Fig. 2b). We therefore interpret these mutations as changes to the local electrostatic and conformational topography, rather than as a simple global stabilisation or destabilisation of the protein.

Solvent-accessible surface areas (SASA) support the same conclusion. Neither the total protein SASA nor the FMN or W89 SASA shows a monotonic change across the evolutionary series, although individual variants and replicas populate distinct solvent-exposure substates (Section S3 of the SI). Accordingly, solvent exposure can contribute to conformational gating but does not, by itself, provide a one-dimensional explanation for the progression of magnetic phenotypes. Hydrogen-bond analysis likewise shows that FMN is persistently embedded in a hydrogen-bonded pocket, whereas W89 has substantially fewer persistent hydrogen bonds (Section S5 of the SI), making the donor more susceptible to mutation-dependent packing and librational changes.

Finally, the RMSF differences relative to *As*LOV2 are modest and spatially localised (Fig. 2d). Several engineered positions lie in or near regions whose fluctuations shift from the parental ensemble, consistent with directed evolution redistributing local packing and long-range dynamical coupling while conserving the overall protein architecture.

### 2.2 Local structural differences and spin relaxation

Because the proposed SCRP contains a comparatively rigid FMN acceptor and the W89 donor, we next separated rigid-body libration from internal torsional motion for both radicals (Fig. 3). This distinction is essential: an increase in one coordinate does not imply a general increase in molecular flexibility, and different coordinates modulate different terms of the spin Hamiltonian.

**Fig. 3.**
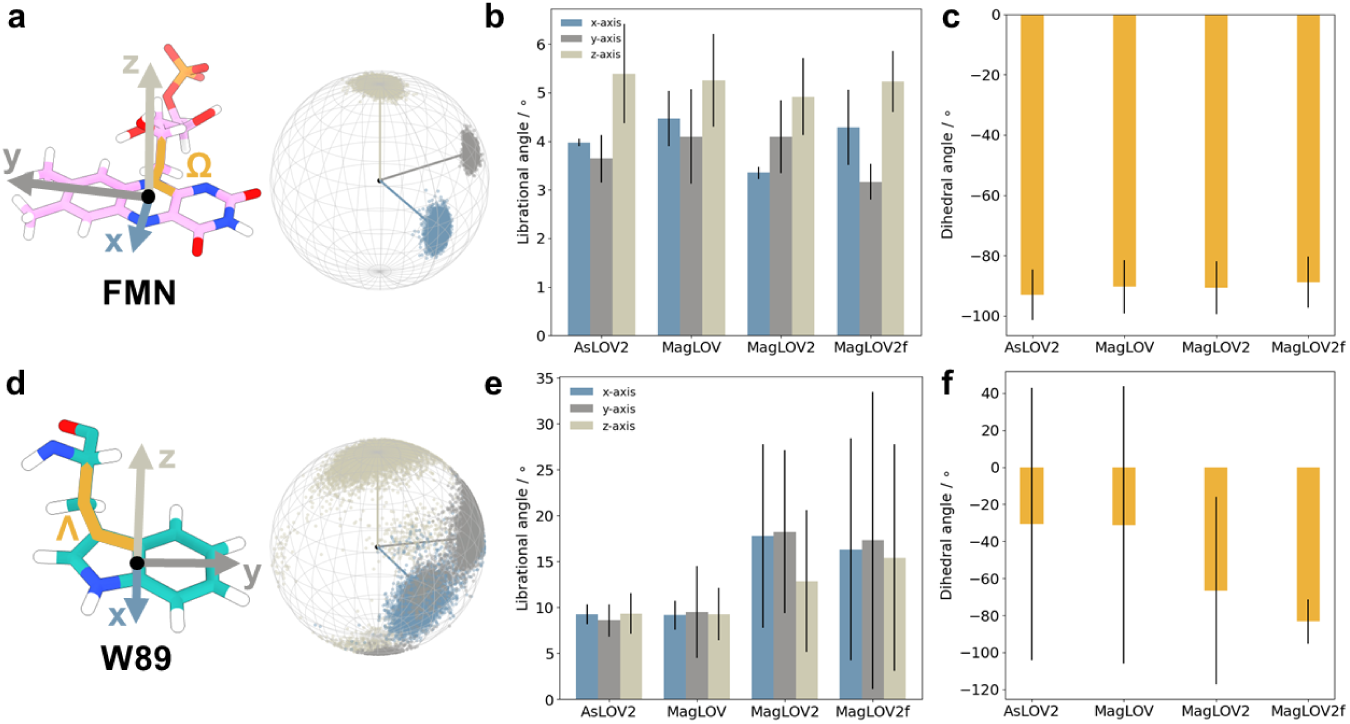
Molecular fluctuations of the proposed SCRP partners. **a,d** Molecular structures and local Cartesian axes used to describe FMN (a) and W89 (d) libration; the representative internal dihedrals Ω and Λ are highlighted. **b,e** Axis-resolved librational angles for FMN (b) and W89 (e) across *As*LOV2, MagLOV, MagLOV2, and MagLOV2f. FMN libration changes only modestly, whereas W89 rigid-body libration is markedly broader in the later evolved variants. **c,f** Representative FMN (c) and W89 (f) dihedral angles with their trajectory-derived spread. FMN remains narrowly distributed. In contrast to the librational trend, the W89 torsional ensemble becomes progressively restricted in MagLOV2 and especially MagLOV2f.

FMN*^•−^* remains conformationally constrained in every variant. Its spherical molecular-axis distributions are compact, the mean librational angles are similar across the four constructs, and the representative FMN dihedral Ω remains narrowly distributed (Fig. 3a–c). Replica-resolved librational distributions and spherical plots are given in Section S4 of the SI. These data show that directed evolution does not substantially loosen the isoalloxazine binding geometry; FMN therefore acts as a relatively rigid anchor for the radical pair. W89H*^•^*^+^ behaves differently. The W89 molecular-axis distributions broaden strongly in MagLOV2 and MagLOV2f, and the axis-resolved librational amplitudes increase from the parental construct to the later evolved variants (Fig. 3d,e). The full replica-resolved distributions are non-Gaussian and, in several cases, multimodal (Section S4 of the SI), indicating exchange among distinct donor orientations rather than a single harmonic fluctuation. This enhanced rigid-body sampling is consistent with the sparse W89 hydrogen-bond network in Section S5 and identifies the donor pocket as the principal source of mutation-sensitive orientational dynamics.

The internal W89 torsion shows the opposite evolutionary trend. Although W89 is intrinsically more torsionally heterogeneous than FMN, the broad/two-basin distribution present in *As*LOV2 and MagLOV becomes less populated at the secondary rotamer in MagLOV2 and is strongly concentrated in one rotameric basin in MagLOV2f (Fig. 3f and Section S4 of the SI). Thus, the later mutations broaden W89 rigid-body libration while narrowing its internal torsional landscape. The accompanying distribution of the angle between the FMN and W89 aromatic-plane normals also shifts, with MagLOV2f preferentially sampling lower angles (Section S4 of the SI). This separation of motional coordinates provides a structural basis for the non-monotonic relaxation behaviour discussed below.

Exchange modulation was evaluated using the distance-dependent model described in Section S6 of the SI and produced a substantially smaller fluctuation integral than the electron–electron dipolar contribution. We therefore focus on dipolar (*k_D_*) and HFC (*k*_HFC_) modulation as the dominant mutation-sensitive random fields in the present model. The FMN–W89 COM (centre of mass) distance itself changes only weakly in its ensemble mean (approximately 15.30–15.41 5), but its distribution and replica dependence vary among constructs (Section S7 of the SI). This observation is important because the relaxation trend cannot be explained by a simple monotonic shortening of the radical pair distance.

Figure 4b shows the effective diagonal dipolar modulation coefficients derived from the tensor autocovariances. *As*LOV2 has the smallest values, MagLOV shows only a modest increase, and both MagLOV2 and MagLOV2f show substantially enhanced dipolar modulation, particularly for the *zz* component. The large error bars reflect genuine replica-to-replica heterogeneity rather than uncertainty from a single fit. The corresponding dipolar auto-correlation functions (Section S8 of the SI) show short correlation times for *As*LOV2 (1.8–5.4 ns), a broader range for MagLOV (0.92–11.2 ns), and slowly decorrelating replicas in MagLOV2 and MagLOV2f reaching approximately 44 and 55 ns, respectively. Because *k_D_* depends on both the tensor variance and its correlation time, these persistent donor conformations contribute strongly only when they also carry appreciable dipolar variance. Together with the librational analysis, the data identify W89 rigid-body reorientation as a major source of the enhanced dipolar random field in the later variants.

**Fig. 4.**
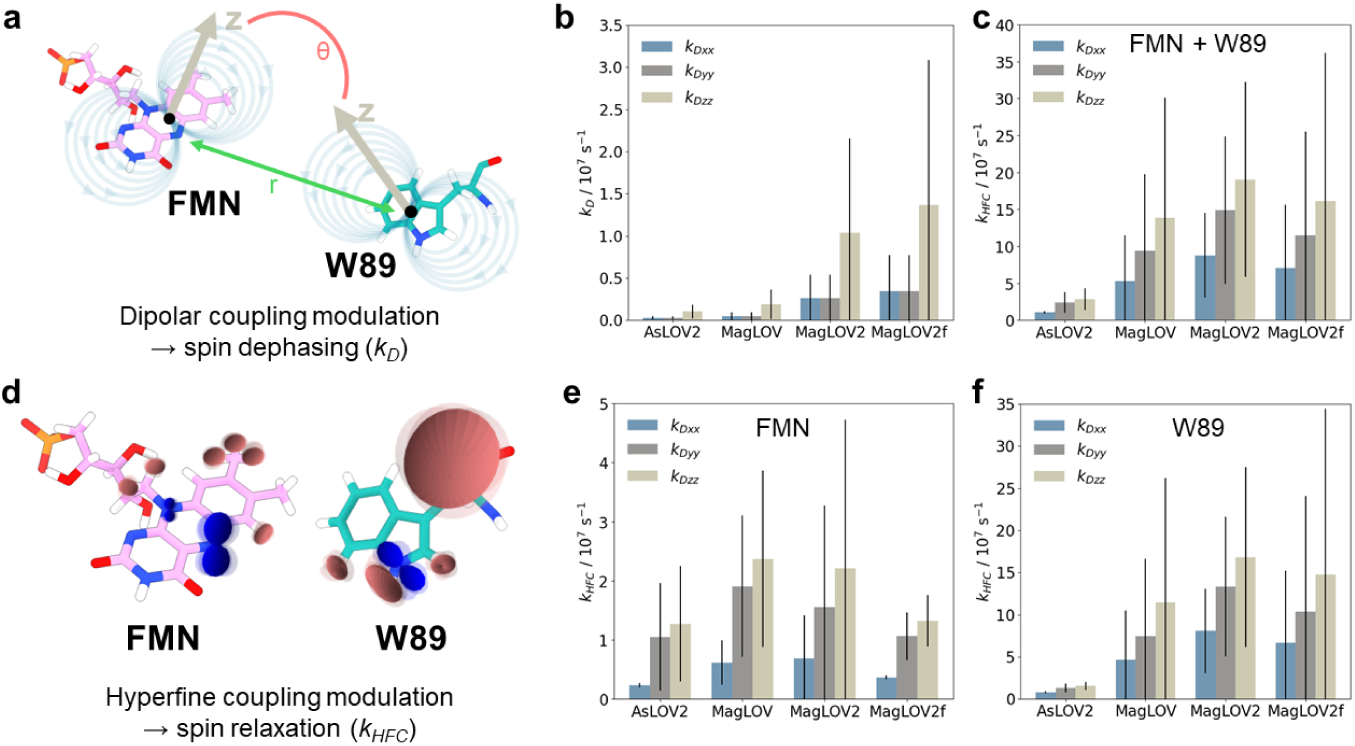
Mutation-dependent dipolar and hyperfine modulation of the proposed FMN–W89 radical pair. **a** Schematic of electron–electron dipolar coupling modulation by fluctuations of the inter-radical vector and relative orientation. **b** Effective diagonal dipolar modulation coefficients *k_D,xx_*, *k_D,yy_*, and *k_D,zz_*for all four variants. Bars show the replica-averaged values and error bars the replica spread. **d** Schematic of HFC modulation by nuclear and conformational motion. **c** Effective HFC random-field coefficients for the combined FMN+W89 nuclear-spin set. **e,f** Corresponding decomposition into FMN (e) and W89 (f) contributions. The HFC contribution is largest in MagLOV2 and decreases in MagLOV2f, whereas the dipolar contribution remains high in both later variants.

HFC modulation provides a complementary and non-monotonic contribution (Fig. 4c,e,f). For the combined FMN+W89 nuclear-spin set, the HFC random-field coefficients increase strongly from *As*LOV2 to MagLOV and reach their largest values in MagLOV2, whereas MagLOV2f decreases again relative to MagLOV2 while remaining above the parental level (Fig. 4c). The decomposition shows that W89 contributes the larger absolute HFC-modulation term (Fig. 4f), although FMN also follows a variant-dependent trend (Fig. 4e). This behaviour is consistent with the structural data: the later evolutionary steps do not simply make the donor “more flexible” but simultaneously broaden rigid-body orientations and restrict internal torsional sampling. The narrowing of the W89 rotameric landscape in MagLOV2f therefore provides a plausible structural contribution to the reduction in *k*_HFC_ relative to MagLOV2, while the remaining HFC differences also reflect changes in local geometry and electrostatics. All isotropic HFC trajectories, full-tensor fluctuation profiles, nuclear labels, and the common nucleus-selection procedure are documented in Section S8 of the SI.

The two relaxation channels, therefore, respond differently to the same evolutionary trajectory. MagLOV2 and MagLOV2f both access stronger dipolar modulation than the parental system, but MagLOV2f partially suppresses the HFC contribution relative to MagLOV2. This redistribution matters for radical pair quantum sensing because relaxation is not universally detrimental: depending on the kinetic regime, moderate relaxation can alter or even enhance a radical pair magnetic response, whereas sufficiently rapid relaxation erases the spin correlations required for field sensitivity [38, 39]. The relevant quantity is consequently the balance of relaxation with spin-selective reaction kinetics, rather than the minimisation of any one coefficient in isolation.

### 2.3 Recombination rates

The radical pair lifetime is determined not only by spin relaxation but also by the rate at which the charge-separated state recombines. We therefore constructed Marcus free-energy profiles for RP→GS back electron transfer (BET) from the energy-gap distributions of all four variants (Fig. 5a; full distributions in Section S9 of the SI). The calculated reaction free energies are strongly exergonic in every case: Δ*G^◦^*= −64.7, −67.1, −63.2, and −75.2 kcal mol*^−^*^1^ for *As*LOV2, MagLOV, MagLOV2, and MagLOV2f, respectively. The corresponding reorganisation energies are *λ_r_* = 74.5, 70.3, 73.3, and 76.8 kcal mol*^−^*^1^. Because |Δ*G^◦^*| is close to *λ_r_* for all four systems, the Marcus activation term is small and sensitive to comparatively modest shifts of either quantity. The energetic term alone is therefore not sufficient to explain the full rate ordering; donor–acceptor coupling and geometry remain important.

**Fig. 5.**
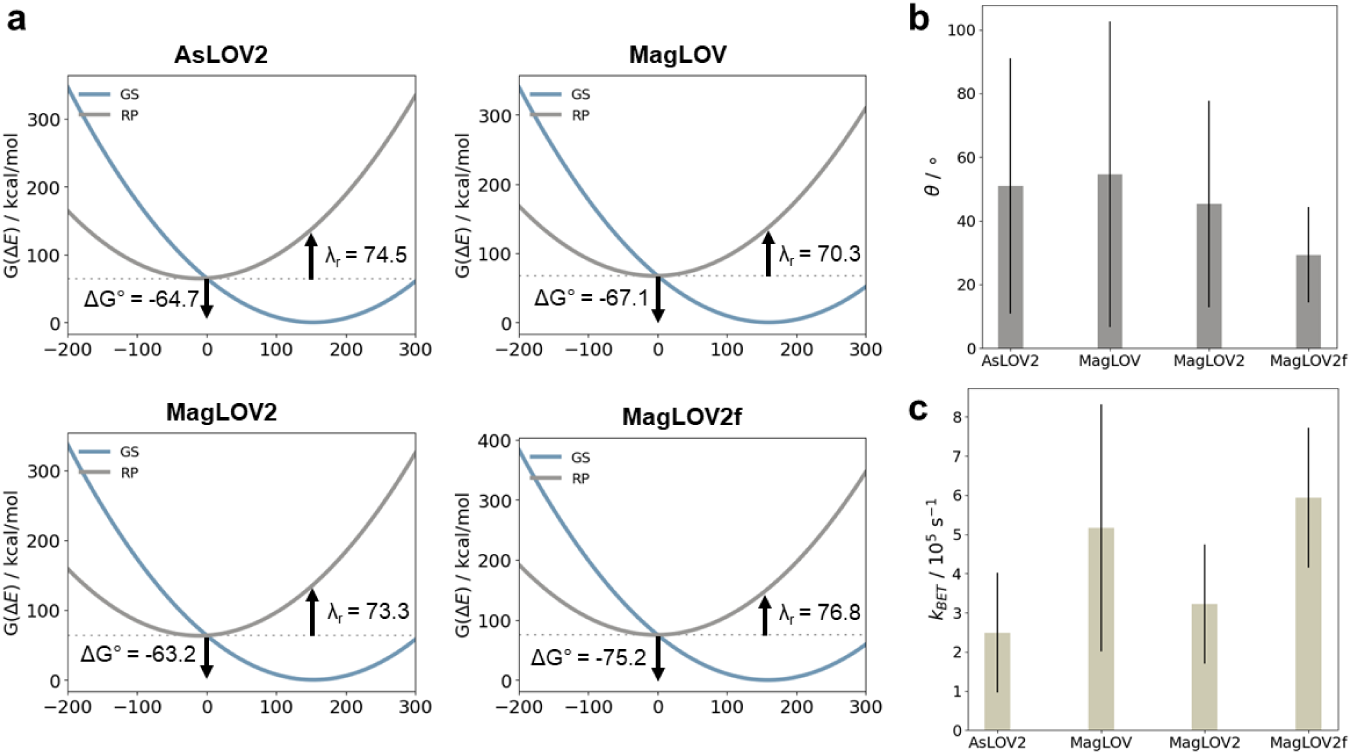
Free-energy, orientation, and back-electron-transfer differences among the four variants. **a** Marcus-type diabatic free-energy parabolas for RP*→*GS recombination in *As*LOV2, MagLOV, MagLOV2, and MagLOV2f. The indicated Δ*G^◦^* and *λr* values are in kcal mol*^−^*^1^. **b** Angle *θ* between the FMN and W89 aromatic-plane normals; bars show the ensemble mean and error bars the trajectory-derived spread. **c** Ensemble-averaged BET rate constants from the geometry-modulated Marcus model. Error bars report the sampled rate spread. Full energy-gap distributions and model definitions are given in Section S9 of the SI.

The structural data support this interpretation. The mean FMN–W89 COM distance is nearly unchanged across variants (Section S7 of the SI), whereas the aromatic-plane orientation changes substantially (Fig. 5b and Section S4 of the SI). The parental construct and MagLOV sample broad angular distributions with mean angles near 50–55*^◦^*, MagLOV2 shifts modestly toward smaller angles, and MagLOV2f shows the clearest enrichment of lower-angle geometries, with a mean near 30*^◦^* and a narrower distribution. Within the empirical overlap model used here, these lower angles increase the electronic-coupling factor without requiring a shorter mean COM separation. This is consistent with the W89 torsional analysis: MagLOV2f sacrifices rotameric heterogeneity while retaining broad rigid-body libration, thereby concentrating the donor in a more restricted internal geometry but allowing substantial orientational motion of the pair as a whole.

Combining the free-energy and geometric terms yields ensemble-averaged BET rates of approximately 2.5, 5.2, 3.2, and 5.9 × 10^5^ s*^−^*^1^ for *As*LOV2, MagLOV, MagLOV2, and MagLOV2f, respectively (Fig. 5c). The rates are therefore not monotonic with the number of mutations. MagLOV and MagLOV2f recombine most rapidly in this model, while MagLOV2 is intermediate and the parental construct is slowest. The comparatively fast MagLOV2f rate is notable because this variant was experimentally selected for a faster MFE saturation response rather than for increased MFE amplitude [5]. The calculated increase in BET provides a molecularly plausible contribution to this phenotype, but it should not be interpreted as a direct one-to-one prediction of the experimental saturation constant: the measured response also contains donor/acceptor recovery, spin mixing, and other photocycle steps [5].

The absolute BET values depend on the linear-response approximation and on the empirical Moser–Dutton/orientation model used for the electronic coupling (Section S9 of the SI). Accordingly, the most robust conclusion is the comparative one: evolution changes the recombination window through a combination of redox-state reorganisation and donor orientation, even though the average FMN–W89 separation remains nearly constant. This complements the relaxation analysis and argues against a single structural descriptor controlling the magnetic phenotype.

The combined rate picture is summarised in Fig. 6 using relative log-scaled coordinates for *k*_HFC_, *k_D_*, and *k*_BET_. These ternary coordinates are a visual comparison of rate classes, not kinetic branching fractions or probabilities. *As*LOV2 lies in a regime in which BET is comparatively prominent because both calculated relaxation contributions are small. MagLOV shifts strongly toward HFC modulation, while MagLOV2 combines the largest HFC contribution with a pronounced increase in dipolar modulation and an intermediate BET rate. MagLOV2f retains the large dipolar contribution of MagLOV2 but shifts away from HFC modulation and toward faster BET. The four constructs therefore do not form a simple scalar progression; each mutation set redistributes the processes that terminate or decohere the SCRP in a distinct way.

**Fig. 6.**
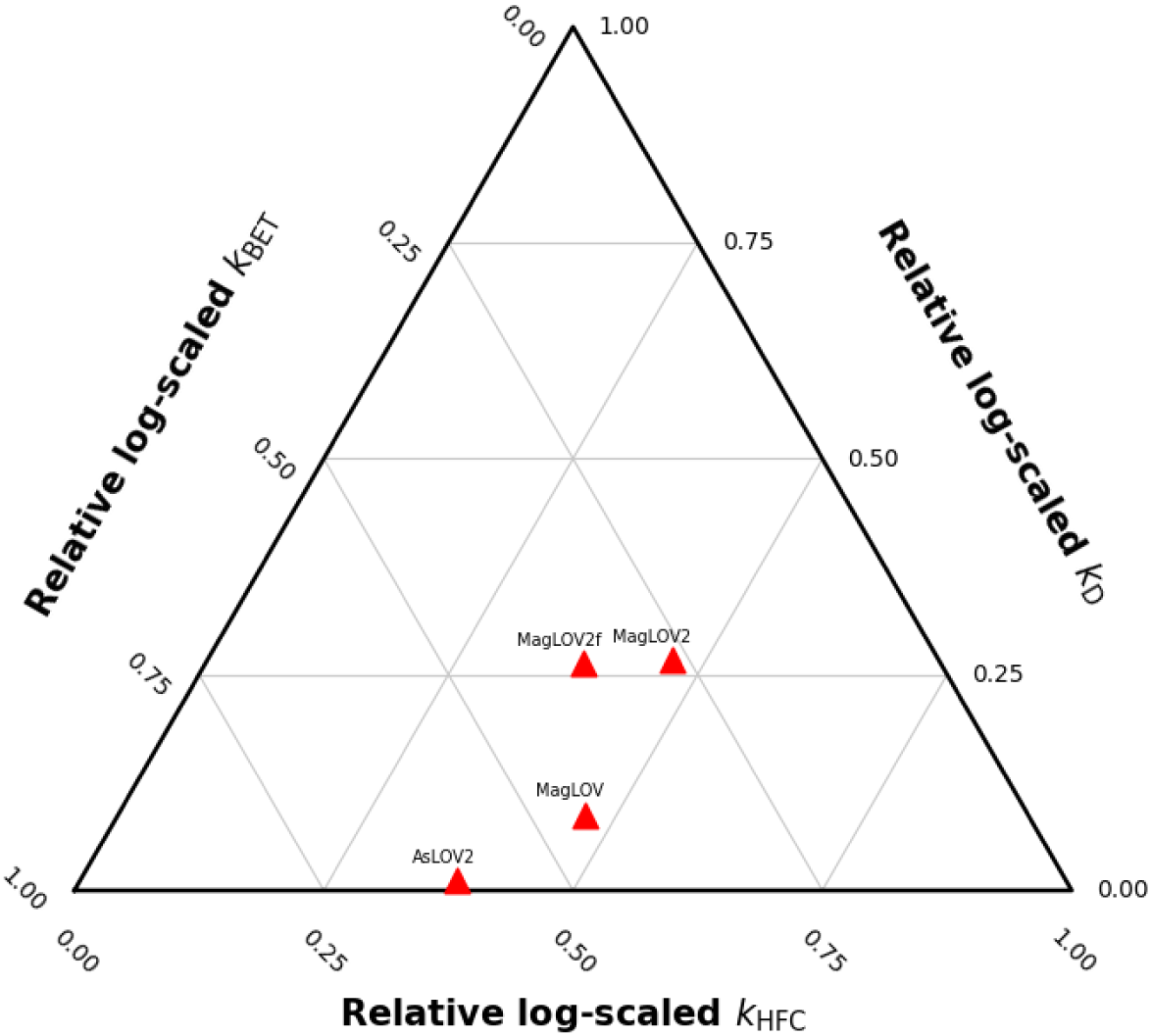
Relative balance of the three calculated mutation-sensitive rate classes. Ternary coordinates represent the normalised relative log-scaled contributions of *k*_HFC_, *k_D_*, and *k*_BET_ for each variant. The plotted coordinates are *As*LOV2 (0.3768, 0.0121, 0.6110), MagLOV (0.4679, 0.0880, 0.4440), MagLOV2 (0.4661, 0.2668, 0.2672), and MagLOV2f (0.3782, 0.2632, 0.3586), listed as (*k*_HFC_, *k_D_*, *k*_BET_). These coordinates are intended only to visualise relative kinetic balance and must not be interpreted as branching fractions.

## 3 Conclusion

The four *AsLOV2-derived* proteins reveal that directed evolution changes radical pair quantum sensing through the redistribution of several coupled dynamical and kinetic processes rather than through a monotonic increase in “stability” or “coherence”. Across the entire series, the LOV fold and the FMN-binding core remain intact, the mean FMN–W89 separation changes little, and FMN remains the comparatively rigid radical pair partner. The dominant structural response instead occurs at W89 and, crucially, separates into two different coordinates: rigid-body libration becomes broader in the later variants, whereas internal W89 torsional heterogeneity is progressively reduced. This resolves an apparent contradiction in the structural data and explains why dipolar- and hyperfine-driven relaxation need not change in parallel.

The individual variants occupy distinct mechanistic regimes. In *As*LOV2, W89 libration is comparatively restricted, and both dipolar- and HFC-modulation coefficients are small; the calculated BET rate is also the lowest of the four systems, giving the SCRP the broadest kinetic window for spin evolution in the present model. MagLOV increases HFC modulation substantially, while dipolar modulation remains relatively modest, and its calculated BET rate rises to approximately 5.2 × 10^5^ s*^−^*^1^. MagLOV2 represents a different balance: W89 rigid-body motion and slow dipolar correlations become much more prominent, HFC modulation reaches its largest value, and BET remains intermediate. This combination is consistent with selection having moved the protein into a strongly interaction-modulated regime rather than merely extending the radical pair lifetime.

MagLOV2f is mechanistically distinct from MagLOV2 despite their close sequence relationship. It retains the broad W89 librational landscape and large dipolar modulation contribution, but its W89 internal torsion is concentrated into a dominant rotameric basin, and its aromatic plane orientation shifts toward lower angles. Concomitantly, the calculated HFC modulation contribution decreases relative to MagLOV2, while BET becomes the fastest of the four variants (approximately 5.9×10^5^ s*^−^*^1^). This redistribution offers a specific molecular explanation for why the “fast” variant can exhibit different response dynamics: the mutations do not simply accelerate all decoherence processes, but combine faster recombination with partial suppression of the HFC random field while preserving strong dipolar modulation. The experimentally observed increase in MFE saturation rate of MagLOV2f [5] is therefore qualitatively consistent with a shortened recombination window, although a quantitative mapping requires full spin-kinetic simulations of the complete photocycle.

These results suggest a practical design principle for future protein radical pair quantum sensors. Engineering should target the *distribution* of donor conformations rather than generic protein rigidity: internal rotameric locking can reduce one source of magnetic interaction noise even when larger scale donor libration remains substantial, while local electrostatics and donor–acceptor orientation can independently tune recombination. The relevant optimisation objective is therefore a controlled balance among spin mixing, dipolar and HFC relaxation, and spin-selective reaction kinetics. By identifying how the same protein scaffold can access markedly different balances of these processes, the present work provides a molecular framework for rationally separating sensor amplitude, response speed, and environmental sensitivity in genetically encoded radical pair quantum probes.

## 4 Methods

### 4.1 Molecular dynamics simulations

Three-dimensional structures of the individual *As*LOV2-derived variants were generated using AlphaFold [40]. Molecular dynamics (MD) simulations were performed with NAMD [41, 42] using the CHARMM36 protein force field [43, 44] together with previously established FMN parameters [45]. The proteins were solvated explicitly with TIP3P water [46], with at least 20 5 of solvent between the protein and each simulation-box boundary, and NaCl was added to a concentration of 50 mM.

Periodic boundary conditions were applied throughout. Long-range electrostatic interactions were evaluated with the particle mesh Ewald method [47], and van der Waals interactions were smoothly switched off between 10 and 12 5. Bonds involving hydrogen atoms were constrained with SHAKE [48]. The temperature was maintained at 310 K with a Langevin thermostat [42]; during constant-pressure equilibration, the pressure was maintained at 1 atm using the Langevin-piston Nosé–Hoover method [49].

Each system was first subjected to 20,000 minimisation steps. Harmonic positional restraints were then gradually released during staged equilibration to relax the solvent and protein environment while avoiding abrupt distortion of the initial structure. Production trajectories were propagated in the NVT ensemble at 310 K as three independent 500 ns replicas for each electronic state and variant. The complete equilibration and production protocol, including system sizes, is given in Section S1 of the Supporting Information (SI). Trajectory visualisation and structural analyses were performed with VMD [50].

### 4.2 Quantum chemical calculations of energies

Thirty representative snapshots were extracted for the quantum-chemical correction used in the energy-gap analysis. For each snapshot, the redox-active fragments were truncated as required, and dangling valences were saturated with hydrogen atoms. Heavy-atom coordinates were retained from the MD structures to preserve the sampled conformations, whereas hydrogen positions, including capping hydrogens, were optimised before energy evaluation.

All quantum-chemical calculations were performed with ORCA [51, 52]. Hydrogen-only geometry refinements employed Kohn–Sham DFT with B3LYP [53, 54], D3 dispersion with Becke–Johnson damping [55, 56], and the def2-TZVP basis set [57]. Single-point energies were subsequently evaluated with CAM-B3LYP [58] and the 6-31G basis set [59]. The same two-step protocol was applied to every sampled structure. Because the fragment-level quantum correction is used within a comparative energy-gap model rather than as a stand-alone prediction of absolute redox thermochemistry, the resulting absolute free energies and rates should be interpreted as model-dependent; the relative changes among variants are the primary quantities considered here.

### 4.3 Back electron transfer rate constants

Back electron transfer (BET) from the charge-separated radical pair state (RP) to the electronic ground state (GS) was treated in the nonadiabatic limit with semiclassical Marcus theory [60, 61]. Vertical RP–GS energy gaps were evaluated on both the GS and RP MD ensembles and converted, under a Gaussian linear-response approximation, into diabatic free-energy parabolas [62, 63]. A subtractive QM/MM-style correction was used to replace the force-field description of the two redox-active fragments with state-specific quantum chemical fragment energies, following the general logic of multi-layer energy corrections [64, 65]. The Marcus prefactor was combined with an electronic-coupling model calibrated to the Moser–Dutton protein electron-transfer ruler [66]. The Moser–Dutton distance term sets the baseline coupling from the ensemble-mean edge-to-edge separation, while the sampled donor– acceptor orientation enters through an empirical |cos *α*| modulation of the aromatic-plane overlap [67, 68]; the same distance attenuation is not applied a second time at the frame level. The complete definitions, assumptions, and uncertainty-relevant limitations of this construction are given in Section S9 of the SI.

### 4.4 Hyperfine coupling calculations

To determine how conformational dynamics modulate the local magnetic interactions of the radical pair, hyperfine coupling (HFC) tensors were evaluated along the RP-state trajectories for FMN*^•−^* and W89H*^•^*^+^. The three independent 500 ns replicas of each protein variant were analysed separately. For each replica, 5000 structures were sampled at 100 ps intervals, resulting in 120,000 fragment HFC calculations and a total of 4,620,001 HFCs across the four protein variants and two radical fragments. The fragments were extracted with identical atom ordering throughout all trajectories. Heavy-atom coordinates were retained from the MD frames, whereas hydrogen positions and terminal capping atoms were relaxed before the HFC calculations.

The constrained optimisations used unrestricted B3LYP with D4 dispersion [69], def2-TZVP [57], RIJCOSX, and TightSCF convergence. HFC tensors were then calculated with B3LYP-D4 and the pcH-2 basis set, which was developed specifically for hyperfine-coupling calculations [70], again using TightSCF convergence in ORCA [51, 52]. The same electronic-structure protocol was applied to every sampled structure and variant. Trajectory-resolved quantum chemical HFC calculations have previously shown that thermal distortions and the protein environment can substantially modulate flavin hyperfine interactions [71, 72]. The globally selected nuclei, complete trajectory-resolved isotropic HFCs, and full-tensor fluctuation amplitudes are reported in Section S8 of the SI.

### 4.5 Spin relaxation rate constants

#### Dipolar modulation

The electron–electron dipolar interaction was analysed in angular-frequency units,

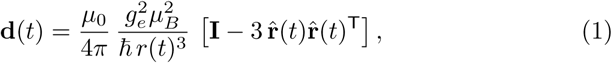

where **r**^(*t*) = **r**(*t*)*/r*(*t*). For any scalar time series *K*(*t*), including an individual tensor element *d_αβ_*(*t*), fluctuations were defined as *δK*(*t*) = *K*(*t*) − ⟨*K*⟩ and the autocovariance as

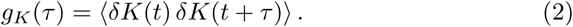

The corresponding integral correlation time is

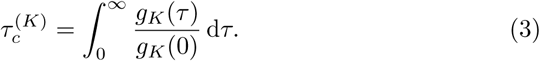

Because the MD correlation functions are generally non-exponential, their normalised forms are represented by non-negative multi-exponential fits [39, 73, 74],

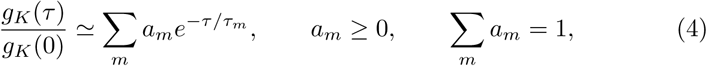

so that 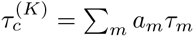. Following the zero-frequency stochastic-modulation treatment used for radical pairs [38], we define the diagonal effective dipolar-modulation coefficients

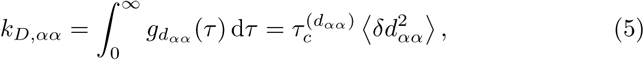

which have units of s*^−^*^1^. These coefficients quantify the relative strength of dipolar random-field modulation among variants; they are not identified with a unique field-dependent 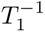 or 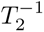 without a full spectral-density treatment. Exchange modulation was evaluated analogously using the distance-dependent model specified in Section S6 of the SI and was found to be much smaller than the dipolar contribution. Dipolar correlation functions, replica-specific correlation times, and FMN–W89 distance distributions are provided in Sections S7–S8 of the SI.

#### Hyperfine modulation

The time-dependent HFC Hamiltonian for nucleus *n* is

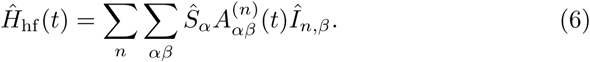

For the relaxation analysis, HFC tensors reported in MHz were converted to angular-frequency units according to

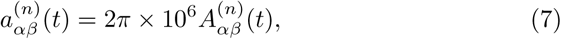

with *a* in rad s*^−^*^1^, only their fluctuating components 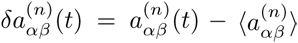 were used. For an unpolarised nuclear spin *I_n_*, the effective Cartesian random-field covariance was evaluated as [71, 75]

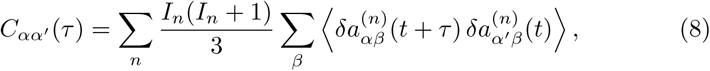

so that the nuclear-spin factors are 1*/*4 for ^1^H and 2*/*3 for ^14^N. The diagonal covariance functions were treated with the same non-negative multi-exponential procedure, and the effective random-field coefficients were

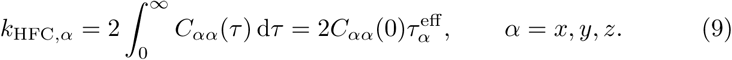

Individual nuclear contributions were analysed separately and summed only after their covariance functions had been evaluated. For compact comparisons, we additionally use *k*_HFC_ = (*k*_HFC_*_,x_* + *k*_HFC_*_,y_* + *k*_HFC_*_,z_*)*/*3. The same globally selected H and N nuclei were retained for every variant, preventing variant-dependent HFC selection from generating artificial rate differences. Correlation-function processing and relaxation analysis were implemented within the *RadicalPy*-based workflow [2]; further details and the complete HFC time series are given in Section S8 of the SI.

## Supplementary information

The Supporting Information contains: (i) the complete MD equilibration and production protocol; (ii) replica-resolved RMSD and RMSF analyses, including the extended MagLOV2f trajectories; (iii) total-protein, FMN, and W89 solvent-accessibility distributions; (iv) spherical molecular-axis plots, axis-resolved librational distributions, FMN/W89 dihedral distributions, and FMN–W89 aromatic-plane-angle distributions; (v) FMN and W89 hydrogen-bond analyses; (vi) the exchange-coupling parameterisation and exchange-modulation assessment; (vii) FMN– W89 centre-of-mass distance distributions; (viii) dipolar autocorrelation functions, HFC nuclear labelling, and the complete isotropic and full-tensor HFC fluctuation trajectories used in the relaxation analysis; and (ix) the complete energy-gap, free-energy, and BET-rate model.

## Acknowledgments

The authors thank H. Steel, C. R. Timmel, G. Abrahams, and A. S^̌^tuhec for fruitful discussions.

## Author information

### Contributions

L.M.A. and L.G. designed and coordinated the study and wrote the manuscript. L.M.A. and L.G. performed simulations and data analysis. J.K.B. performed data analysis.

### Ethics declarations

The authors declare no competing interests.

