## Supplementary Information for "Revealing essential properties for enhanced quantum sensing in engineered *As*LOV2 proteins"

### Supplementary Information: Revealing essential properties for enhanced quantum sensing in engineered AsLOV2 proteins

### Contents

**S1 Molecular dynamics** **S3**

**S2 Protein stability** **S4**

**S3 Surface accessibility** **S7**

**S4 Molecular fluctuations** **S10**

**S5 Hydrogen bonding** **S22**

**S6 Exchange interaction** **S26**

**S7 Centre-to-centre distances** **S27**

**S8 Spin relaxation** **S29**

S8.1 Dipolar coupling modulation . . . . . . . . . . . . . . . . . . . . . . . . . . . . S29

S8.2 Hyperfine coupling modulation . . . . . . . . . . . . . . . . . . . . . . . . . . . S30

S8.2.1 FMN hyperfine coupling modulation . . . . . . . . . . . . . . . . . . . S31

S8.2.2 Trp hyperfine coupling modulation . . . . . . . . . . . . . . . . . . . . S58

**S9 Back electron transfer rate constants** **S84**

### S1 Molecular dynamics

Table S1: Summary of the molecular dynamics simulations performed in this study. Three independent 500 ns production runs were generated for each state, corresponding to a total sampling time of 1.5 $\mu$s per state.

| State | Equilibration phase | | | | Production simulation (3 runs each) | System information | |
|---|---|---|---|---|---|---|---|
| | Stage 0 | Stage 1 | Stage 2 | Stage 3 | | Box size/Å | Number of atoms |
| | Minimization | Water and ions | Water, ions, side chains | All atoms | | | |
| *As*LOV2 DS | 20,000 steps | NPT, 1 ns | NPT, 2 ns | NVT, 2 ns | NVT, 500 ns | 92.08 × 104.89 × 100.31 | 57,611 |
| *As*LOV2 RP | 20,000 steps | NPT, 1 ns | NPT, 2 ns | NVT, 2 ns | NVT, 500 ns | 92.08 × 104.89 × 100.31 | 57,613 |
| MagLOV DS | 20,000 steps | NPT, 1 ns | NPT, 2 ns | NVT, 2 ns | NVT, 500 ns | 92.08 × 104.89 × 100.31 | 57,636 |
| MagLOV RP | 20,000 steps | NPT, 1 ns | NPT, 2 ns | NVT, 2 ns | NVT, 500 ns | 92.08 × 104.89 × 100.31 | 57,638 |
| MagLOV2 DS | 20,000 steps | NPT, 1 ns | NPT, 2 ns | NVT, 2 ns | NVT, 500 ns | 92.08 × 104.89 × 100.31 | 58,168 |
| MagLOV2 RP | 20,000 steps | NPT, 1 ns | NPT, 2 ns | NVT, 2 ns | NVT, 500 ns | 92.08 × 104.89 × 100.31 | 58,170 |
| MagLOV2f DS | 20,000 steps | NPT, 1 ns | NPT, 2 ns | NVT, 2 ns | NVT, 500 ns | 92.08 × 104.89 × 100.31 | 57,540 |
| MagLOV2f RP | 20,000 steps | NPT, 1 ns | NPT, 2 ns | NVT, 2 ns | NVT, 500 ns | 92.08 × 104.89 × 100.31 | 57,542 |

#### S2 Protein stability

The structural stability of each simulated state (Table S1) was assessed using the backbone root-mean-square deviation (RMSD) and the per-residue root-mean-square fluctuation (RMSF). Prior to both analyses, each trajectory frame was least-squares fitted to the reference structure to remove overall translation and rotation.

For a set of $n$ fitted atomic coordinates, the RMSD relative to the reference structure is

$$\mathrm{RMSD}_{\mathbf{x},\mathbf{x}^{\mathrm{ref}}} = \sqrt{\frac{1}{n}\sum_{i=1}^{n}\left|\mathbf{x}_i - \mathbf{x}_i^{\mathrm{ref}}\right|^2}, \tag{S1}$$

where $\mathbf{x}_i^{\mathrm{ref}}$ and $\mathbf{x}_i$ are the fitted positions of atom $i$ in the reference structure and at the analysed time point, respectively.

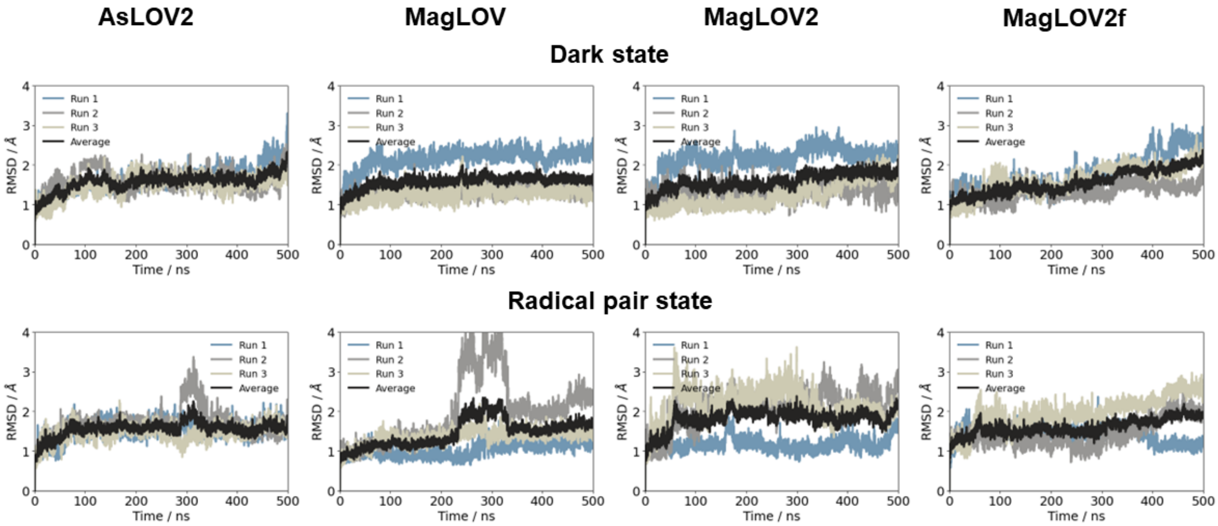


Figure S1: Time evolution of the backbone RMSD for each protein variant in the dark state and radical pair state during the three independent 500 ns production simulations. The black trace is the pointwise average over the three replicas.

Figure S1 shows that the LOV2 fold remains stable in all variants and electronic states. Most trajectories rapidly reach plateaus in the approximate range of 1–2.5 Å, while several replicas undergo larger but bounded excursions. The most pronounced event occurs in the

MagLOV radical pair state, where one replica enters a distinct conformational basin after approximately 250 ns and subsequently stabilises at a higher RMSD. The absence of a sustained, unbounded increase across the ensemble argues against global unfolding. Instead, the RMSD changes reflect replica-specific transitions between metastable conformations, primarily involving flexible surface and terminal regions.

The MagLOV2f dark-state trajectory exhibits a gradual increase during the first 500 ns. The extended trajectory in Fig. S2 shows that this increase corresponds to a transition to a second plateau rather than continued structural drift. The radical-pair-state trajectory likewise remains bounded over 900 ns. This behaviour supports the interpretation that the elevated RMSD results from a persistent conformational rearrangement within an intact fold.

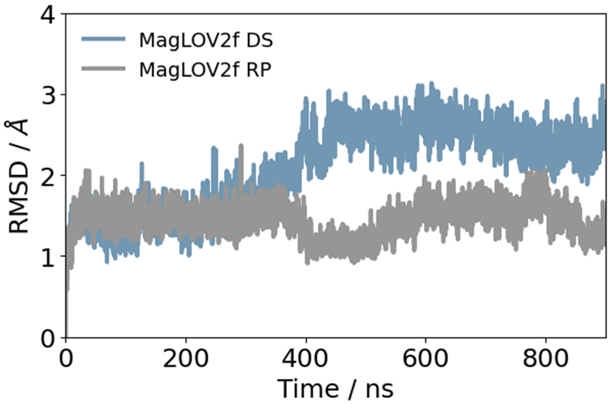


Figure S2: Backbone RMSD of MagLOV2f in the dark state and radical pair state for production simulations extended to 900 ns. Both trajectories remain bounded after the conformational changes observed during the first 500 ns.

The RMSF quantifies the positional variance of an atom or residue $i$ about its time-averaged position,

$$\mathrm{RMSF}_i = \sqrt{\left\langle \left| \mathbf{x}_i - \langle \mathbf{x}_i \rangle \right|^2 \right\rangle}, \tag{S2}$$

where $\langle \cdots \rangle$ denotes an average over the trajectory. Whereas the RMSD reports the global displacement from a reference structure, the RMSF identifies local regions that repeatedly depart from their mean positions.

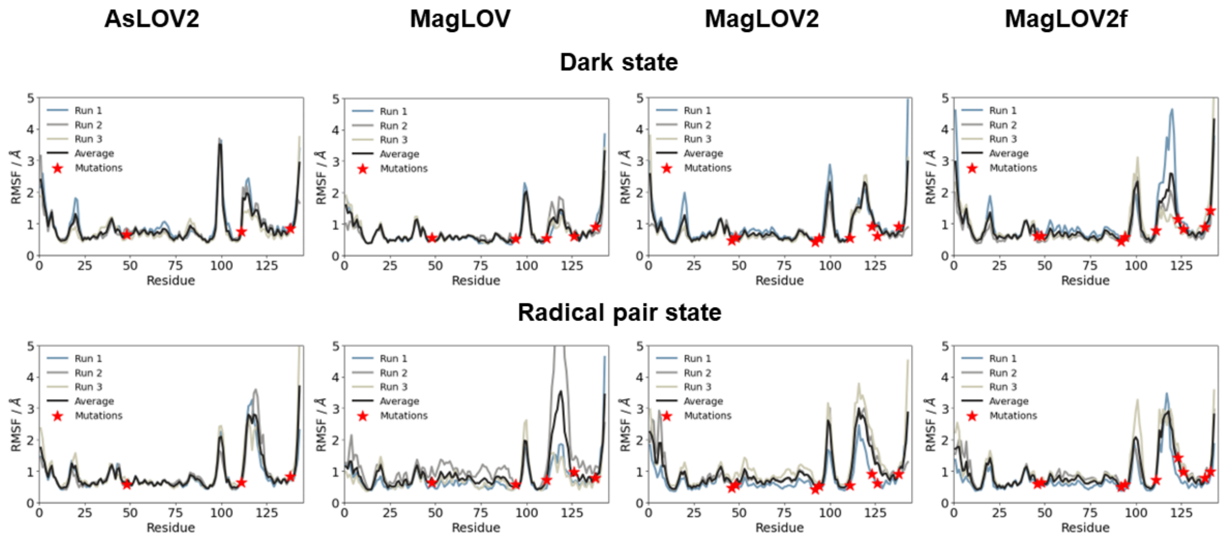


Figure S3: Per-residue RMSF for each protein variant in the dark state and radical pair state. The coloured traces represent the three independent replicas, the black trace is their average, and red stars mark the positions of the introduced mutations.

The low RMSF baseline across the structured core confirms that the mutations do not destabilise the canonical LOV2 architecture (Fig. S3). The largest fluctuations are localised to the N-terminal segment, short solvent-exposed loops, the region around residues 95–105, and the J$\alpha$/C-terminal region around residues 115–145. The amplitudes of these peaks vary between replicas, most notably for the MagLOV radical pair state, which is consistent with the conformational transition observed in the RMSD. The mutation sites do not coincide systematically with global maxima; rather, the mutations redistribute mobility locally and through longer-range coupling to flexible loops and terminal elements.

#### S3 Surface accessibility

Solvent-accessible surface areas (SASA) were evaluated for the complete protein and separately for FMN and W89. The total SASA reflects global expansion or compaction, whereas the fragment-resolved SASA is sensitive to local pocket breathing and hydration.

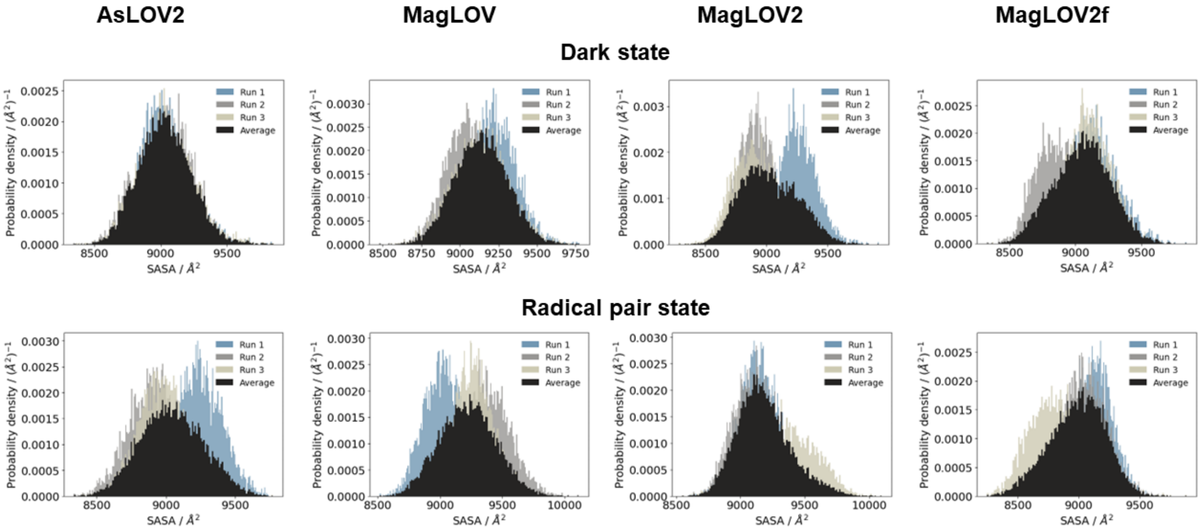


Figure S4: Probability-density distributions of the total protein SASA for each variant in the dark state and radical pair state. The coloured distributions correspond to the three replicas and the black distribution represents their average.

The total protein SASA is comparable across all variants and states (Fig. S4); the reported means span approximately 8900–9200 Å$^2$. The distributions overlap strongly despite replica-specific shifts and widths. Together with the RMSD and RMSF analyses, this confirms that neither radical-pair formation nor the introduced mutations cause global unfolding or persistent exposure of a substantially larger protein surface.

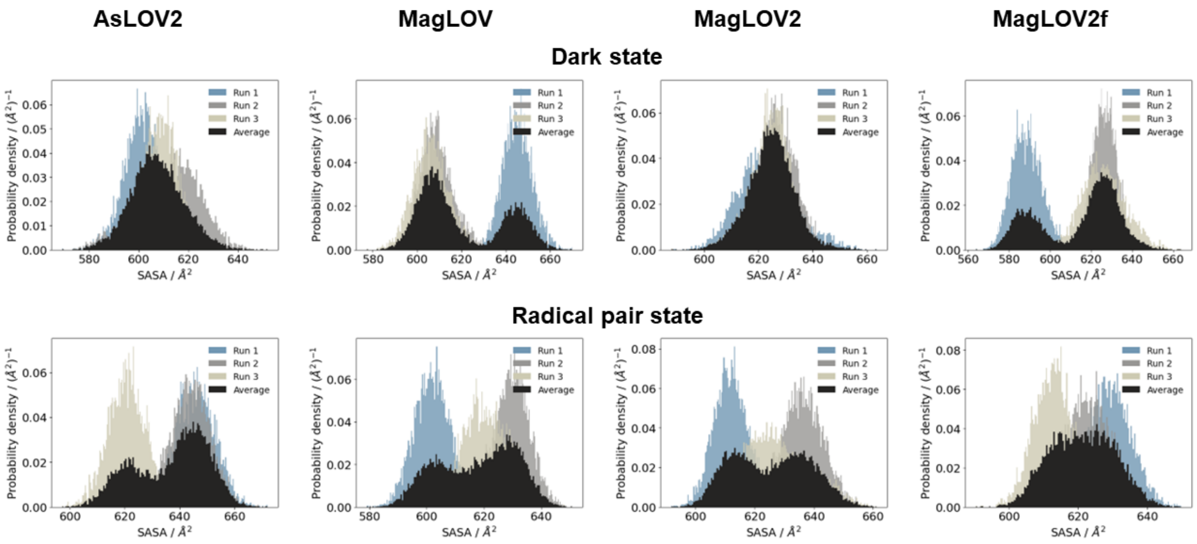


Figure S5: Probability-density distributions of the FMN SASA for each variant in the dark state and radical pair state. The coloured distributions correspond to the three replicas and the black distribution represents their average.

The FMN SASA exhibits stronger replica and state dependence than the total protein SASA (Fig. S5). *As*LOV2 and MagLOV2 develop clearly bimodal averaged distributions in the radical pair state because individual replicas occupy pocket conformations with different FMN exposure. MagLOV and MagLOV2f are already bimodal in the dark state and become more strongly overlapping in the radical pair state. These changes predominantly affect the populations of pre-existing exposure states rather than the overall range of FMN SASA. The cofactor therefore remains embedded in the binding pocket, while local hydration and pocket breathing are redistributed by the charge state and the mutation pattern.

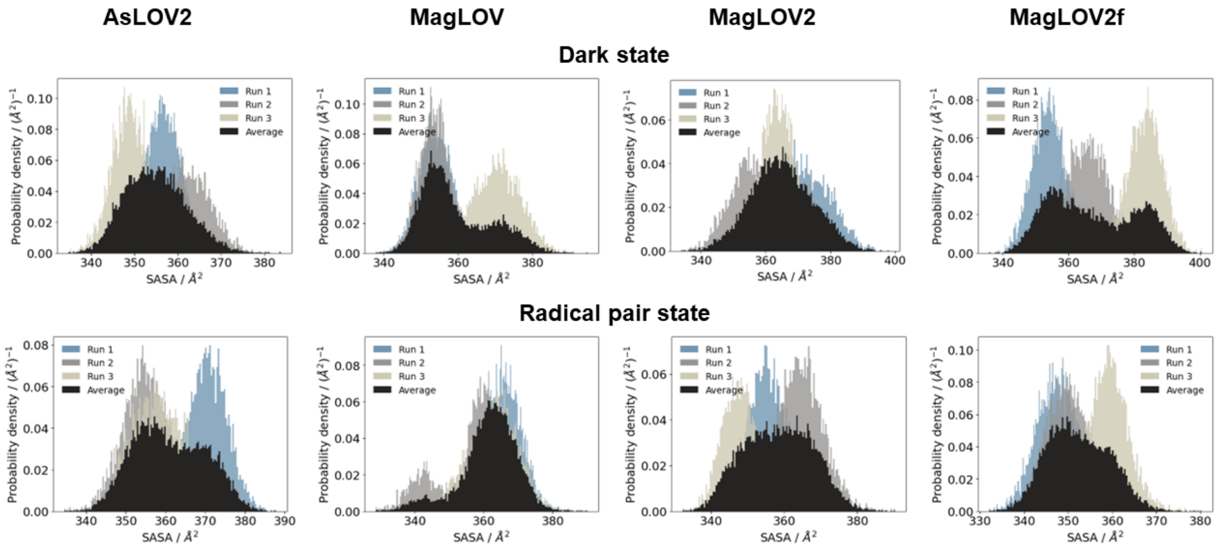


Figure S6: Probability-density distributions of the W89 SASA for each variant in the dark state and radical pair state. The coloured distributions correspond to the three replicas and the black distribution represents their average.

The W89 distributions also contain replica-specific and, in some cases, bimodal populations (Fig. S6), but no systematic increase in solvent exposure accompanies either radical-pair formation or progressive mutation. *As*LOV2 and MagLOV2 show broadly similar dark-state distributions, whereas MagLOV and MagLOV2f display a more evident two-state character. In the radical pair state, the distributions remain within a comparable SASA window for all variants. Consequently, the altered donor dynamics discussed below cannot be reduced to a simple monotonic change in W89 solvent exposure. Changes in packing, electrostatics, and access to distinct orientational substates are also required.

### S4 Molecular fluctuations

The local motion of the radical-pair partners was decomposed into rigid-body libration of three molecular axes and internal torsion about a representative dihedral angle. The spherical plots display the directions sampled by the molecular axes, while the corresponding one-dimensional distributions resolve the angular amplitudes for each replica. This separation is important because broad rigid-body reorientation and broad internal torsional sampling represent distinct physical modes and need not follow the same mutation trend.

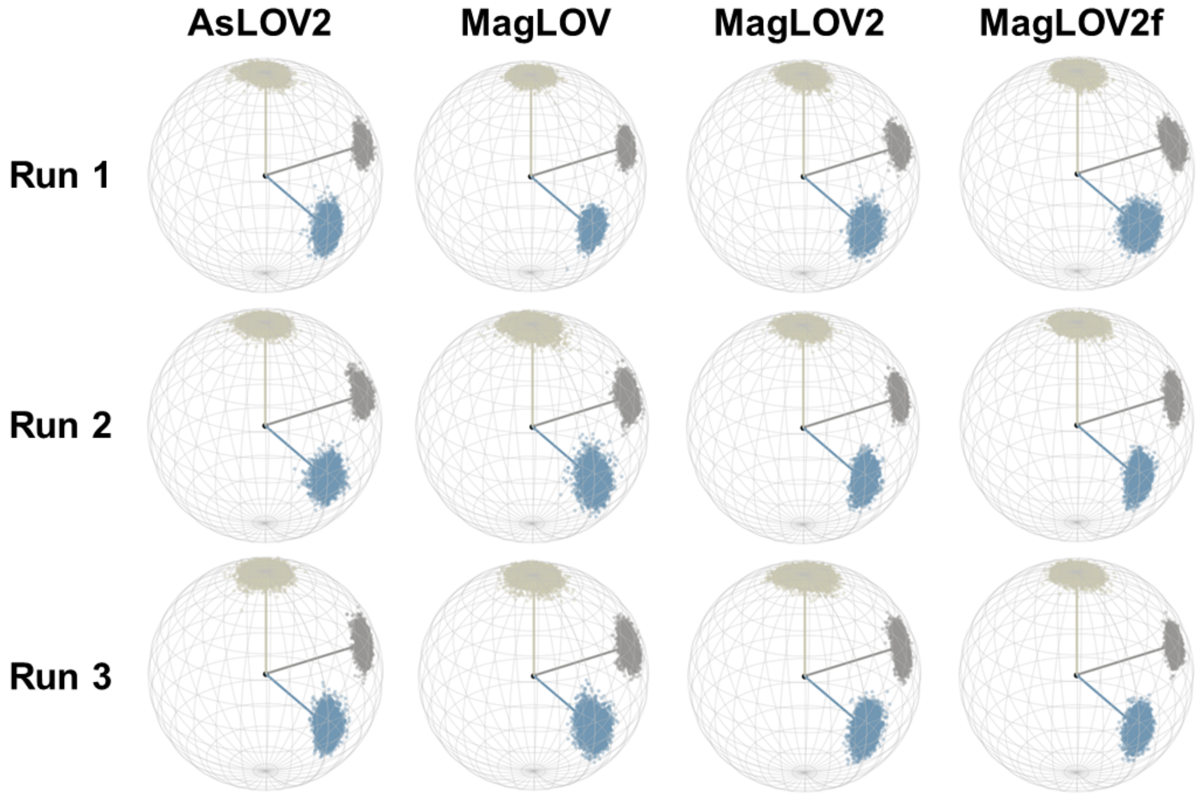


Figure S7: Spherical scatter plots of the $FMN^{\bullet-}$ molecular-axis directions sampled during each 500 ns radical-pair-state trajectory. Columns correspond to the protein variants and rows to the three independent replicas.

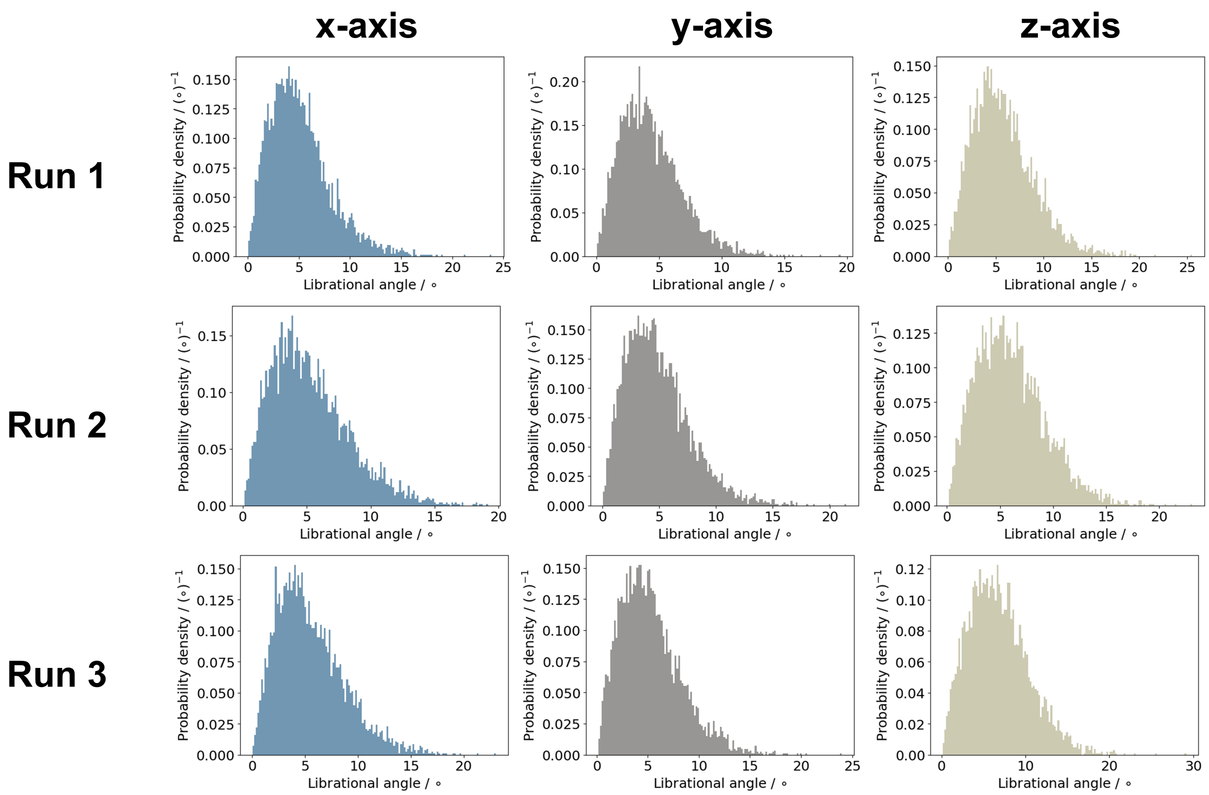


Figure S8: Axis-resolved probability-density distributions of FMN$^{\bullet -}$ librational angles in *As*LOV2 for each 500 ns trajectory.

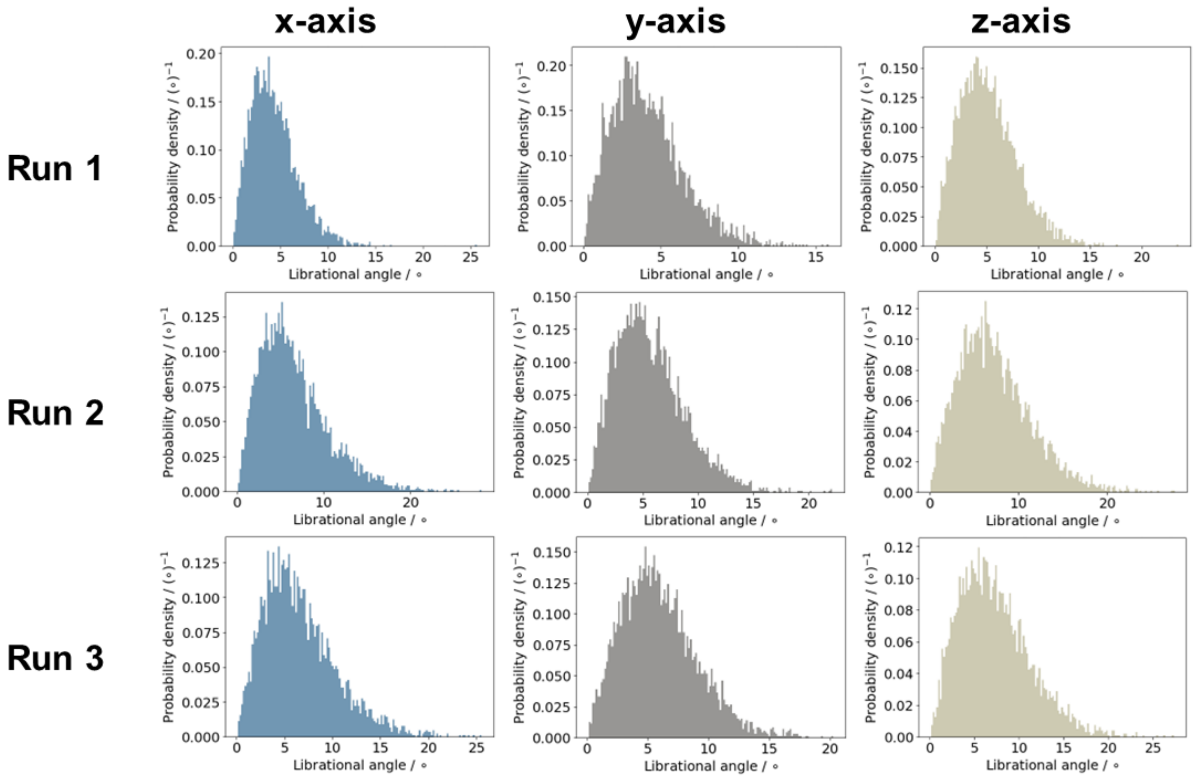


Figure S9: Axis-resolved probability-density distributions of $FMN^{\bullet-}$ librational angles in MagLOV for each 500 ns trajectory.

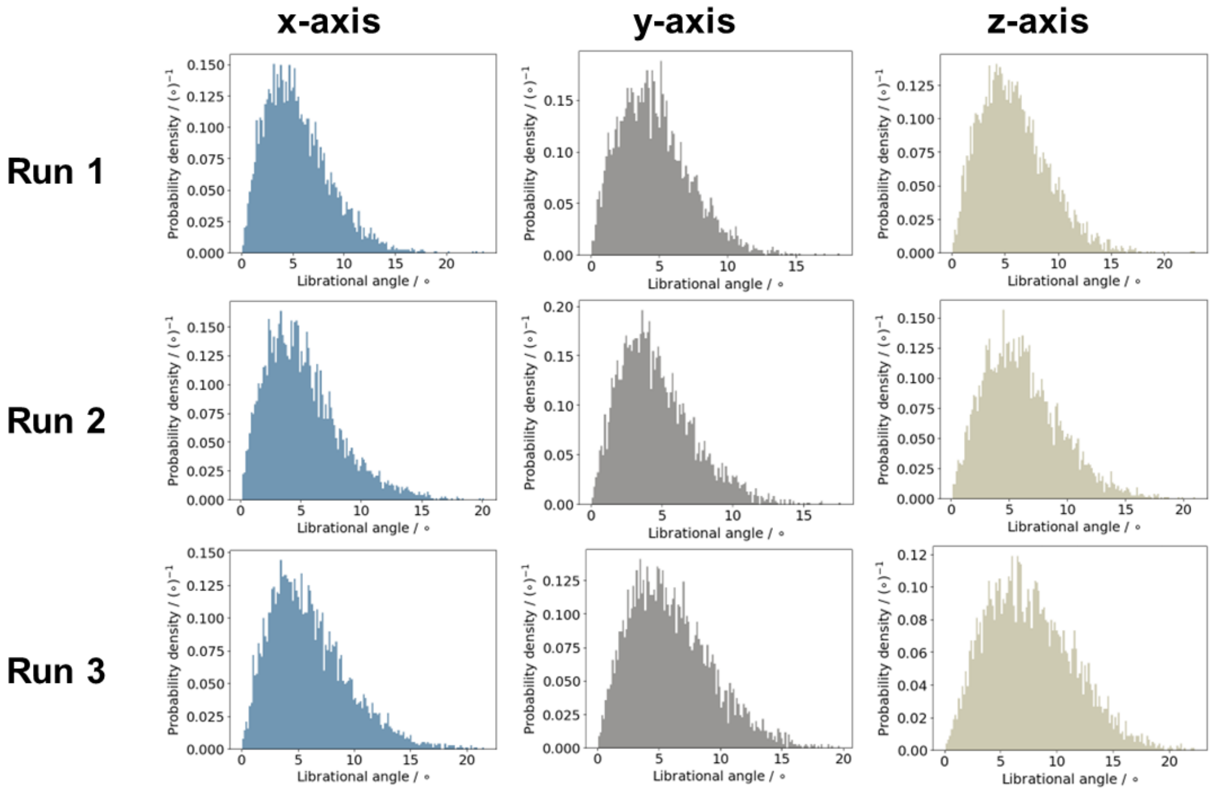


Figure S10: Axis-resolved probability-density distributions of FMN$^{\bullet-}$ librational angles in MagLOV2 for each 500 ns trajectory.

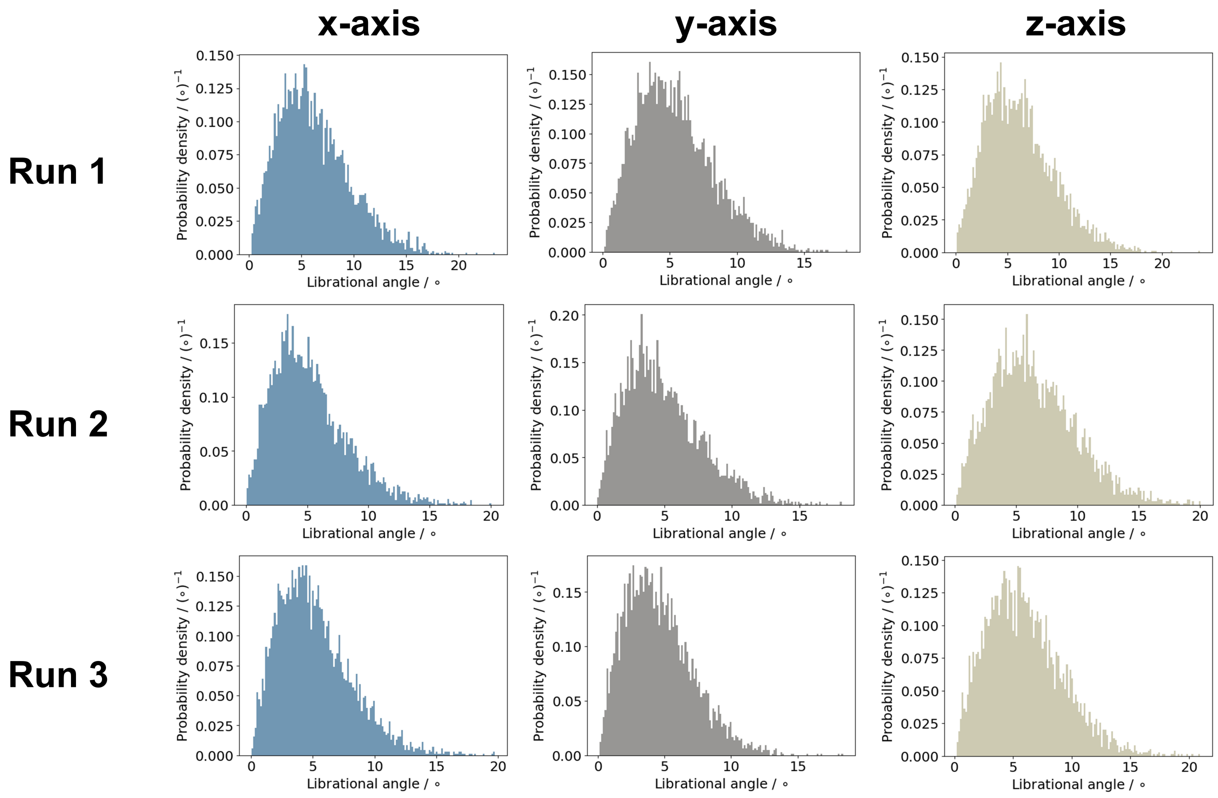


Figure S11: Axis-resolved probability-density distributions of $FMN^{\bullet-}$ librational angles in MagLOV2f for each 500 ns trajectory.

The FMN axis clouds are compact and occupy similar regions of the sphere in all variants (Fig. S7). Correspondingly, the axis-resolved distributions in Figs. S8–S11 are predominantly unimodal and concentrated at small angles, with only modest axis and replica dependence. No monotonic broadening is observed along the *As*LOV2–MagLOV–MagLOV2–MagLOV2f series. FMN therefore acts as a comparatively rigid structural anchor, consistent with the preservation of the cofactor pocket inferred from the RMSF, SASA, and hydrogen-bond analyses.

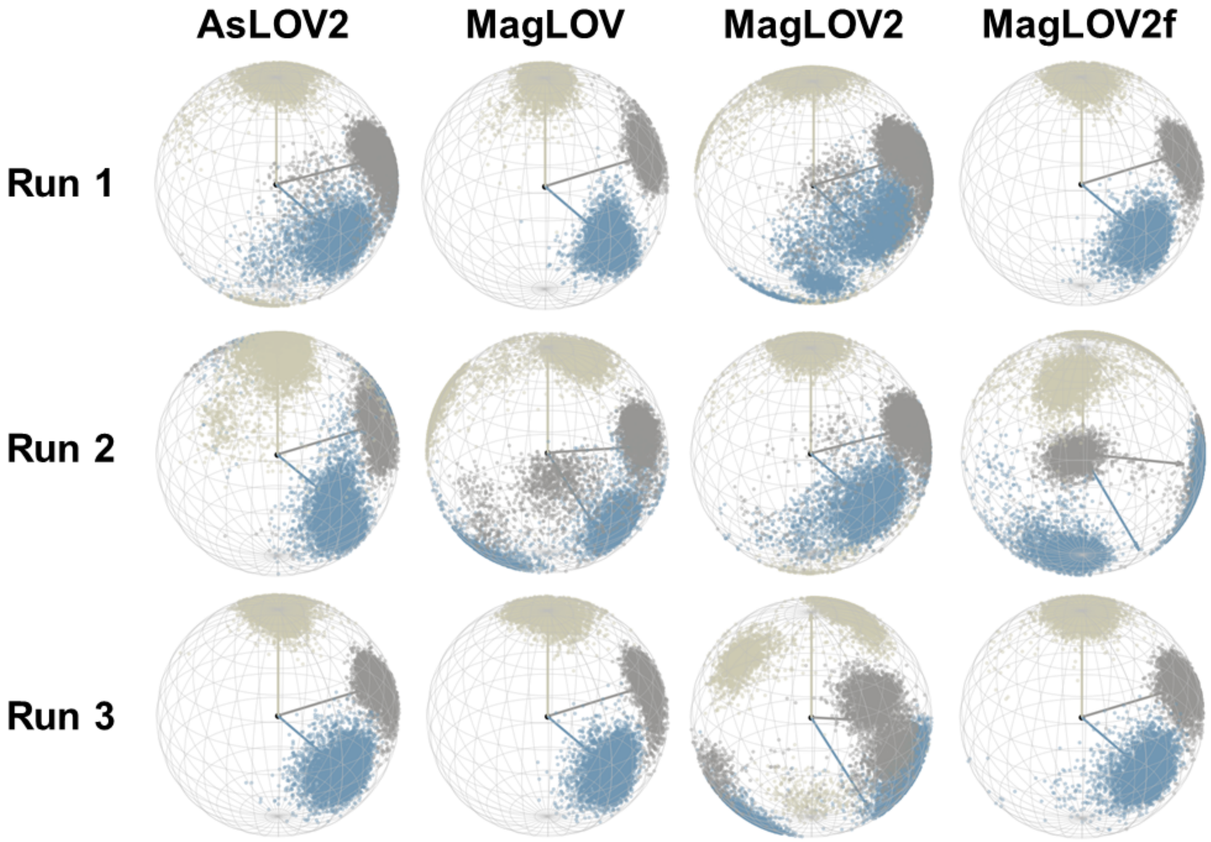


Figure S12: Spherical scatter plots of the W89H$^{\bullet+}$ molecular-axis directions sampled during each 500 ns radical-pair-state trajectory. Columns correspond to the protein variants and rows to the three independent replicas.

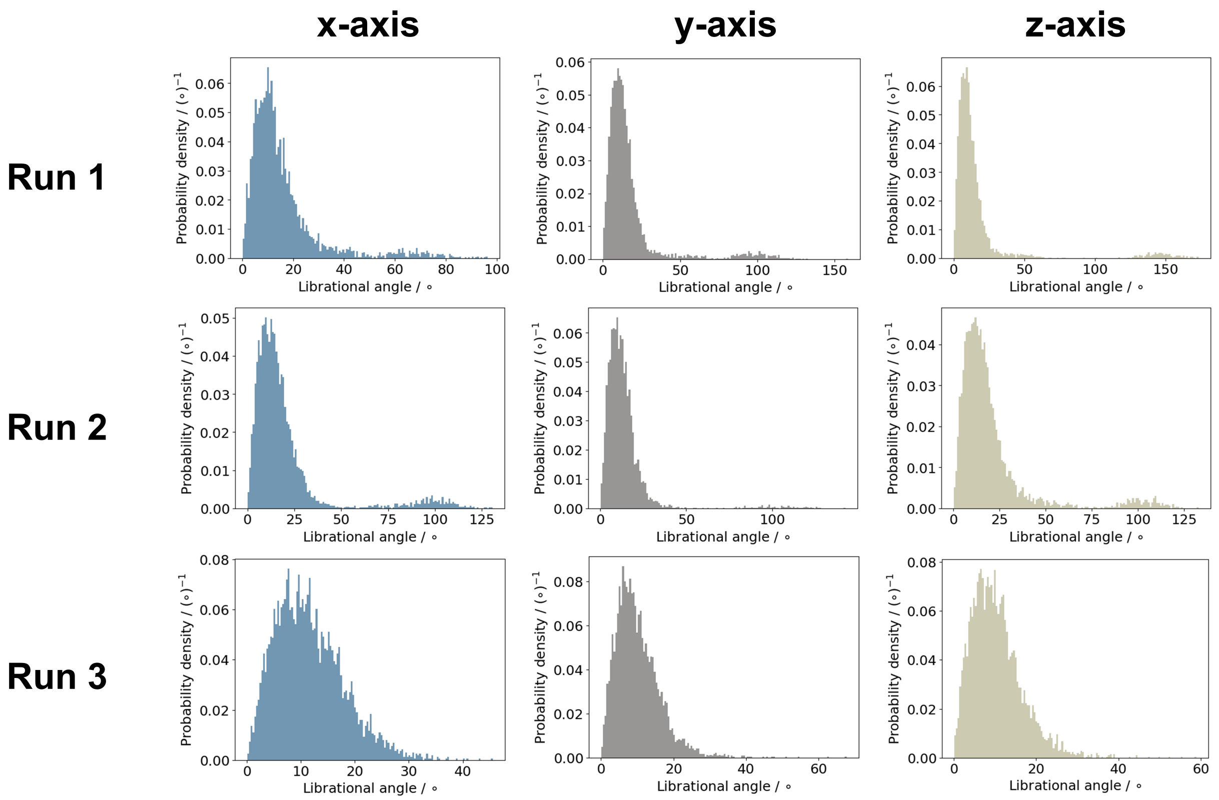


Figure S13: Axis-resolved probability-density distributions of W89H$^{\bullet+}$ librational angles in *As*LOV2 for each 500 ns trajectory.

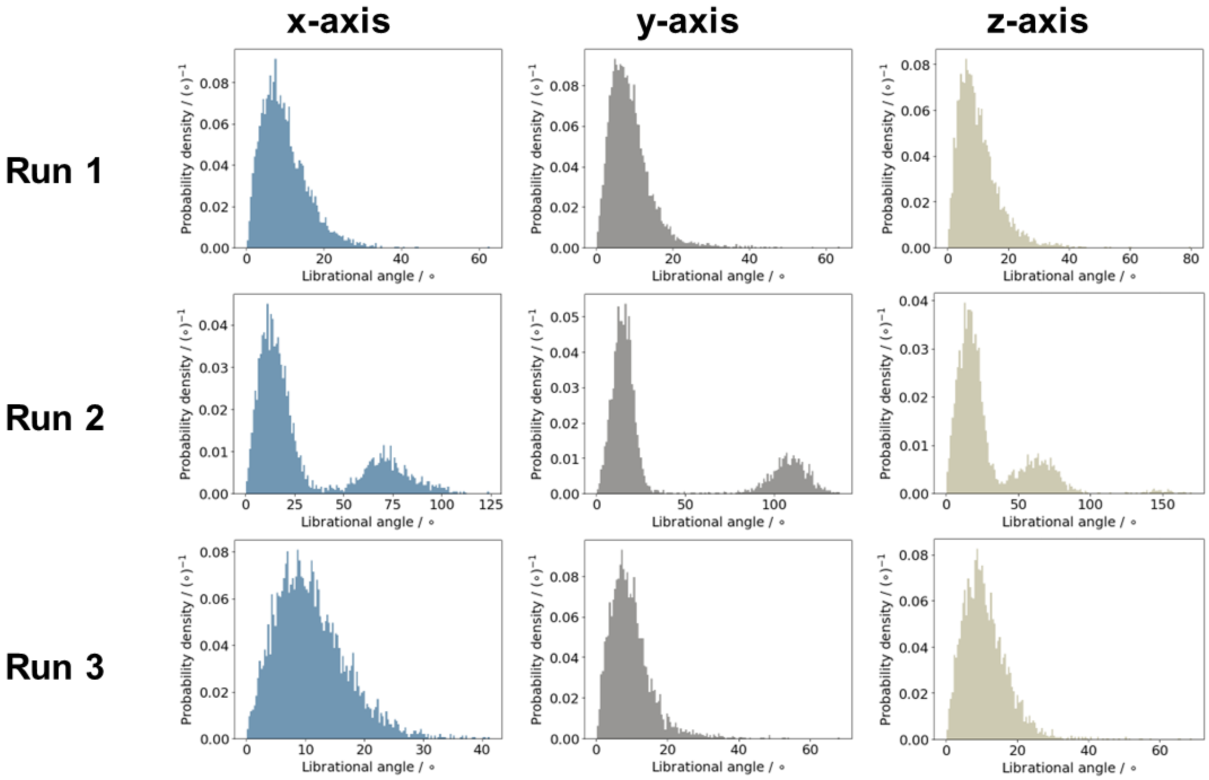


Figure S14: Axis-resolved probability-density distributions of W89H$^{\bullet+}$ librational angles in MagLOV for each 500 ns trajectory.

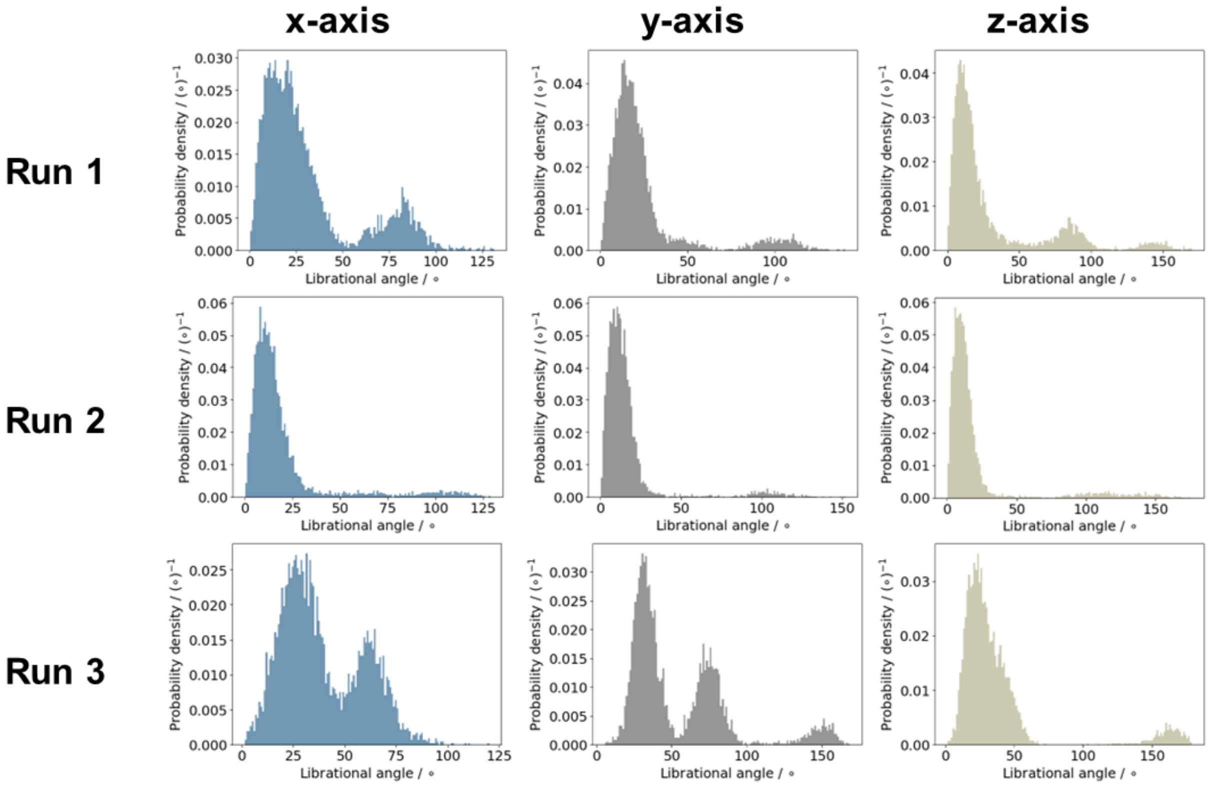


Figure S15: Axis-resolved probability-density distributions of W89H$^{\bullet+}$ librational angles in MagLOV2 for each 500 ns trajectory.

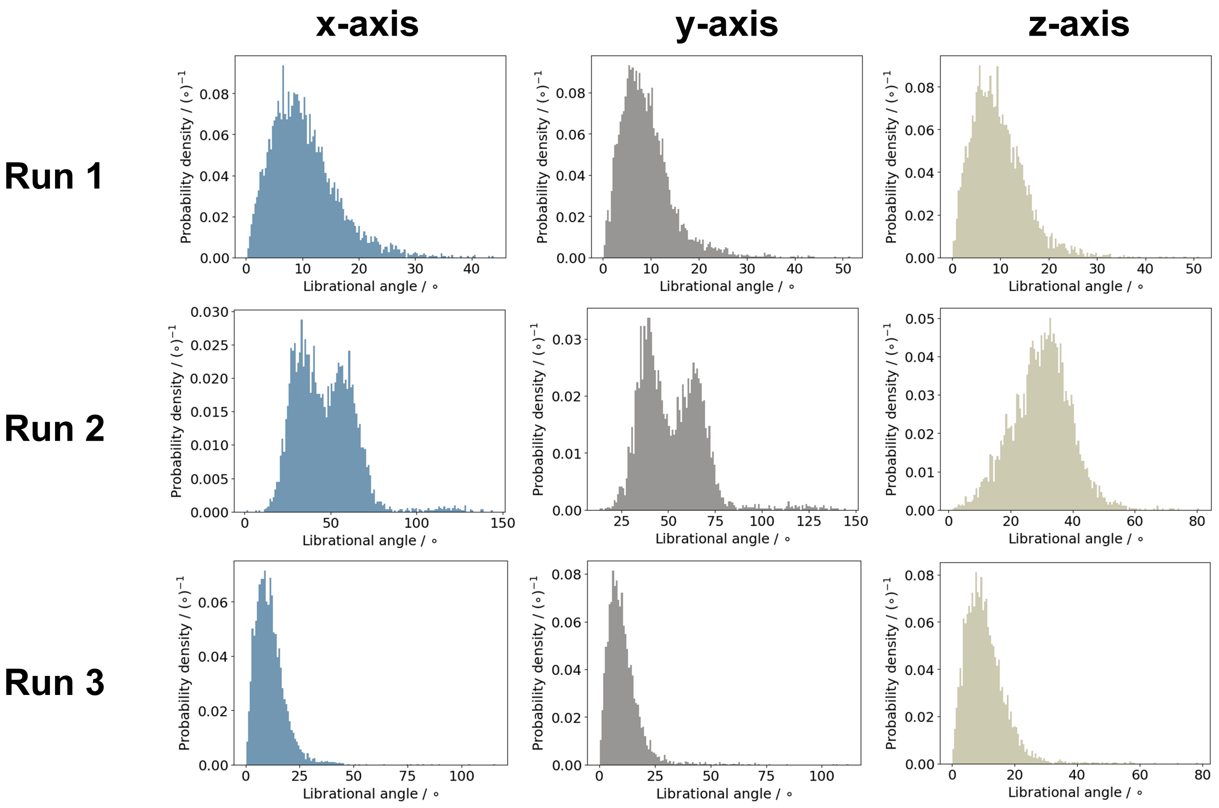


Figure S16: Axis-resolved probability-density distributions of W89H$^{\bullet+}$ librational angles in MagLOV2f for each 500 ns trajectory.

In contrast to FMN, W89H$^{\bullet+}$ samples broad and frequently non-Gaussian orientational distributions (Figs. S12–S16). The behaviour is strongly replica dependent. Individual trajectories can remain in a compact low-angle basin or access additional orientations separated by tens of degrees. MagLOV replica 2, several MagLOV2 replicas, and MagLOV2f replica 2 provide particularly clear examples of such alternative orientational basins. The increasing mutation load, therefore, does not simply increase harmonic fluctuation amplitude. It modifies the accessible donor-side conformational landscape and the exchange kinetics between substates. This distinction is relevant to spin dynamics because broad angular sampling changes the dipolar tensor, whereas slow interconversion between the sampled basins can produce long correlation times.

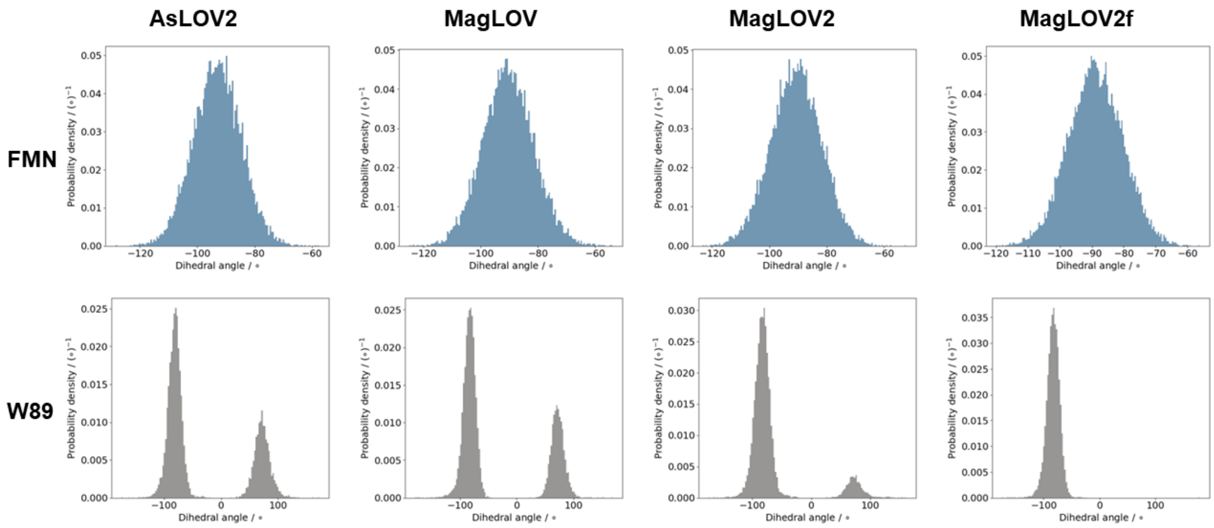


Figure S17: Probability-density distributions of the representative $FMN^{\bullet-}$ and $W89H^{\bullet+}$ dihedral angles for each variant, accumulated over the three 500 ns radical-pair-state trajectories.

The FMN dihedral remains narrowly distributed around the same conformational minimum in all variants (Fig. S17). W89 displays a different trend. *As*LOV2 and MagLOV populate two well-resolved rotameric basins, whereas the population of the second basin is reduced in MagLOV2 and is nearly absent in MagLOV2f. Thus, the internal torsional heterogeneity of W89 decreases with progressive mutation, even though its rigid-body librational sampling becomes broader. The mutations therefore restrict the indole side chain to a dominant rotamer while simultaneously allowing that rotamer to reorient more extensively within the donor pocket.

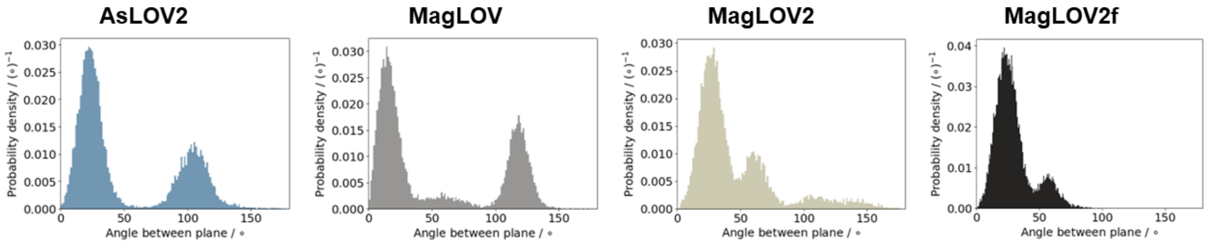


Figure S18: Probability-density distributions of the angle $\alpha$ between the $FMN^{\bullet-}$ and $W89H^{\bullet+}$ aromatic-plane normals for each variant, accumulated over the three 500 ns radical-pair-state trajectories.

The relative plane-angle distributions mirror the W89 rotamer populations (Fig. S18). *As*LOV2 and MagLOV contain two major orientational families, while the higher-angle family loses population in MagLOV2 and is strongly suppressed in MagLOV2f. The evolved variants therefore preferentially populate the lower-angle donor–acceptor arrangement, although residual replica-specific substates remain. Because the normals to a plane have an arbitrary sign, $\alpha$ and $180^\circ - \alpha$ represent the same undirected plane geometry. The electronic-coupling model accordingly uses $|\cos\alpha|$, so that the calculated rate is invariant to this sign convention. These distributions provide a structural basis for variant-dependent electronic coupling and back electron transfer without requiring a large change in the mean COM distance.

### S5 Hydrogen bonding

Hydrogen-bond occupancies were analysed as the number of simultaneous protein–cofactor or protein–W89 hydrogen bonds present in each trajectory frame. The same geometric criterion was applied to every variant, state, and replica, allowing the distributions to be compared directly.

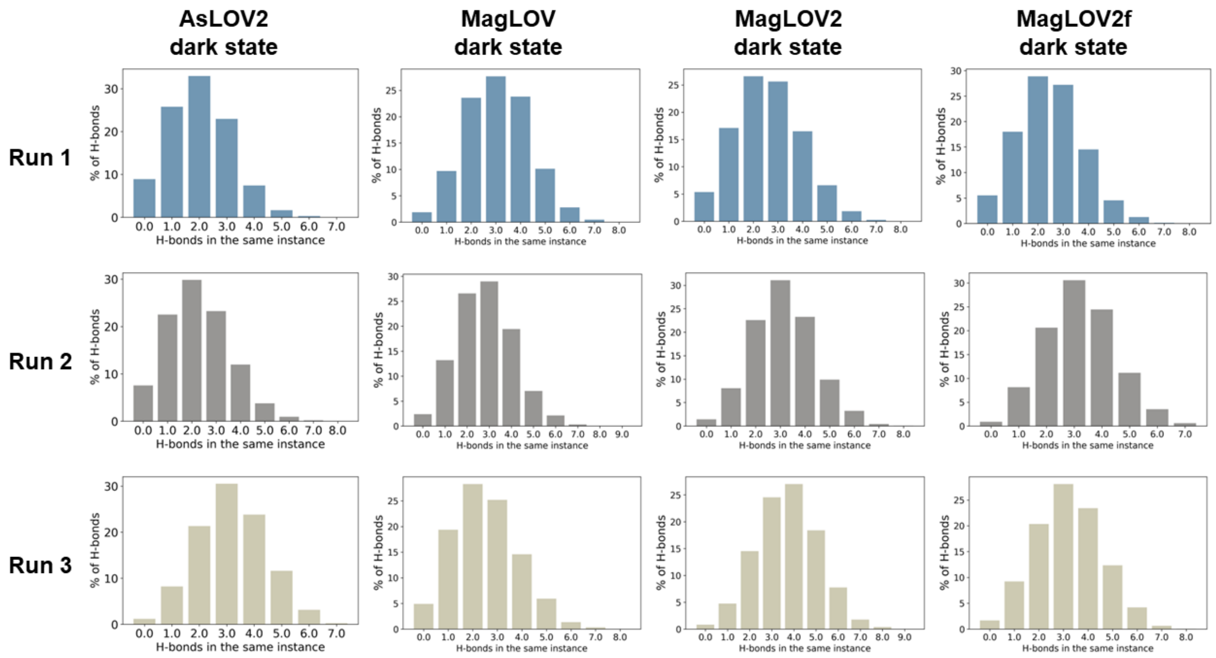


Figure S19: Percentage of dark-state trajectory frames containing the indicated number of simultaneous hydrogen bonds between FMN and its environment. Rows correspond to the three 500 ns replicas and columns to the protein variants.

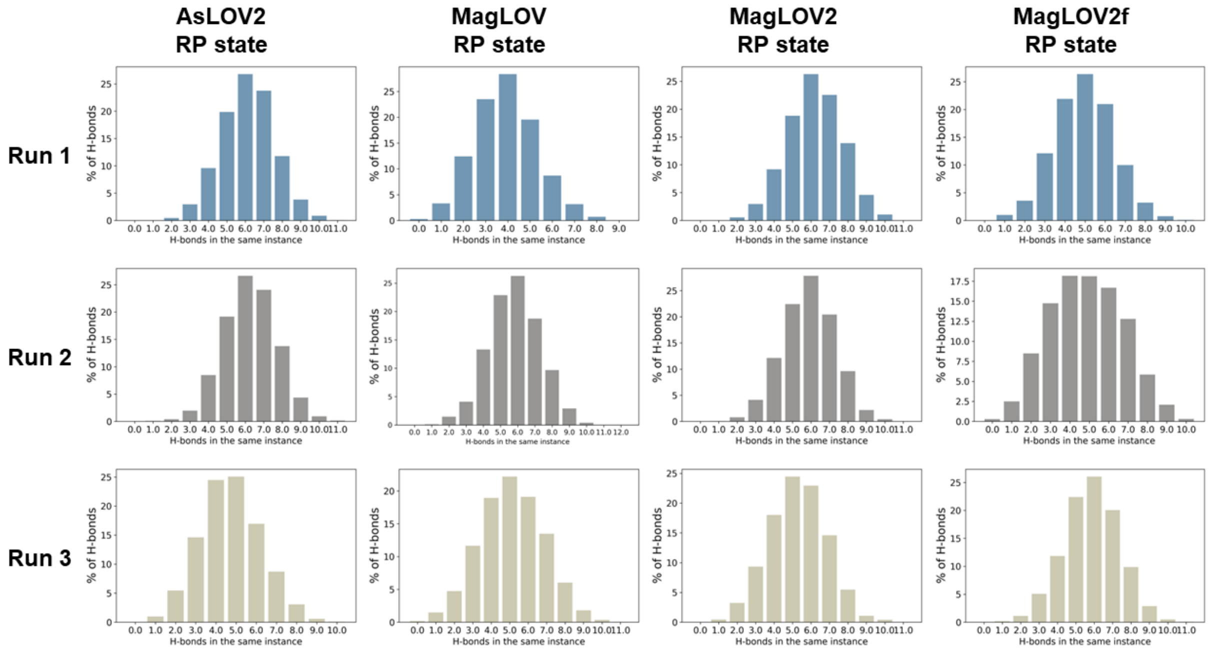


Figure S20: Percentage of radical-pair-state trajectory frames containing the indicated number of simultaneous hydrogen bonds between $FMN^{\bullet-}$ and its environment. Rows correspond to the three 500 ns replicas and columns to the protein variants.

In the dark state, FMN most frequently forms approximately two to four simultaneous hydrogen bonds (Fig. S19). In the radical pair state, the distributions shift to approximately five to seven hydrogen bonds (Fig. S20) for all variants. The increase is consistent with stronger electrostatic and hydrogen-bond stabilisation of the anionic $FMN^{\bullet-}$ state. Although the precise maxima vary between replicas, the state-dependent increase is much larger than the differences among variants. The persistent, multidentate hydrogen-bond network helps explain why FMN remains conformationally restricted throughout the mutation series.

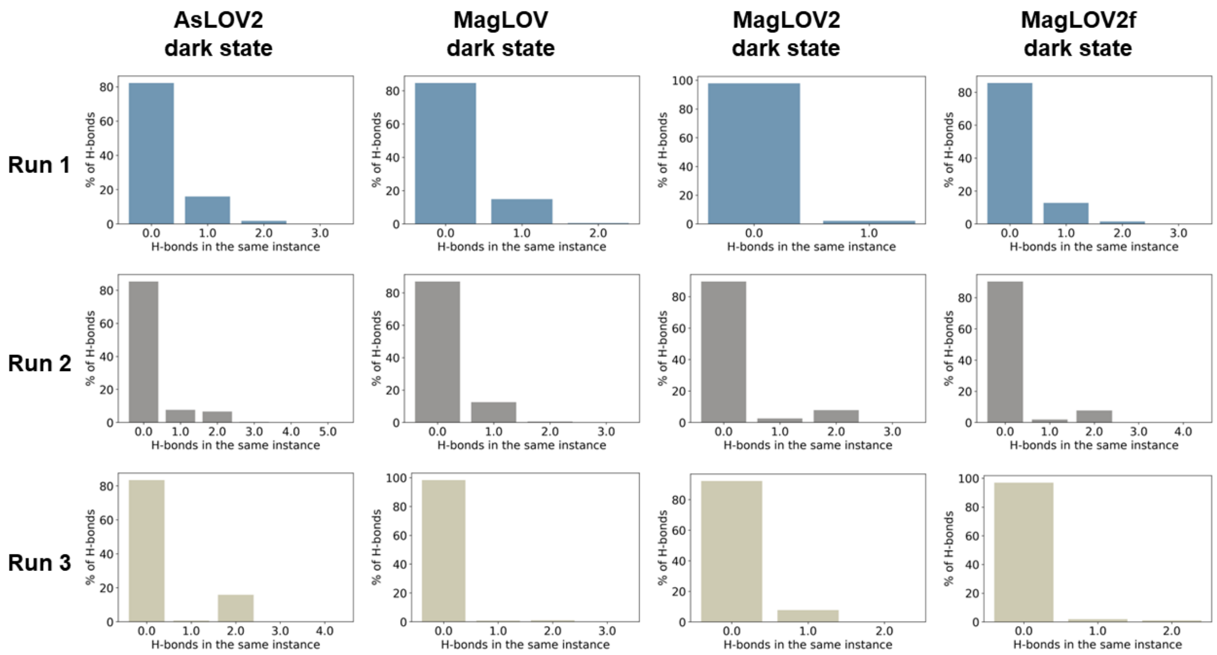


Figure S21: Percentage of dark-state trajectory frames containing the indicated number of simultaneous hydrogen bonds between W89 and its environment. Rows correspond to the three 500 ns replicas and columns to the protein variants.

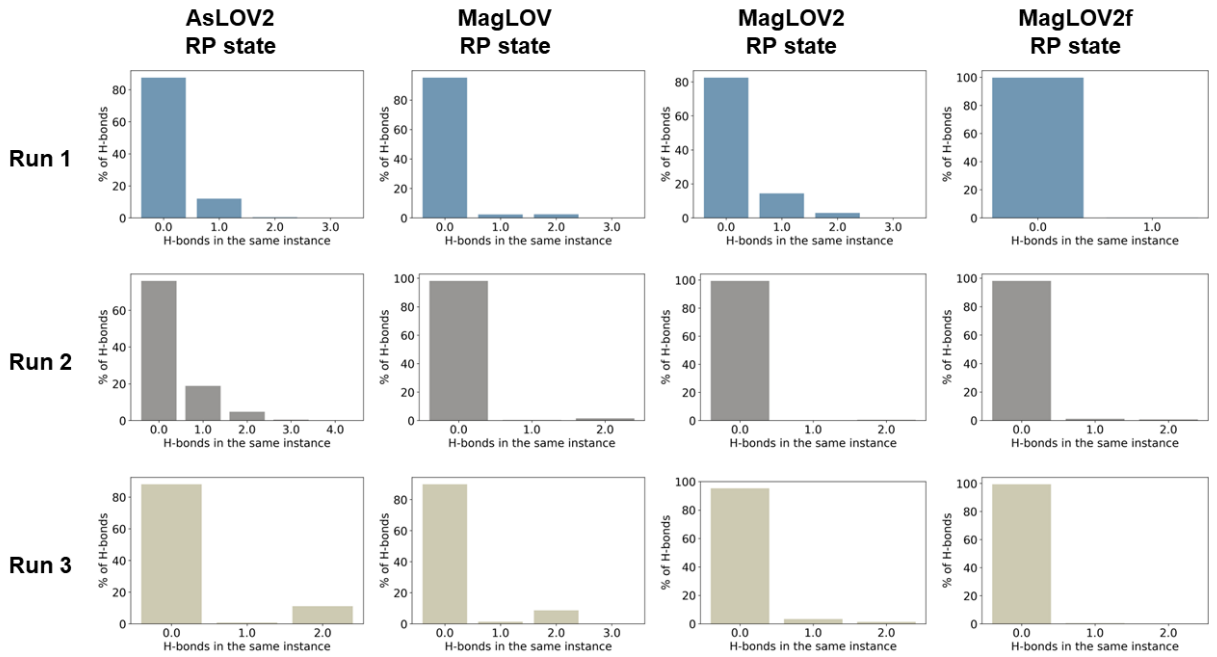


Figure S22: Percentage of radical-pair-state trajectory frames containing the indicated number of simultaneous hydrogen bonds between $W89H^{\bullet+}$ and its environment. Rows correspond to the three 500 ns replicas and columns to the protein variants.

W89 forms no hydrogen bonds in the large majority of frames in either electronic state (Figs. S21 and S22); one or two transient contacts occur only in small subpopulations. No systematic increase in W89 hydrogen bonding accompanies radical-pair formation or progressive mutation. The donor is therefore not immobilised by a persistent hydrogen-bond network. Its conformational stability must instead arise mainly from steric packing, hydrophobic contacts, and the surrounding electrostatic field, making it more susceptible than FMN to mutation-dependent orientational fluctuations.

### S6 Exchange interaction

The isotropic exchange interaction was represented phenomenologically as an exponential function of the instantaneous inter-radical separation $r(t)$,

$$J(t) = J\big(r(t)\big) = J_0 \exp\big[-\beta r(t)\big]. \tag{S3}$$

The parameters $J_0 = 786.54$ mT and $\beta = 6.31 \times 10^9$ m$^{-1}$ were obtained from experimental exchange couplings measured by out-of-phase electron spin echo envelope modulation for two radical pairs in avian cryptochrome 4a.[1] Specifically, the values $J_{RP_D} = 0.001$ mT at $r_{RP_D} = 2.14$ nm and $J_{RP_C} = 0.011$ mT at $r_{RP_C} = 1.77$ nm were related through

$$\beta = \frac{\ln(J_{RP_C}/J_{RP_D})}{r_{RP_D} - r_{RP_C}}, \tag{S4}$$

and

$$J_0 = J_{RP_D} \exp(\beta r_{RP_D}). \tag{S5}$$

The numerical values above retain the parameterisation obtained from the experimental data used in this study. Because this calibration is transferred from cryptochrome to the LOV scaffold, it should be interpreted as a physically motivated order-of-magnitude model rather than a system-specific electronic-structure calculation of $J$.

The resulting exchange couplings remain below approximately 100 $\mu$T over the sampled distance range and are comparable to values reported for *Cr*LOV1 C57S.[2,3] When the trajectory-derived fluctuations were propagated through the same correlation-function treatment used for the dipolar interaction, the exchange-modulation contribution was substantially smaller than the dipolar contribution. Exchange-induced singlet–triplet dephasing was therefore neglected in the subsequent spin-dynamics analysis. This conclusion is based on the calculated fluctuation integral, rather than on the mean value of $J$ alone.

### S7 Centre-to-centre distances

The relative position of the two proposed radical-pair partners was first characterised by the centre-of-mass (COM) distance between FMN and W89. This descriptor provides a global measure of donor–acceptor separation, although electron transfer and electron–electron dipolar coupling additionally depend on the edge-to-edge distance, spin-density distribution, and relative orientation of the two moieties.

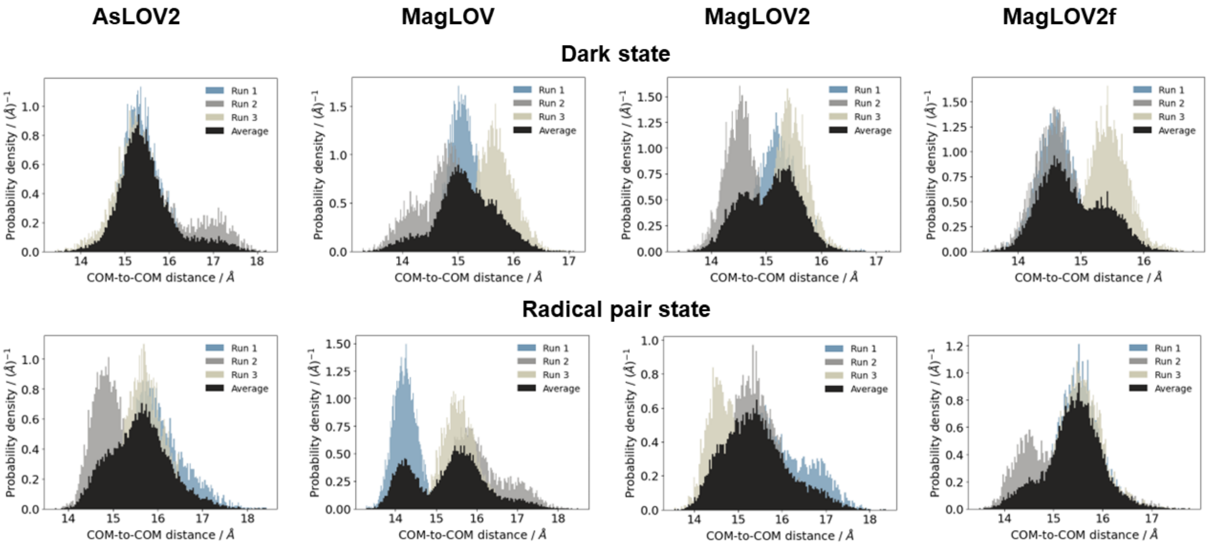


Figure S23: Probability-density distributions of the FMN–W89 COM distance for each variant in the dark state and radical pair state. The coloured distributions correspond to the three independent replicas and the black distribution represents their equally weighted average.

The ensemble-averaged COM distances are closely clustered, with mean values between approximately 15.30 and 15.41 Å. Thus, directed evolution does not produce a monotonic expansion or contraction of the native FMN–W89 pair. The principal differences instead appear in the widths and multimodal character of the distributions, indicating that the variants sample different populations of donor–acceptor substates. This replica dependence is particularly pronounced for MagLOV in the radical pair state, where a short-distance population near 14–15 Å coexists with a broader population near 15.5–16 Å. The latter is associated with the conformational transition seen in Fig. S1. These results show that a similar mean separation can conceal substantially different conformational gating of the radical pair.

### S8 Spin relaxation

#### S8.1 Dipolar coupling modulation

Fluctuations of the electron–electron dipolar tensor were characterised by trajectory-derived autocorrelation functions. The solid curves in Fig. S24 represent the correlation functions obtained from the MD time series, and the dashed curves are the non-negative multi-exponential fits used to determine the effective, amplitude-weighted correlation times. A longer correlation time indicates that a particular dipolar environment persists for longer before motional decorrelation.

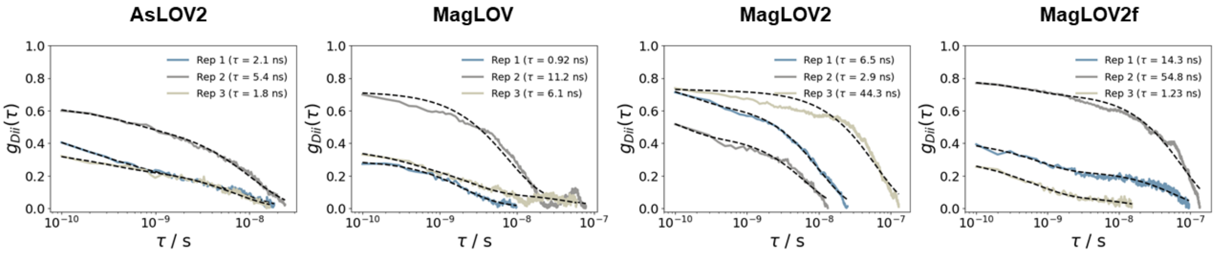


Figure S24: Dipolar-tensor autocorrelation functions for the three 500 ns radical-pair-state replicas of *As*LOV2, MagLOV, MagLOV2, and MagLOV2f. Solid curves are obtained directly from the MD trajectories and dashed curves show the fitted correlation functions. The effective correlation time $\tau_c$ extracted for each replica is reported in the corresponding legend.

All three *As*LOV2 replicas decorrelate on relatively short timescales, with $\tau_c$ = 1.8–5.4 ns. MagLOV spans 0.92–11.2 ns, whereas MagLOV2 and MagLOV2f each contain replicas with markedly slower components, reaching 44.3 and 54.8 ns, respectively. The evolved variants, therefore, access dipolar environments that can remain correlated for tens of nanoseconds, but the large replica-to-replica spread shows that these long-lived substates are not sampled uniformly. This heterogeneity is consistent with the multimodal W89 librational distributions.

The correlation time alone does not determine the dipolar-induced relaxation rate. Within the adopted zero-frequency stochastic-modulation treatment, $k_D$ is proportional to the time integral of the dipolar autocovariance (with the dipolar tensor expressed in angular-frequency units) and therefore depends on both the fluctuation amplitude $\langle \delta d^2 \rangle$ and its persistence $\tau_c$. Consequently, a long $\tau_c$ enhances relaxation only when accompanied by appreciable dipolar-tensor variance. The present figure establishes the slower dynamical component. The rate comparison in the main manuscript additionally incorporates the fluctuation amplitudes.

#### S8.2 Hyperfine coupling modulation

Protein dynamics modulate the electron–nuclear hyperfine coupling (HFC) tensors of the $FMN^{\bullet-}$ and $TrpH^{\bullet+}$ radicals, thereby providing a molecular contribution to spin relaxation. To resolve these fluctuations, HFC tensors were evaluated along each of the three 500 ns radical-pair-state MD trajectories of *As*LOV2, MagLOV, MagLOV2, and MagLOV2f. The labels in the following plots follow the convention illustrated in Fig. S25.

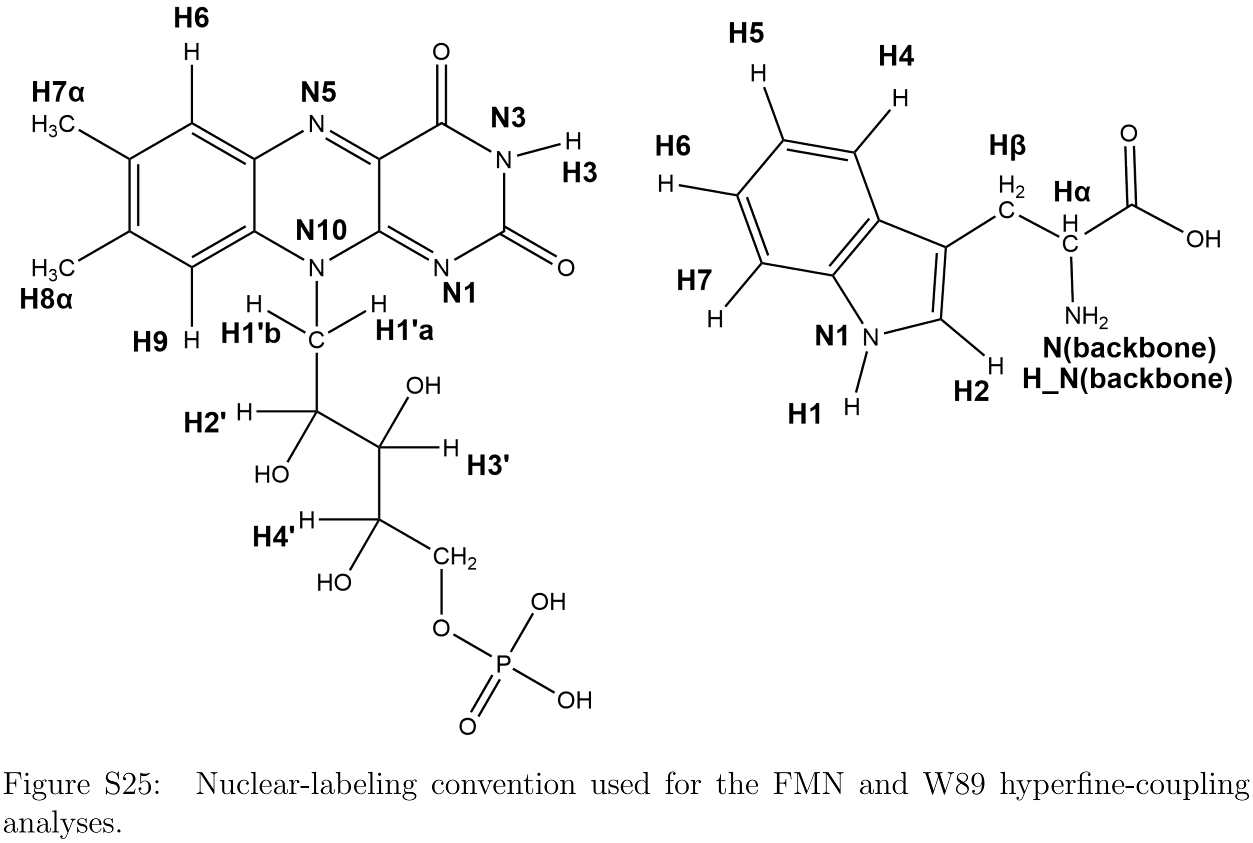


Figure S25: Nuclear-labeling convention used for the FMN and W89 hyperfine-coupling analyses.

Structures were sampled at 100 ps intervals, yielding 5000 HFC tensors per radical and trajectory. The HFC data set used for the spin-relaxation analysis is shown throughout this section. The electronic-structure protocol is identical to that specified in the main-text Methods; no variant-dependent change of method or nuclear selection was introduced.

For the comparison between variants, the plotted HFC entities were selected globally for each radical according to the trajectory-averaged tensor magnitude,

$$A_{\mathrm{RMS}} = \sqrt{\langle \|\mathbf{A}(t)\|_{\mathrm{F}}^{2} \rangle / 3}, \tag{S6}$$

such that the same nuclei are compared for all protein variants and replicas. Equivalent methyl protons of FMN and the two Trp H$\beta$ protons were averaged only for visualisation; the individual nuclear spins were retained separately in the relaxation analysis. Artificial capping nuclei of the truncated Trp fragment were excluded.

##### S8.2.1 FMN hyperfine coupling modulation

The FMN HFC trajectories characterise how the motions of the isoalloxazine and ribityl moieties modulate the magnetic interaction between the flavin electron spin and the surrounding magnetic nuclei. The same set of dominant FMN HFC is shown for all four variants and all three independent MD replicas, allowing for differences in both the magnitude and temporal persistence of the HFC fluctuations to be compared directly.

**Isotropic hyperfine fluctuations** The isotropic component of each HFC tensor was calculated as

$$A_{\mathrm{iso}}(t) = \frac{1}{3}\mathrm{Tr}\left[\mathbf{A}(t)\right]. \tag{S7}$$

The following figures show the time-dependent $A_{\mathrm{iso}}$ values of the 15 dominant FMN HFC entities over each 500 ns trajectory. Dashed horizontal lines indicate the trajectory-averaged isotropic coupling. These profiles, therefore, distinguish changes in the mean HFC from fluctuations around that mean and reveal slower conformational shifts, where the isotropic coupling remains displaced over extended parts of a trajectory.

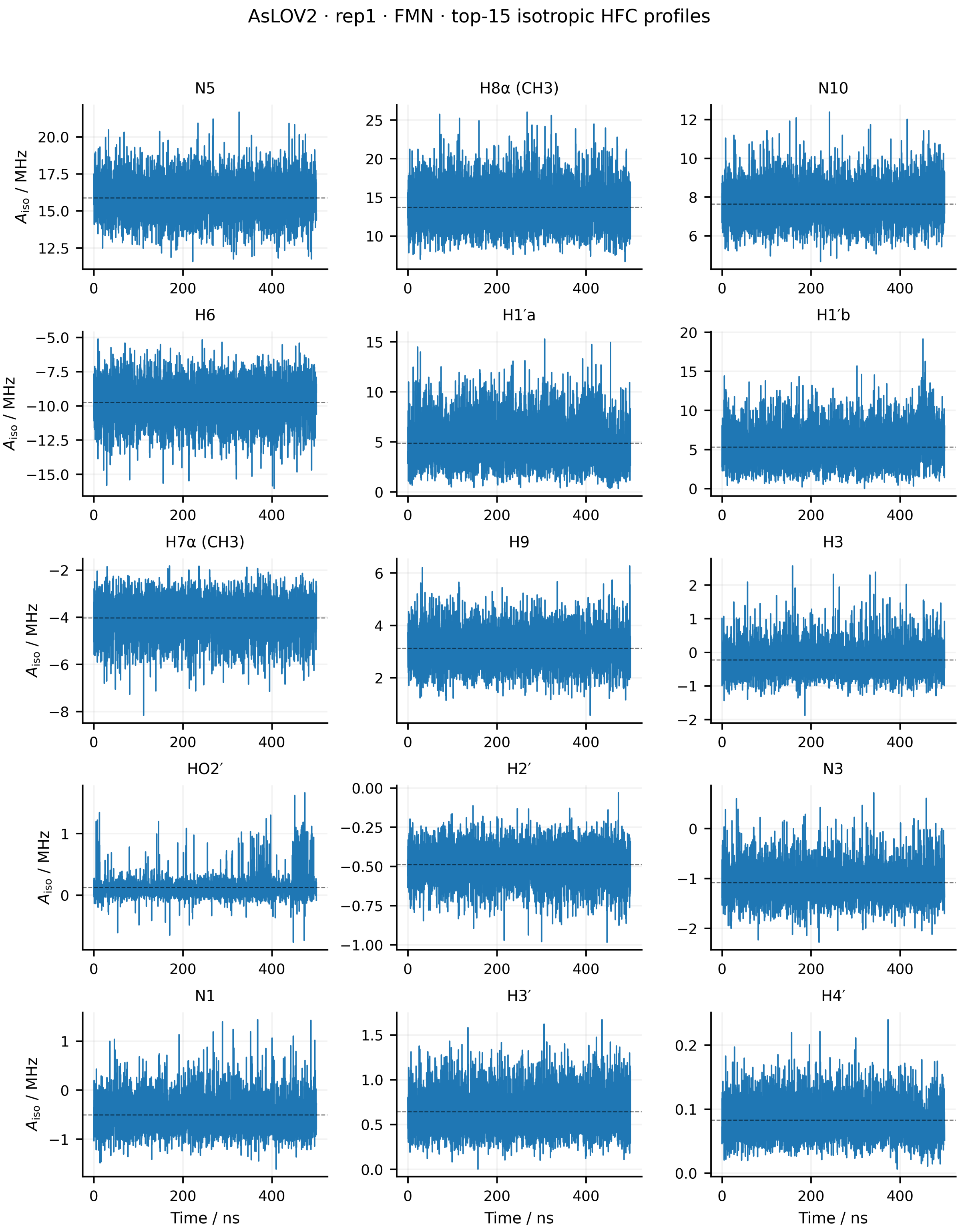


Figure S26: Time evolution of the isotropic hyperfine couplings of the 15 dominant FMN HFC entities for replica 1 of *As*LOV2. The isotropic coupling was calculated as $A_{\mathrm{iso}}(t) = \mathrm{Tr}[\mathbf{A}(t)]/3$. Dashed horizontal lines indicate the mean value over the 500 ns trajectory. Equivalent methyl-proton groups are averaged for visualisation only.

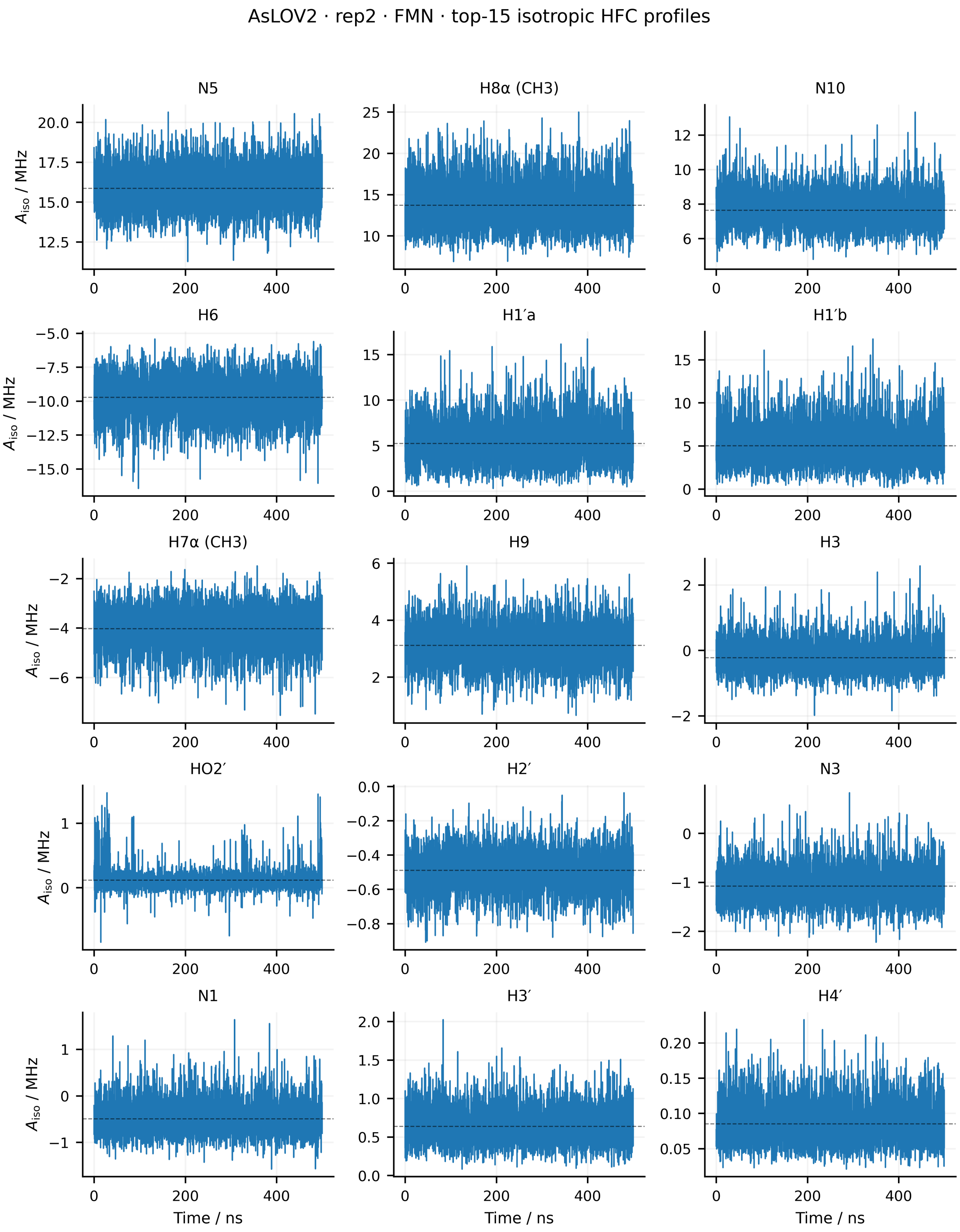


Figure S27: Time evolution of the isotropic hyperfine couplings of the 15 dominant FMN HFC entities for replica 2 of *As*LOV2. The isotropic coupling was calculated as $A_{\mathrm{iso}}(t) = \mathrm{Tr}[\mathbf{A}(t)]/3$. Dashed horizontal lines indicate the mean value over the 500 ns trajectory. Equivalent methyl-proton groups are averaged for visualisation only.

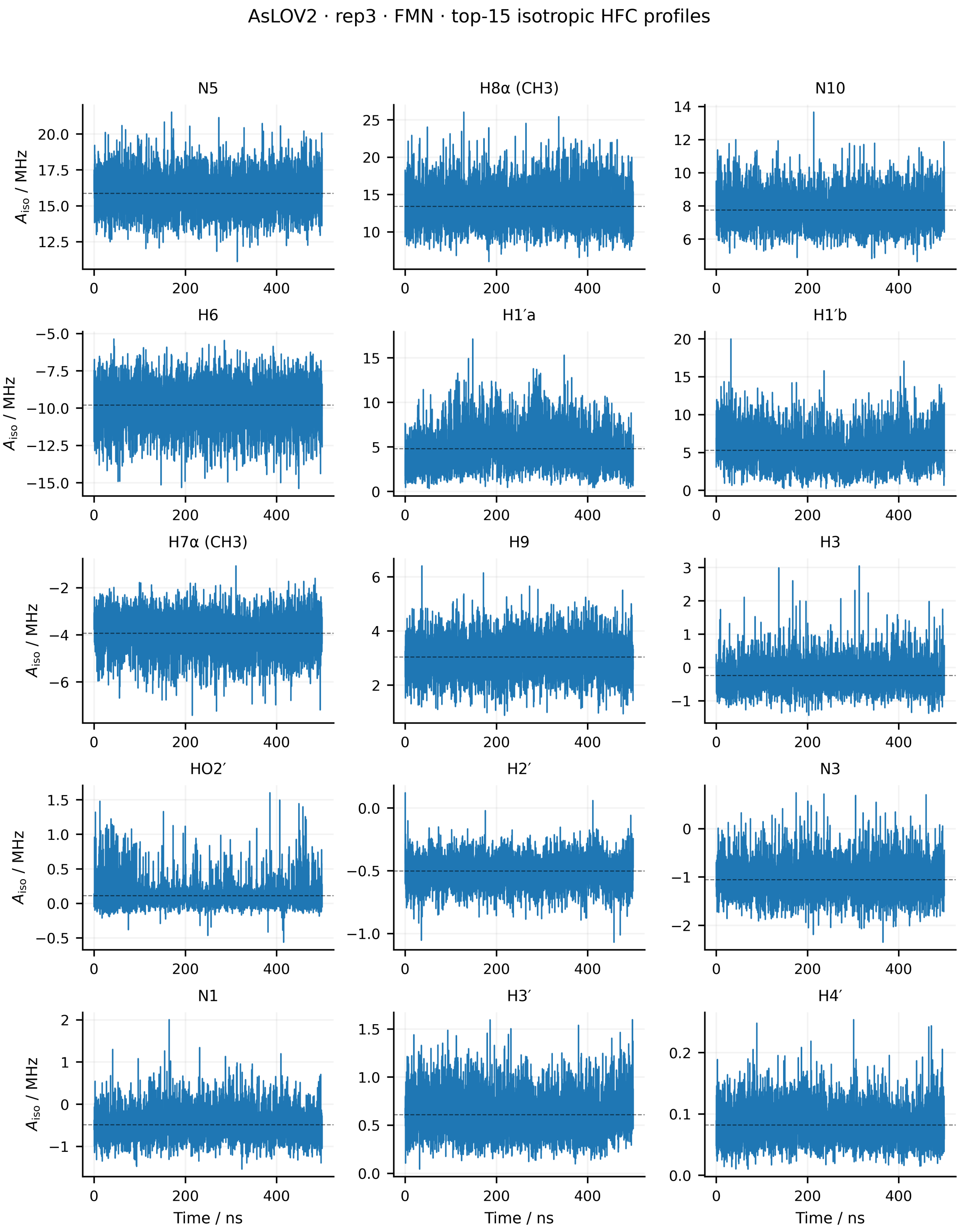


Figure S28: Time evolution of the isotropic hyperfine couplings of the 15 dominant FMN HFC entities for replica 3 of *As*LOV2. The isotropic coupling was calculated as $A_{\mathrm{iso}}(t) = \mathrm{Tr}[\mathbf{A}(t)]/3$. Dashed horizontal lines indicate the mean value over the 500 ns trajectory. Equivalent methyl-proton groups are averaged for visualisation only.

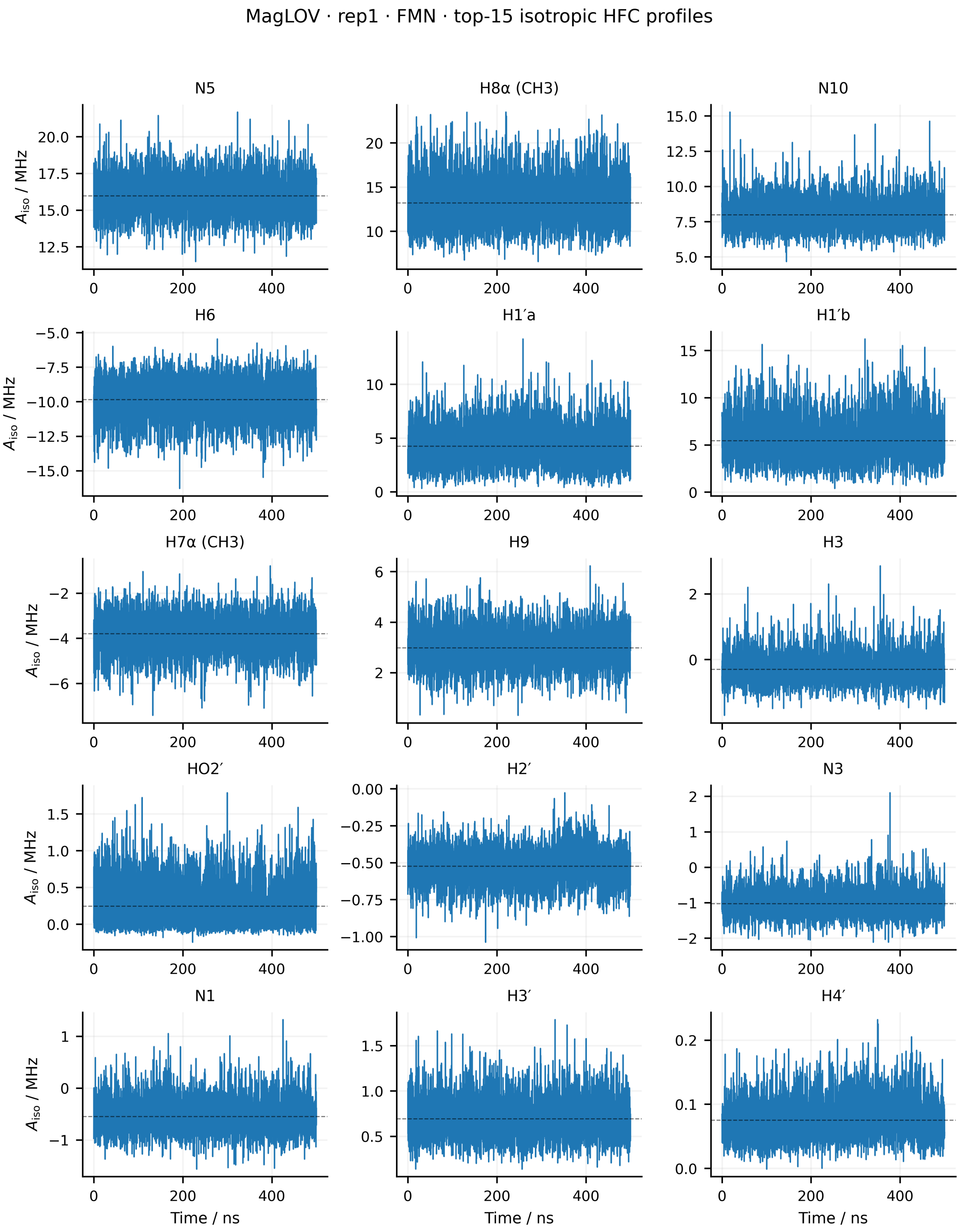


Figure S29: Time evolution of the isotropic hyperfine couplings of the 15 dominant FMN HFC entities for replica 1 of MagLOV. The isotropic coupling was calculated as $A_{\text{iso}}(t) = \text{Tr}[\mathbf{A}(t)]/3$. Dashed horizontal lines indicate the mean value over the 500 ns trajectory. Equivalent methyl-proton groups are averaged for visualisation only.

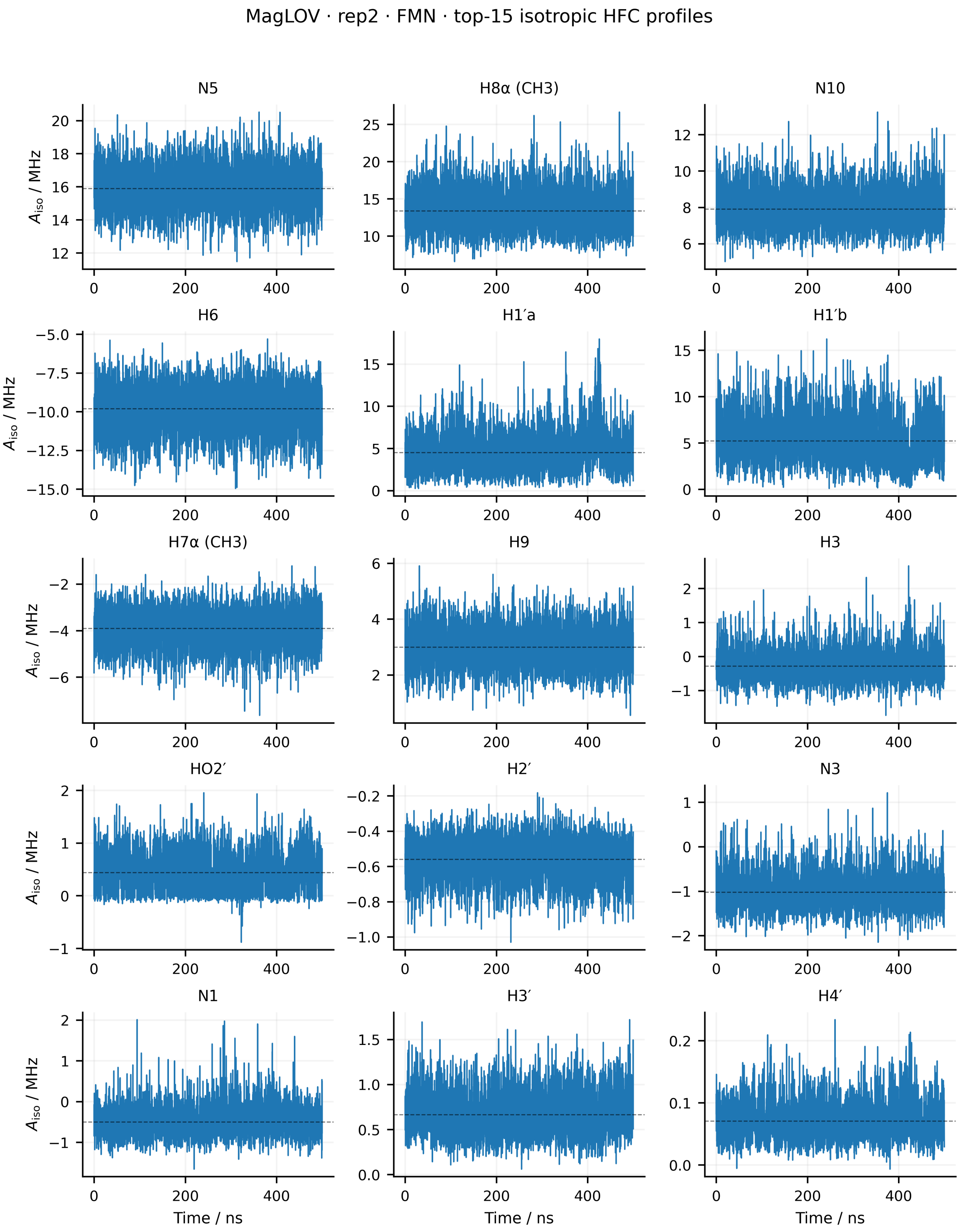


Figure S30: Time evolution of the isotropic hyperfine couplings of the 15 dominant FMN HFC entities for replica 2 of MagLOV. The isotropic coupling was calculated as $A_{\mathrm{iso}}(t) = \mathrm{Tr}[\mathbf{A}(t)]/3$. Dashed horizontal lines indicate the mean value over the 500 ns trajectory. Equivalent methyl-proton groups are averaged for visualisation only.

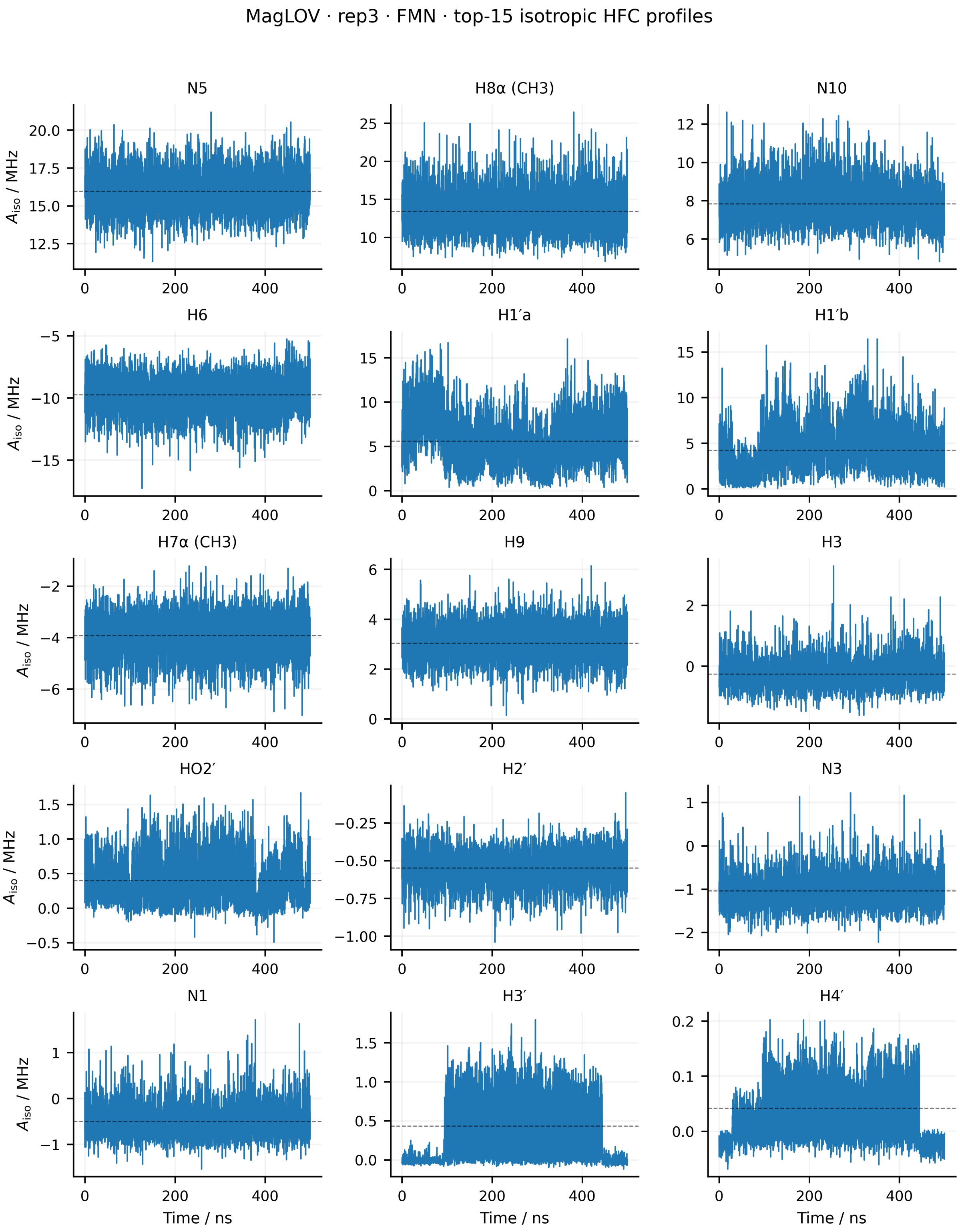


Figure S31: Time evolution of the isotropic hyperfine couplings of the 15 dominant FMN HFC entities for replica 3 of MagLOV. The isotropic coupling was calculated as $A_{\text{iso}}(t) = \text{Tr}[\mathbf{A}(t)]/3$. Dashed horizontal lines indicate the mean value over the 500 ns trajectory. Equivalent methyl-proton groups are averaged for visualisation only.

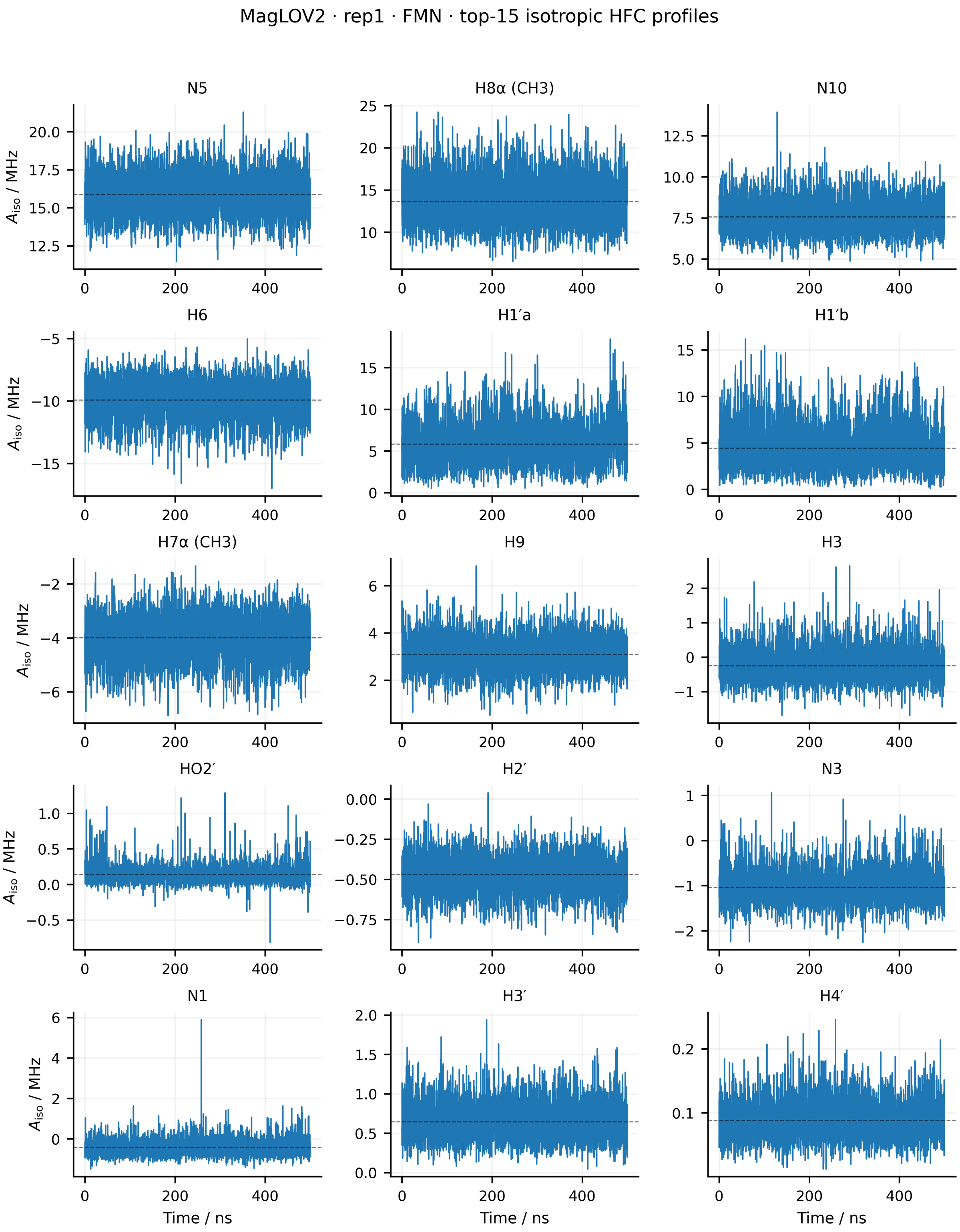


Figure S32: Time evolution of the isotropic hyperfine couplings of the 15 dominant FMN HFC entities for replica 1 of MagLOV2. The isotropic coupling was calculated as $A_{\mathrm{iso}}(t) = \mathrm{Tr}[\mathbf{A}(t)]/3$. Dashed horizontal lines indicate the mean value over the 500 ns trajectory. Equivalent methyl-proton groups are averaged for visualisation only.

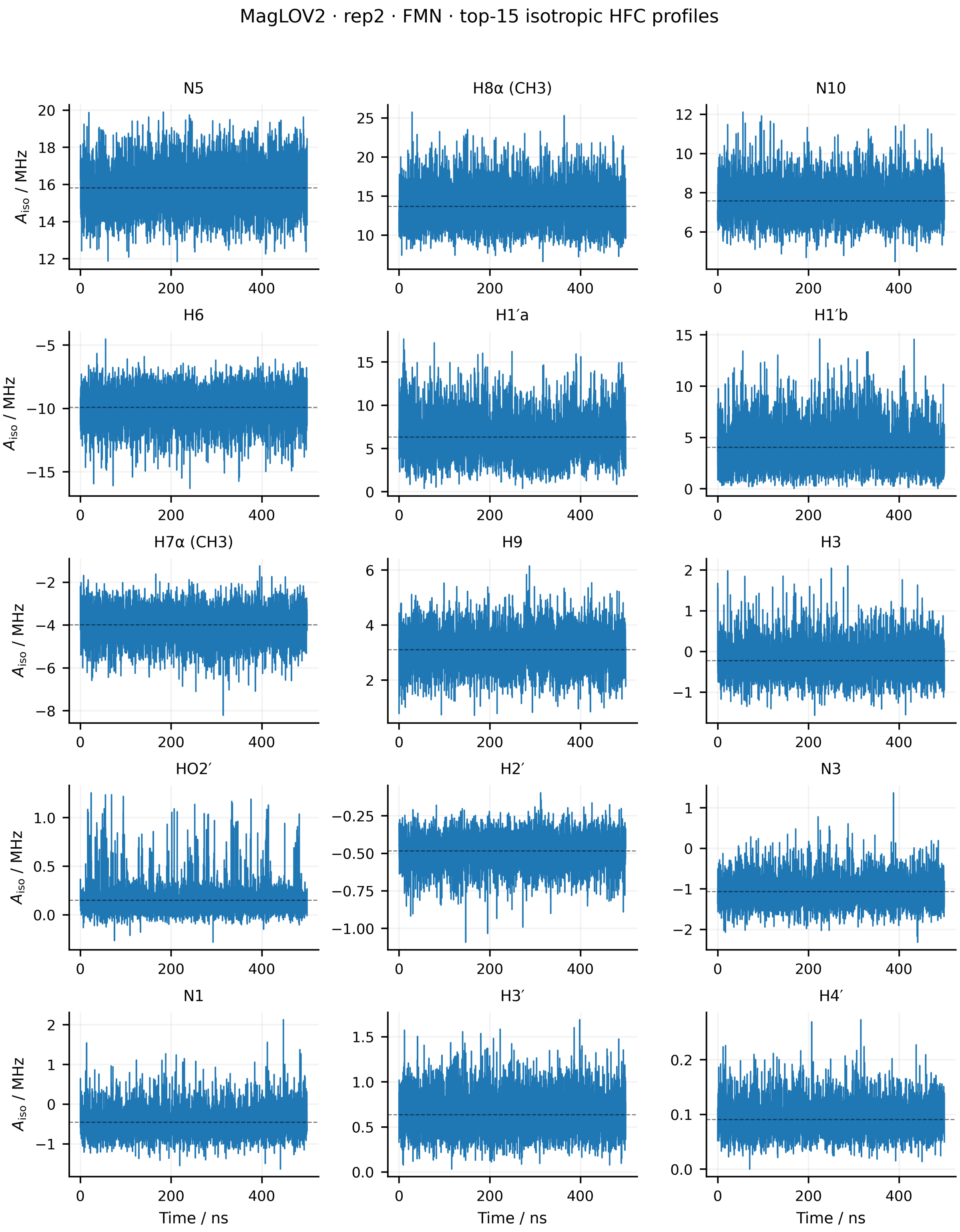


Figure S33: Time evolution of the isotropic hyperfine couplings of the 15 dominant FMN HFC entities for replica 2 of MagLOV2. The isotropic coupling was calculated as $A_{\mathrm{iso}}(t) = \mathrm{Tr}[\mathbf{A}(t)]/3$. Dashed horizontal lines indicate the mean value over the 500 ns trajectory. Equivalent methyl-proton groups are averaged for visualisation only.

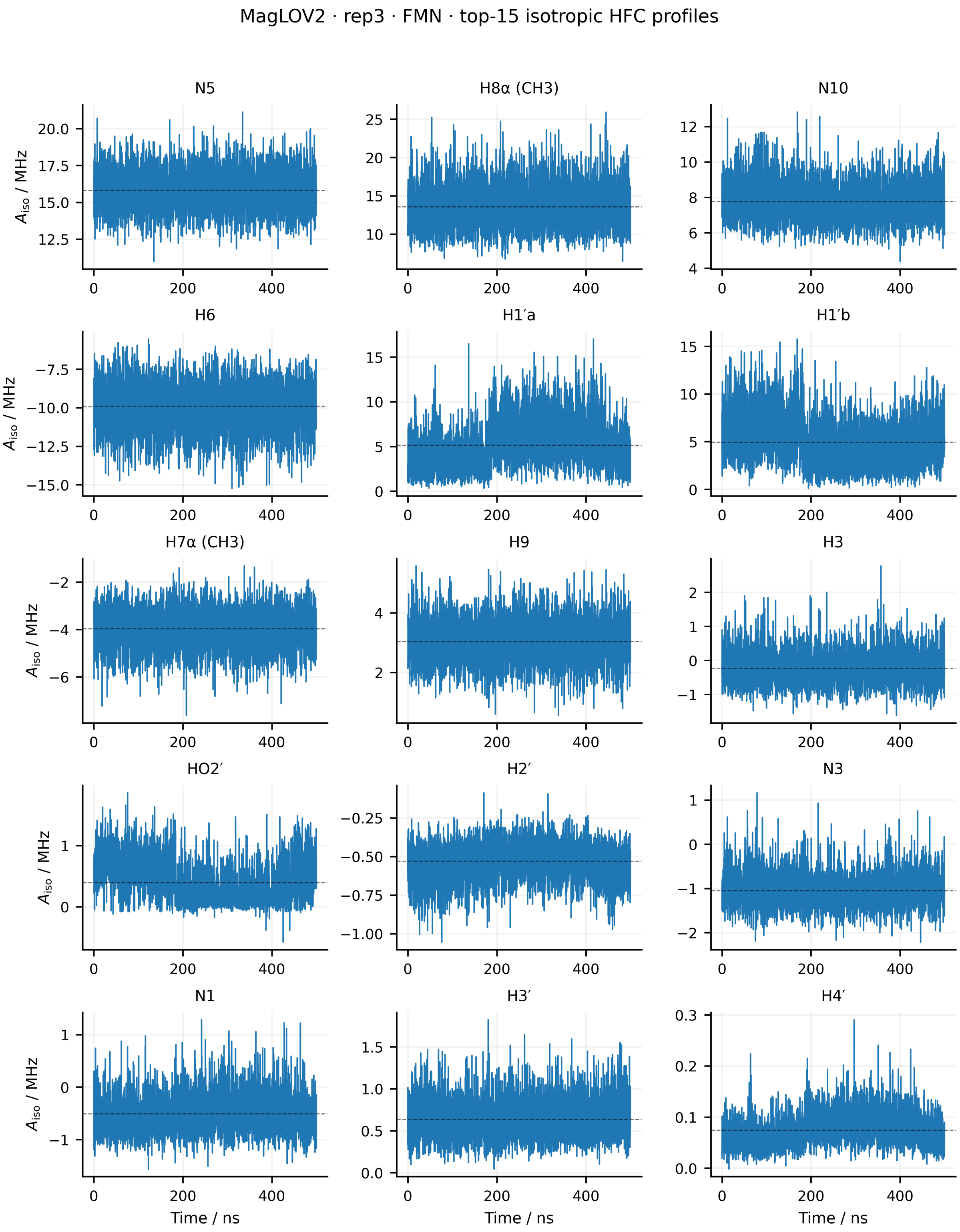


Figure S34: Time evolution of the isotropic hyperfine couplings of the 15 dominant FMN HFC entities for replica 3 of MagLOV2. The isotropic coupling was calculated as $A_{\mathrm{iso}}(t) = \mathrm{Tr}[\mathbf{A}(t)]/3$. Dashed horizontal lines indicate the mean value over the 500 ns trajectory. Equivalent methyl-proton groups are averaged for visualisation only.

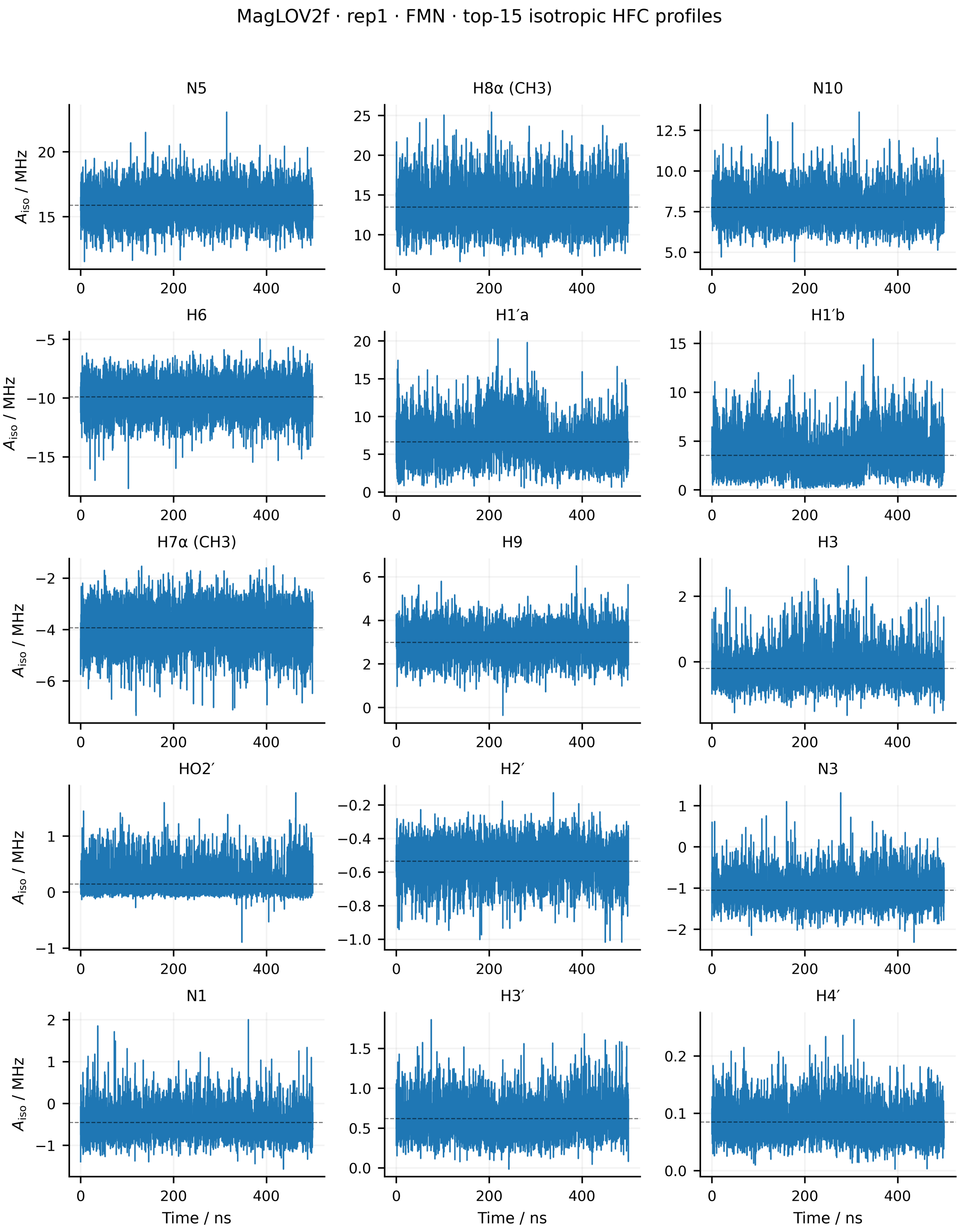


Figure S35: Time evolution of the isotropic hyperfine couplings of the 15 dominant FMN HFC entities for replica 1 of MagLOV2f. The isotropic coupling was calculated as $A_{\text{iso}}(t) = \text{Tr}[\mathbf{A}(t)]/3$. Dashed horizontal lines indicate the mean value over the 500 ns trajectory. Equivalent methyl-proton groups are averaged for visualisation only.

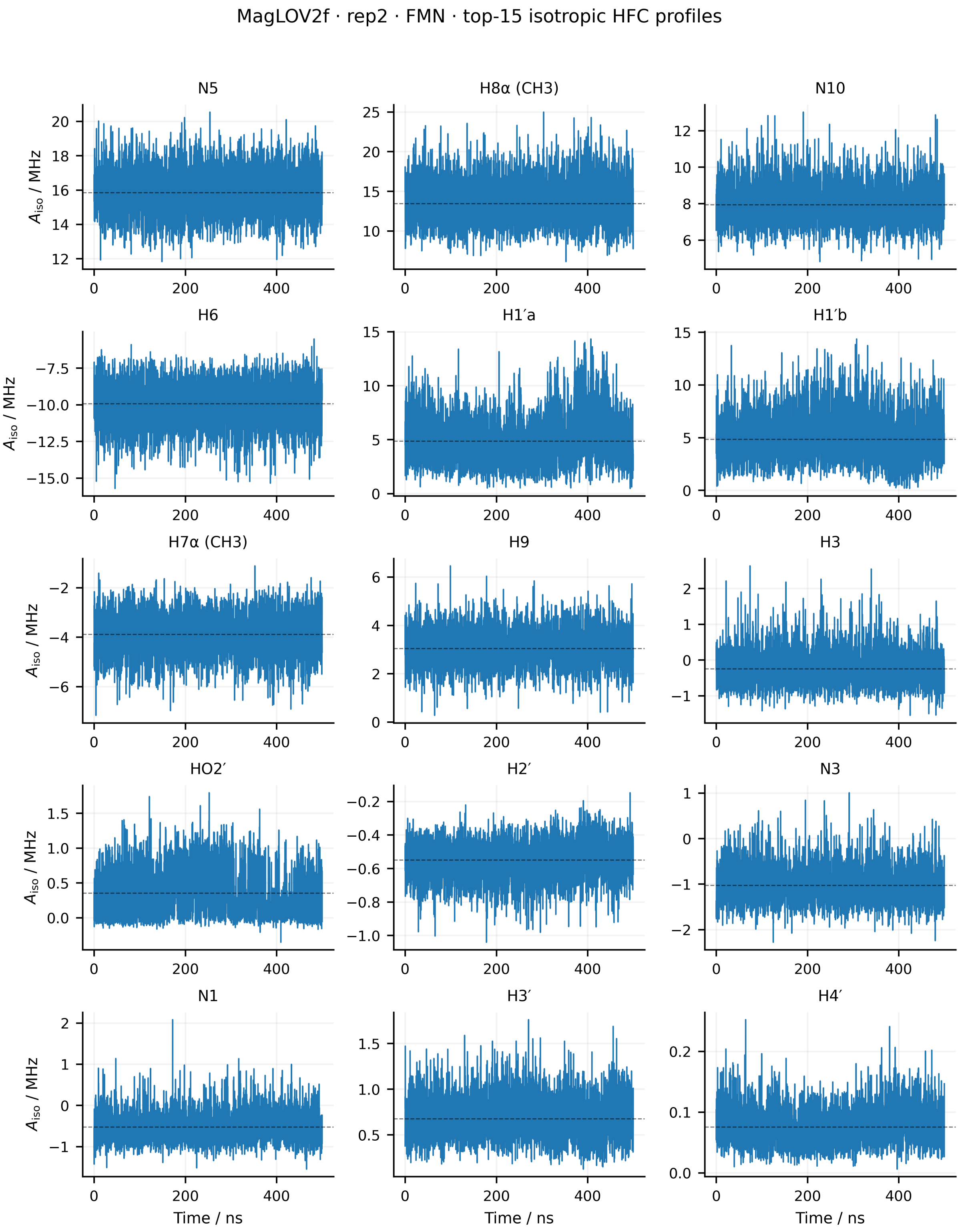


Figure S36: Time evolution of the isotropic hyperfine couplings of the 15 dominant FMN HFC entities for replica 2 of MagLOV2f. The isotropic coupling was calculated as $A_{\text{iso}}(t) = \text{Tr}[\mathbf{A}(t)]/3$. Dashed horizontal lines indicate the mean value over the 500 ns trajectory. Equivalent methyl-proton groups are averaged for visualisation only.

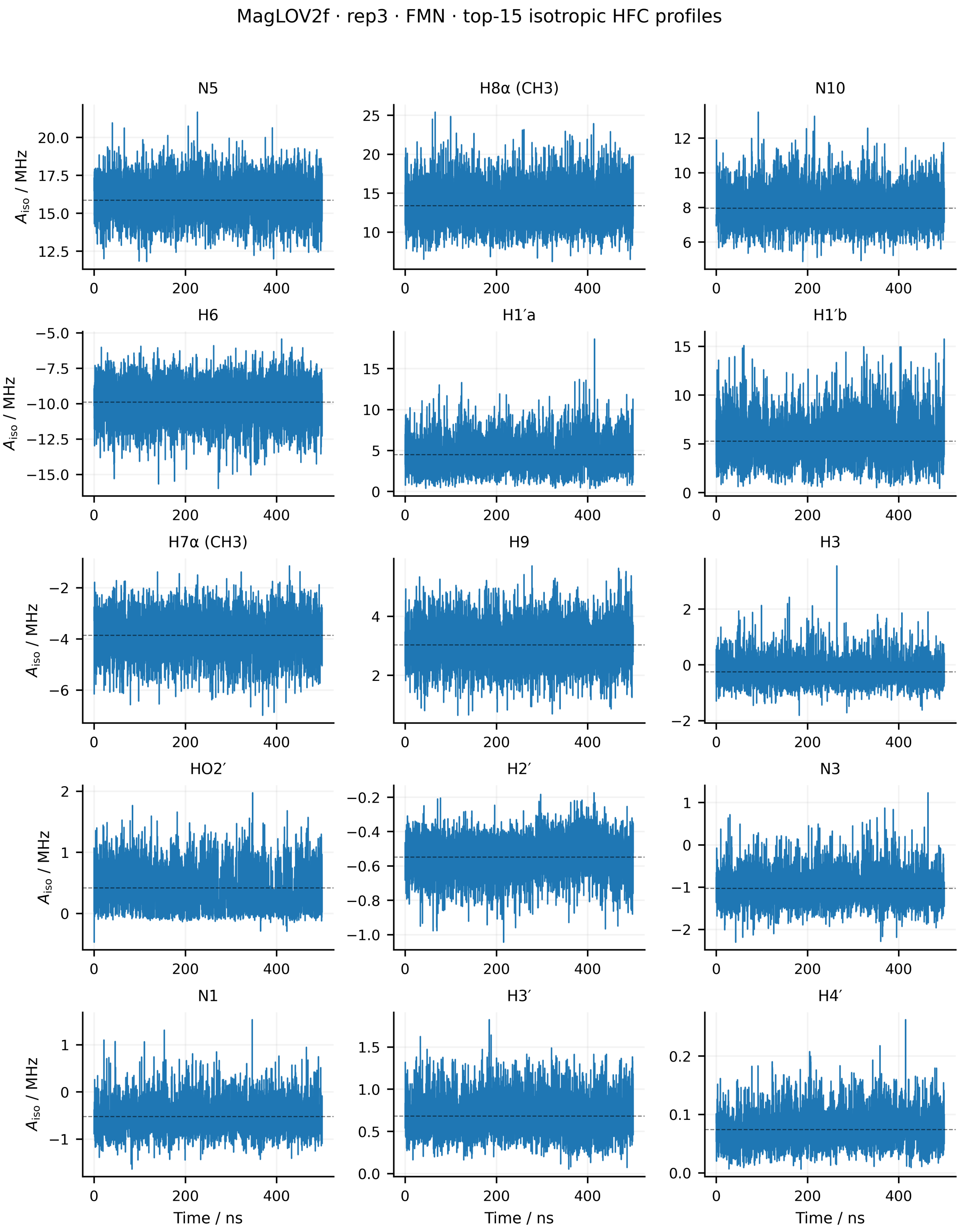


Figure S37: Time evolution of the isotropic hyperfine couplings of the 15 dominant FMN HFC entities for replica 3 of MagLOV2f. The isotropic coupling was calculated as $A_{\rm iso}(t) = {\rm Tr}[\mathbf{A}(t)]/3$. Dashed horizontal lines indicate the mean value over the 500 ns trajectory. Equivalent methyl-proton groups are averaged for visualisation only.

**Full hyperfine tensor fluctuations** Spin relaxation depends on fluctuations of the complete HFC tensor rather than on its isotropic component alone. The instantaneous full-tensor fluctuation amplitude was therefore evaluated relative to the trajectory-averaged tensor according to

$$\Delta A_{\text{tensor}}(t) = \frac{\|\mathbf{A}(t) - \langle \mathbf{A} \rangle\|_{\text{F}}}{\sqrt{3}}. \tag{S8}$$

This quantity includes fluctuations of both the isotropic and anisotropic components of the HFC tensor. The resulting time profiles identify periods of enhanced tensor modulation and form the underlying trajectory data from which the HFC autocorrelation functions and relaxation contributions are determined.

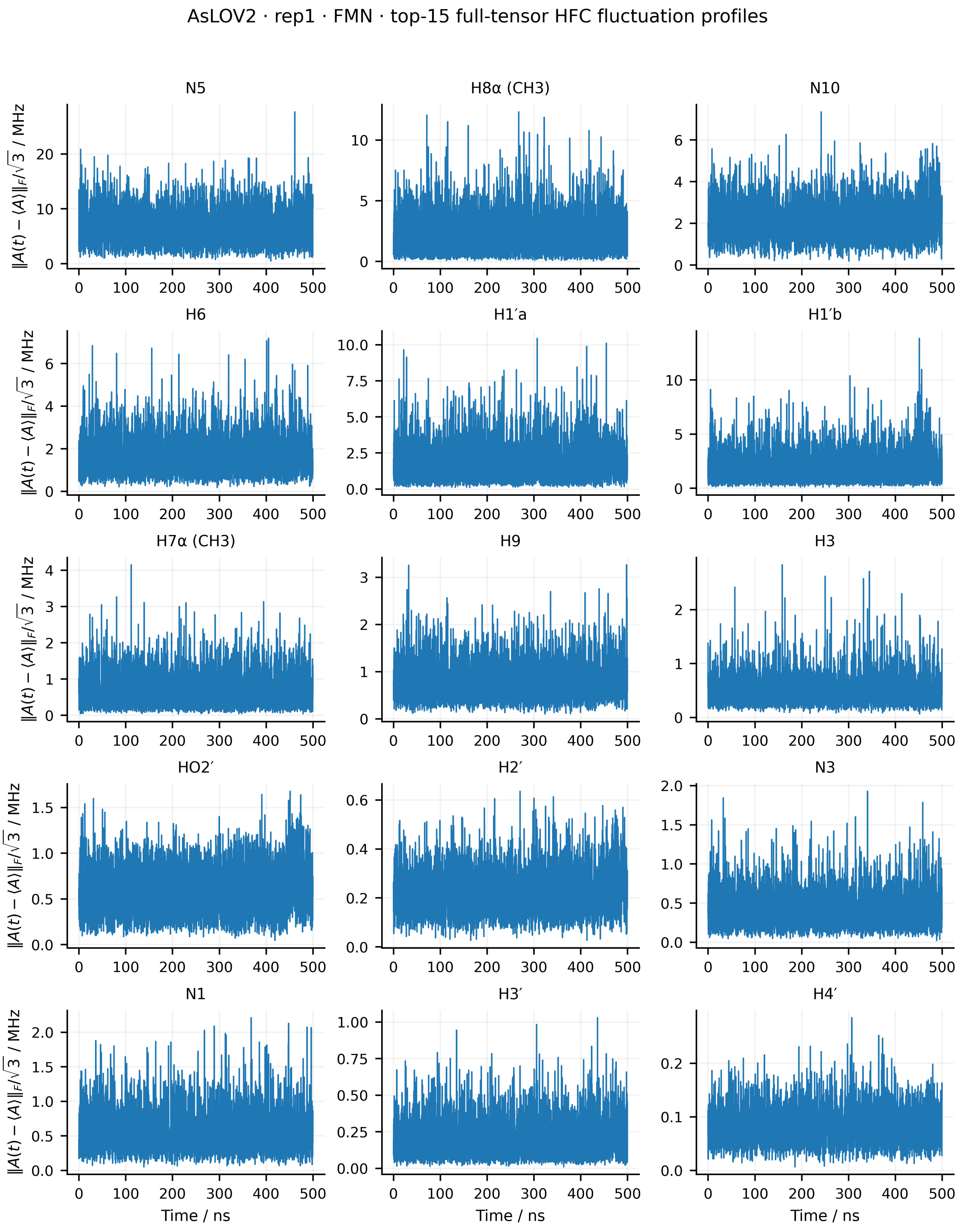


Figure S38: Full-tensor hyperfine-coupling fluctuations of the 15 dominant FMN HFC entities for replica 1 of *As*LOV2. The instantaneous fluctuation amplitude is defined as $\|\mathbf{A}(t) - \langle\mathbf{A}\rangle\|_{\mathrm{F}}/\sqrt{3}$, where $\langle\mathbf{A}\rangle$ is the mean HFC tensor over the 500 ns trajectory. Equivalent methyl-proton groups are averaged for visualisation only.

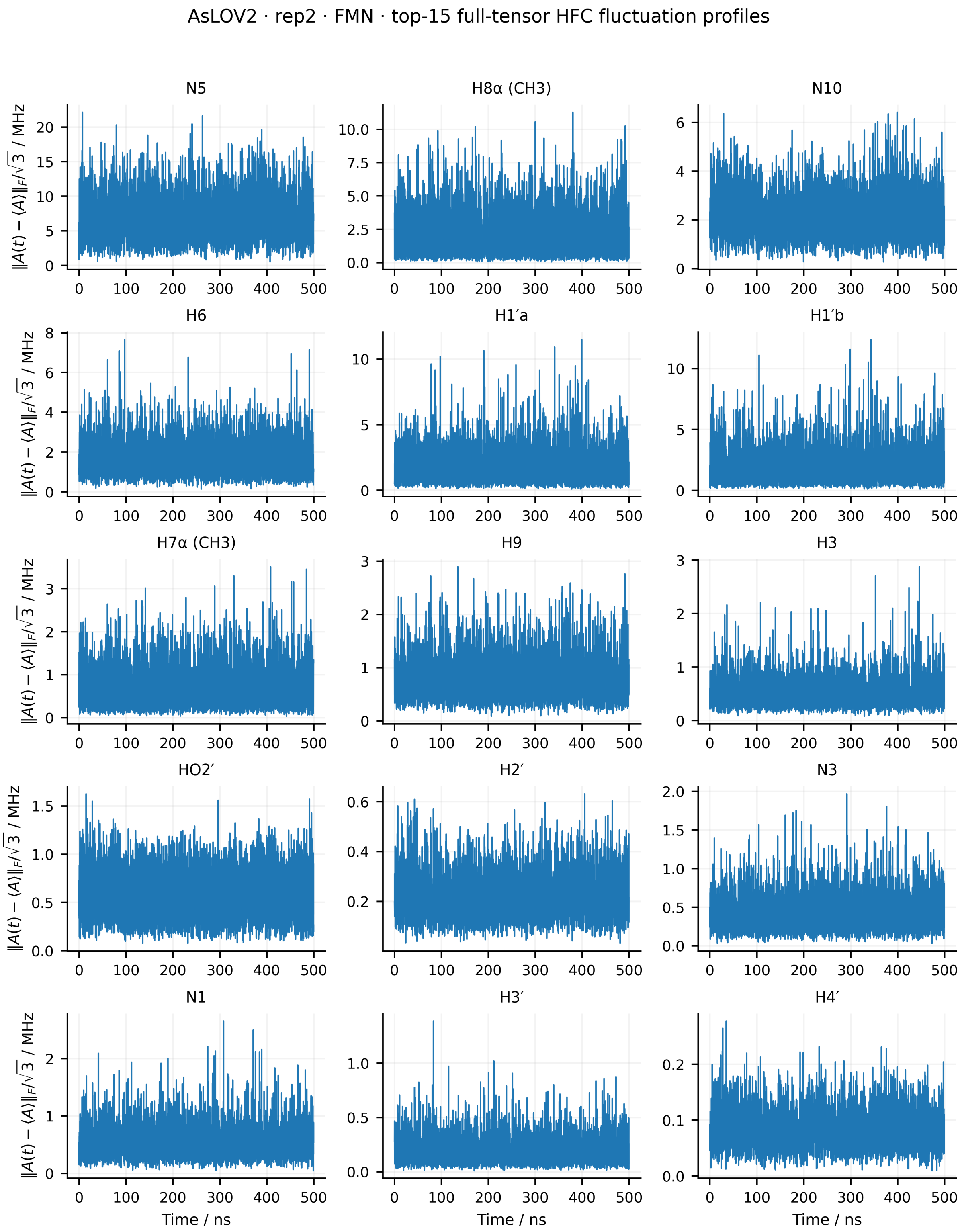


Figure S39: Full-tensor hyperfine-coupling fluctuations of the 15 dominant FMN HFC entities for replica 2 of *As*LOV2. The instantaneous fluctuation amplitude is defined as $\|\mathbf{A}(t) - \langle\mathbf{A}\rangle\|_{\mathrm{F}}/\sqrt{3}$, where $\langle\mathbf{A}\rangle$ is the mean HFC tensor over the 500 ns trajectory. Equivalent methyl-proton groups are averaged for visualisation only.

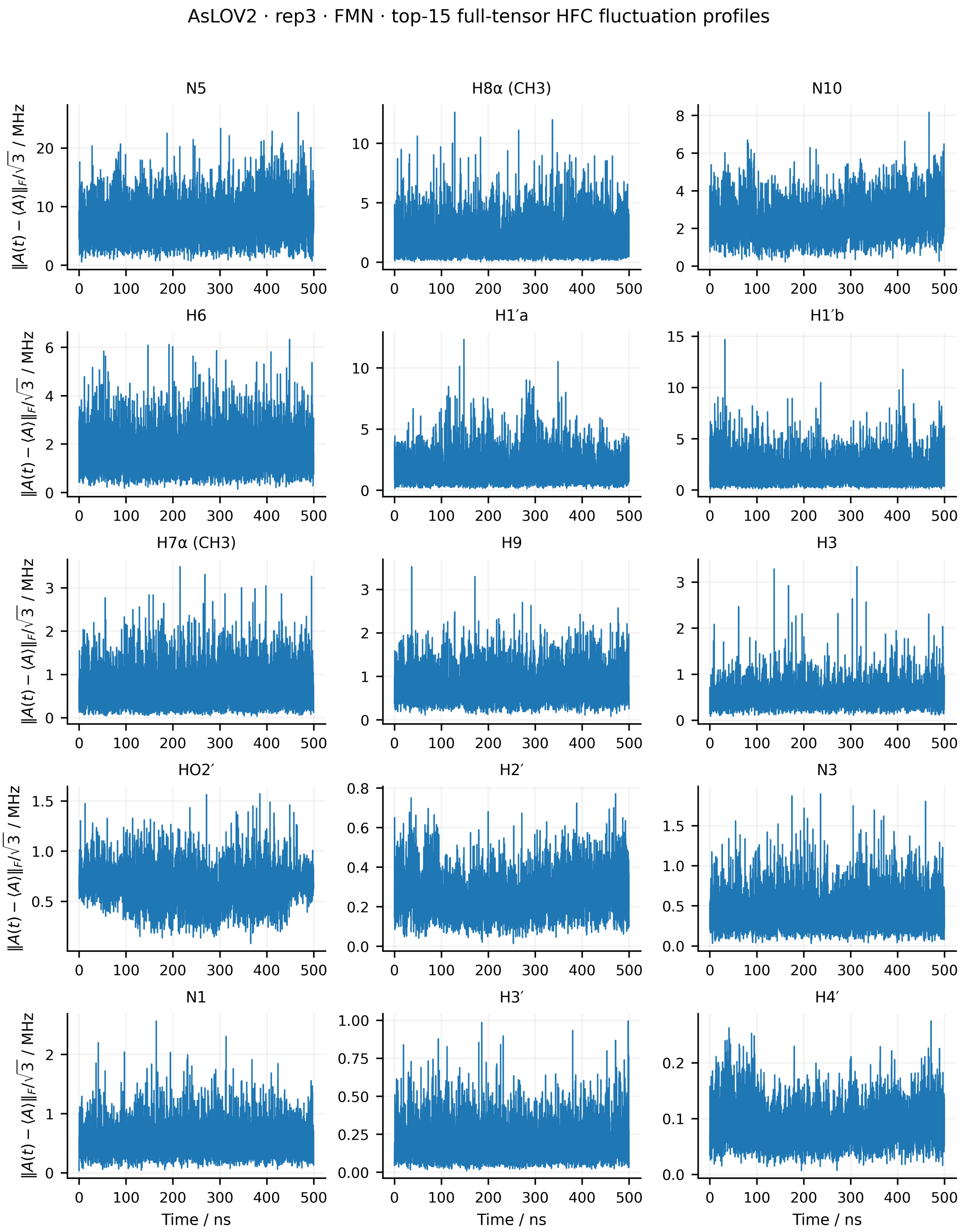


Figure S40: Full-tensor hyperfine-coupling fluctuations of the 15 dominant FMN HFC entities for replica 3 of *As*LOV2. The instantaneous fluctuation amplitude is defined as $\|\mathbf{A}(t) - \langle\mathbf{A}\rangle\|_{\mathrm{F}}/\sqrt{3}$, where $\langle\mathbf{A}\rangle$ is the mean HFC tensor over the 500 ns trajectory. Equivalent methyl-proton groups are averaged for visualisation only.

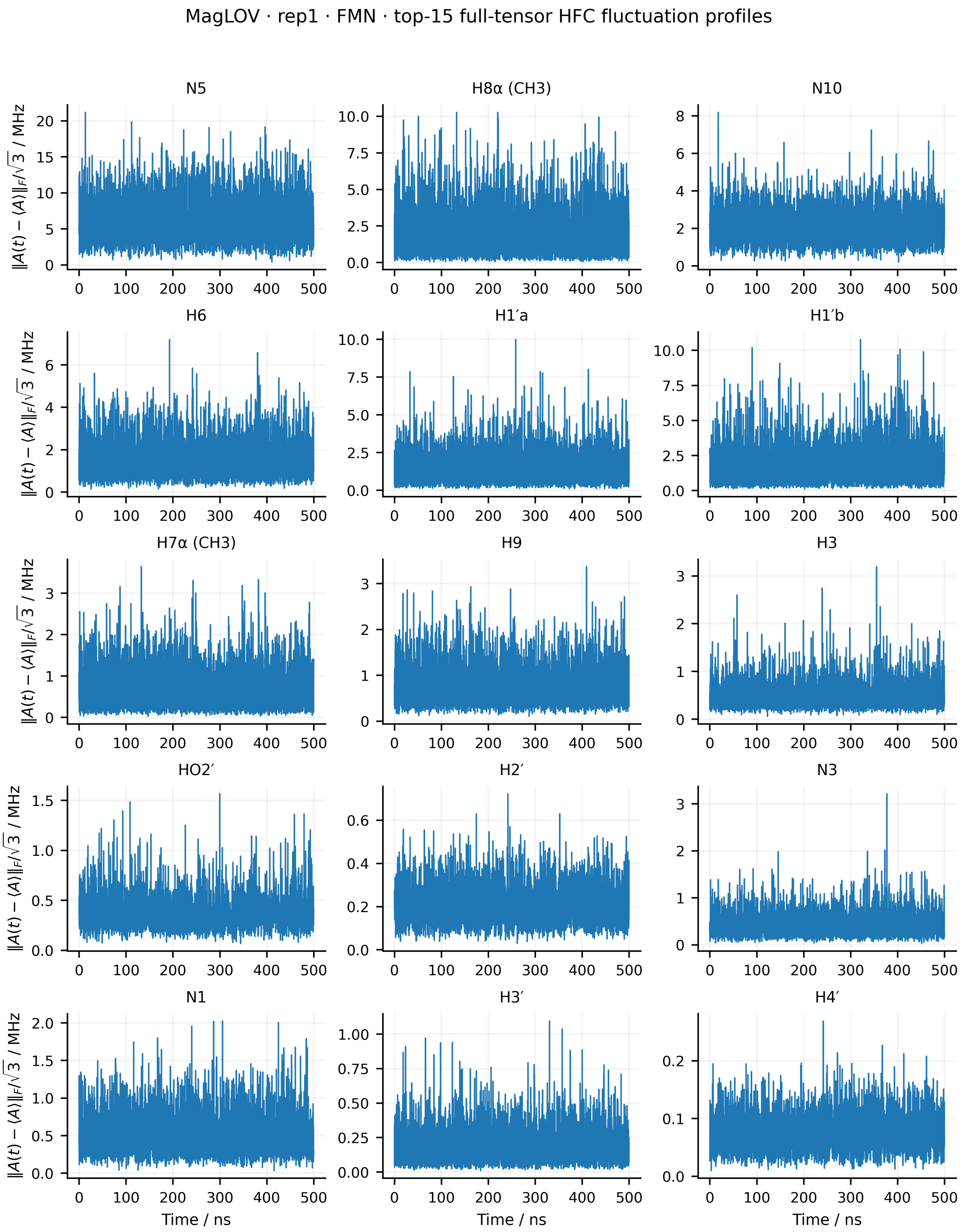


Figure S41: Full-tensor hyperfine-coupling fluctuations of the 15 dominant FMN HFC entities for replica 1 of MagLOV. The instantaneous fluctuation amplitude is defined as $\|\mathbf{A}(t) - \langle \mathbf{A} \rangle\|_{\mathrm{F}}/\sqrt{3}$, where $\langle \mathbf{A} \rangle$ is the mean HFC tensor over the 500 ns trajectory. Equivalent methyl-proton groups are averaged for visualisation only.

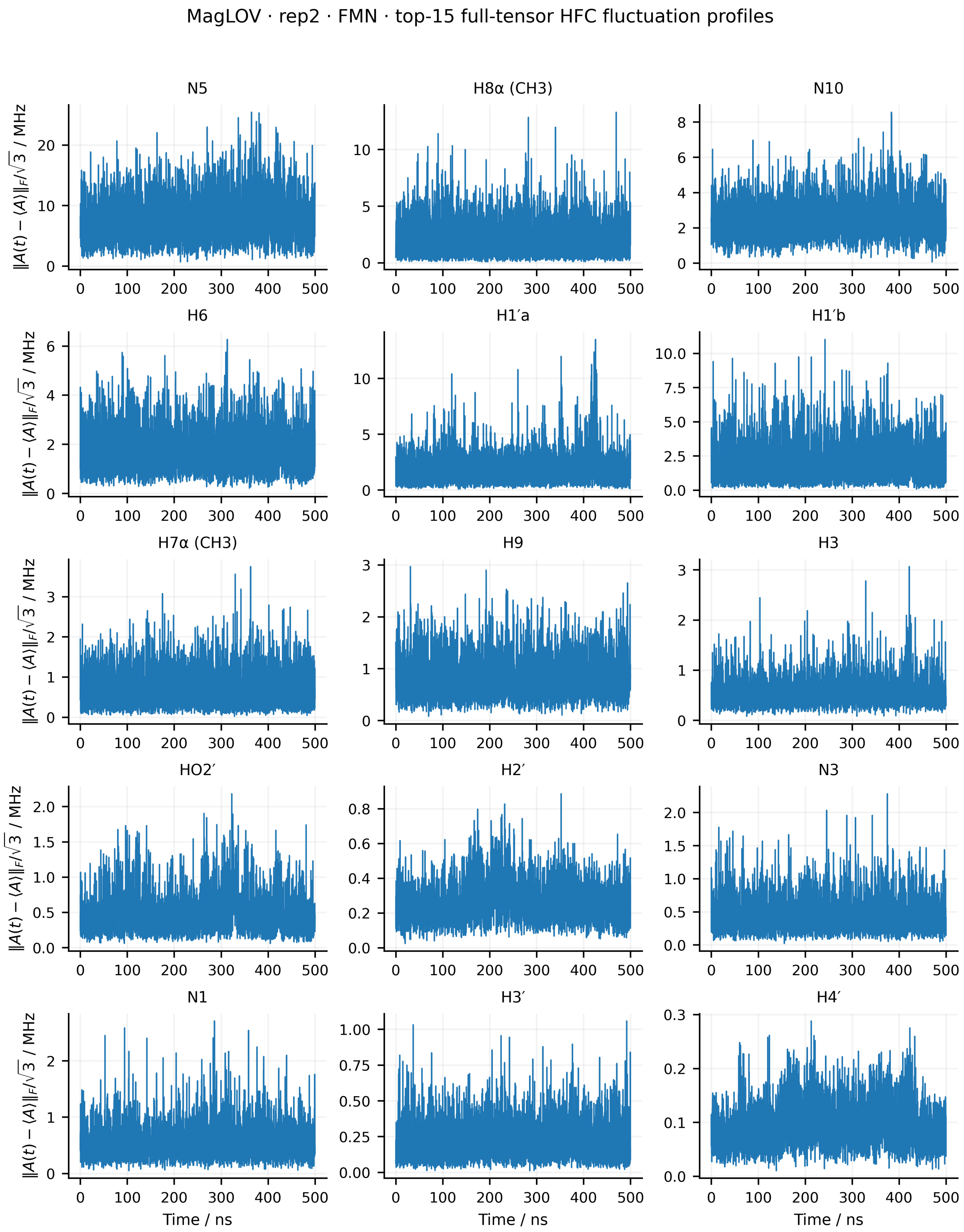


Figure S42: Full-tensor hyperfine-coupling fluctuations of the 15 dominant FMN HFC entities for replica 2 of MagLOV. The instantaneous fluctuation amplitude is defined as $\|\mathbf{A}(t) - \langle \mathbf{A} \rangle\|_{\mathrm{F}}/\sqrt{3}$, where $\langle \mathbf{A} \rangle$ is the mean HFC tensor over the 500 ns trajectory. Equivalent methyl-proton groups are averaged for visualisation only.

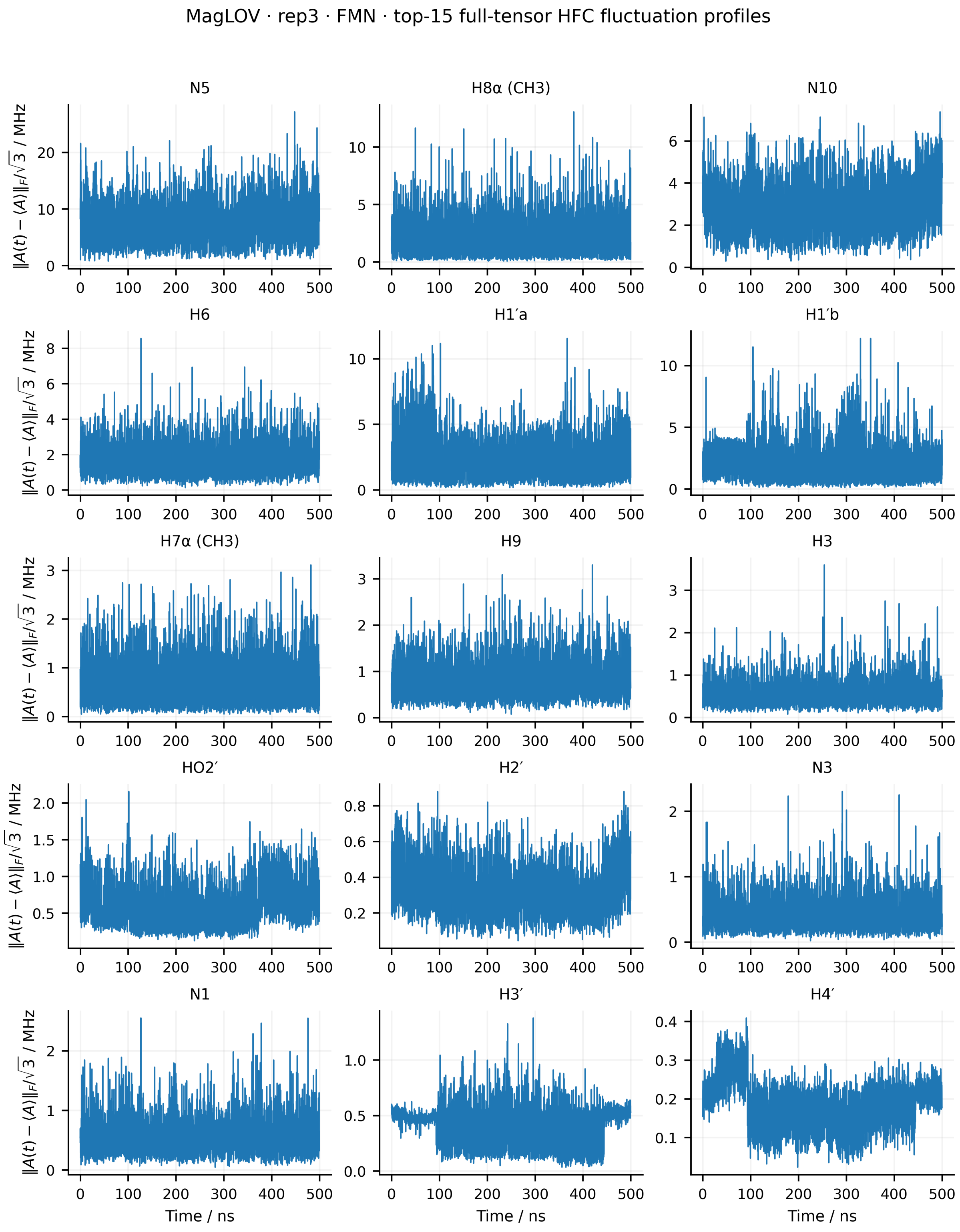


Figure S43: Full-tensor hyperfine-coupling fluctuations of the 15 dominant FMN HFC entities for replica 3 of MagLOV. The instantaneous fluctuation amplitude is defined as $\|\mathbf{A}(t)-\langle\mathbf{A}\rangle\|_{\mathrm{F}}/\sqrt{3}$, where $\langle\mathbf{A}\rangle$ is the mean HFC tensor over the 500 ns trajectory. Equivalent methyl-proton groups are averaged for visualisation only.

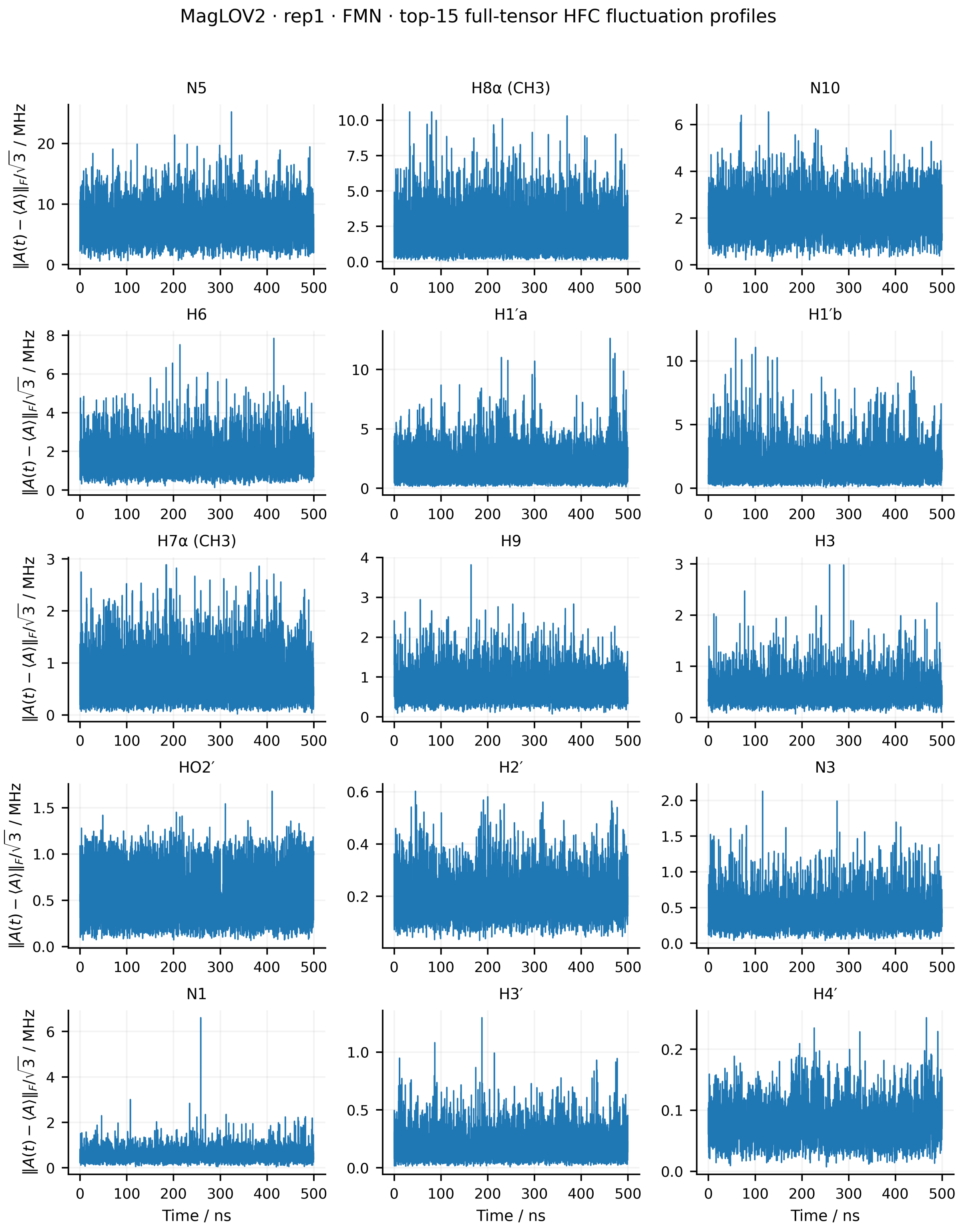


Figure S44: Full-tensor hyperfine-coupling fluctuations of the 15 dominant FMN HFC entities for replica 1 of MagLOV2. The instantaneous fluctuation amplitude is defined as $\|\mathbf{A}(t) - \langle\mathbf{A}\rangle\|_{\mathrm{F}}/\sqrt{3}$, where $\langle\mathbf{A}\rangle$ is the mean HFC tensor over the 500 ns trajectory. Equivalent methyl-proton groups are averaged for visualisation only.

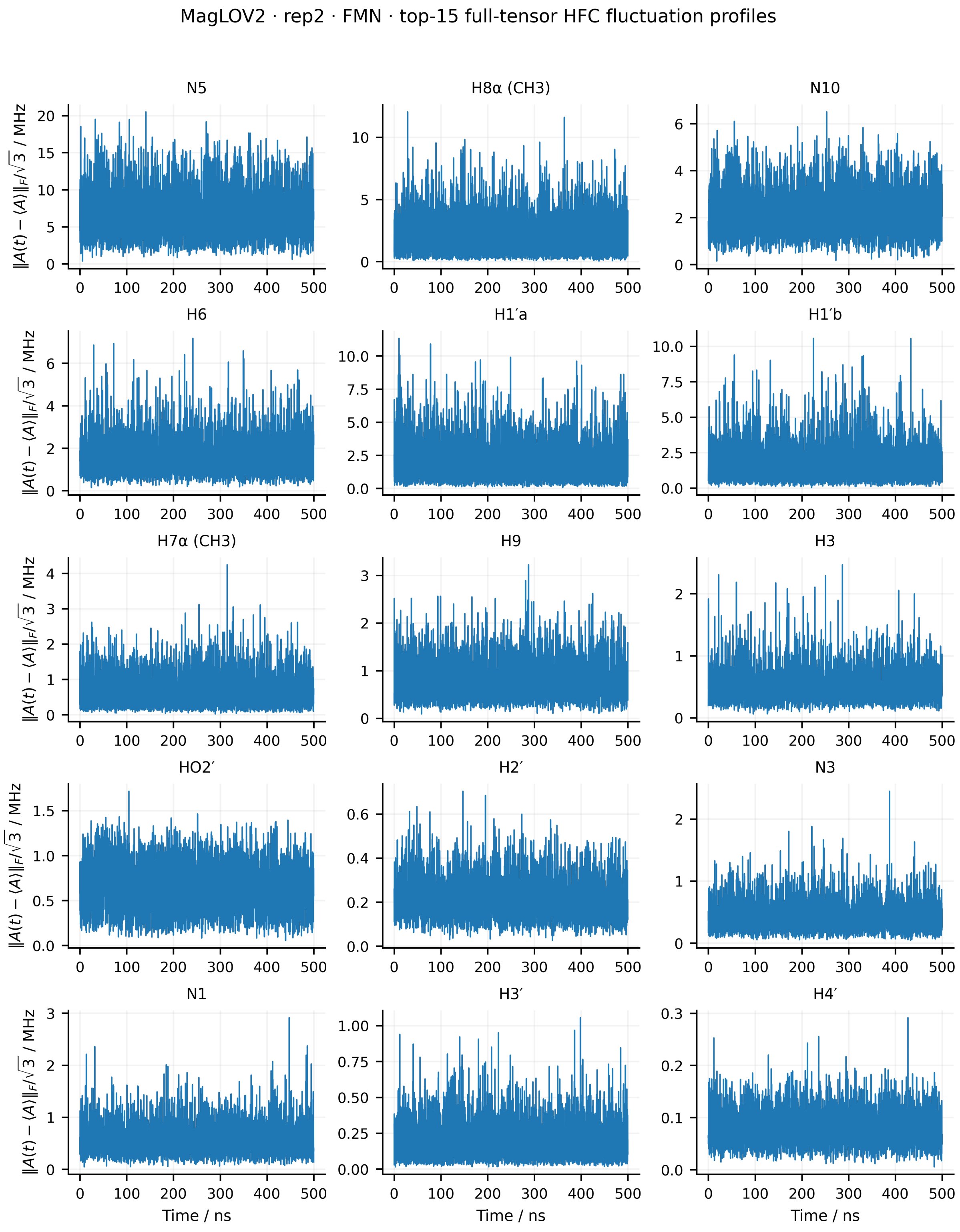


Figure S45: Full-tensor hyperfine-coupling fluctuations of the 15 dominant FMN HFC entities for replica 2 of MagLOV2. The instantaneous fluctuation amplitude is defined as $\|\mathbf{A}(t) - \langle \mathbf{A} \rangle\|_\mathrm{F}/\sqrt{3}$, where $\langle \mathbf{A} \rangle$ is the mean HFC tensor over the 500 ns trajectory. Equivalent methyl-proton groups are averaged for visualisation only.

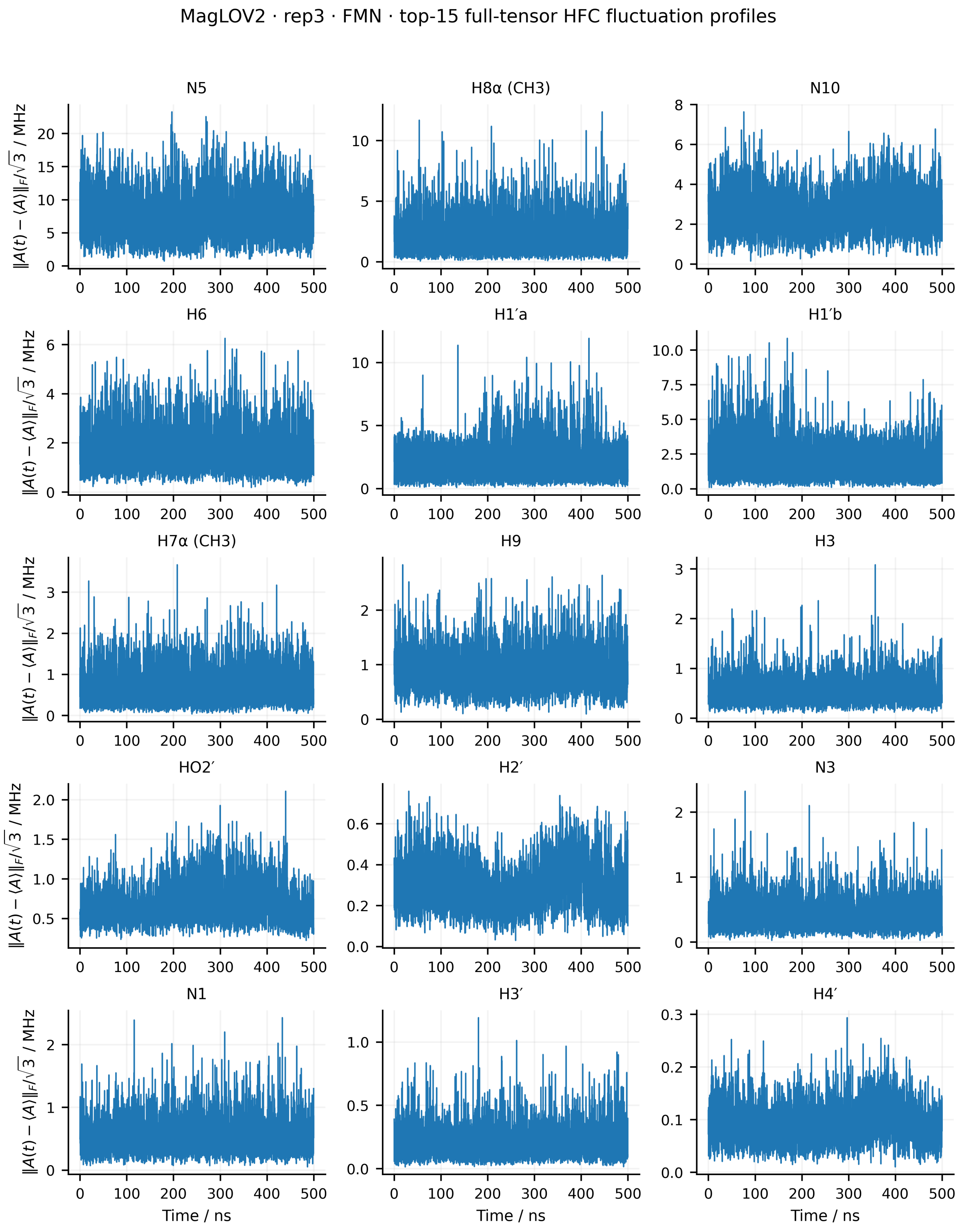


Figure S46: Full-tensor hyperfine-coupling fluctuations of the 15 dominant FMN HFC entities for replica 3 of MagLOV2. The instantaneous fluctuation amplitude is defined as $\|\mathbf{A}(t)-\langle\mathbf{A}\rangle\|_{\mathrm{F}}/\sqrt{3}$, where $\langle\mathbf{A}\rangle$ is the mean HFC tensor over the 500 ns trajectory. Equivalent methyl-proton groups are averaged for visualisation only.

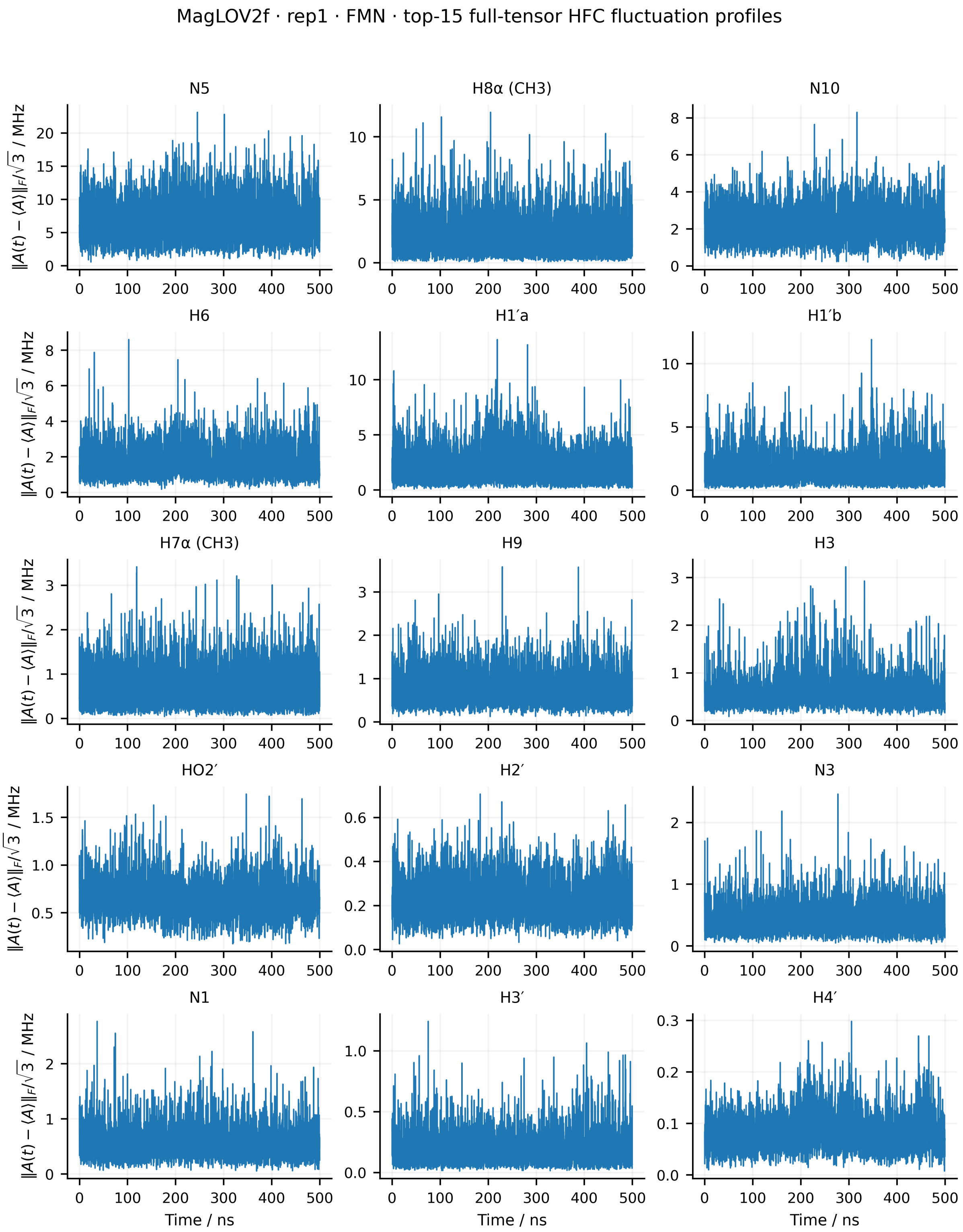


Figure S47: Full-tensor hyperfine-coupling fluctuations of the 15 dominant FMN HFC entities for replica 1 of MagLOV2f. The instantaneous fluctuation amplitude is defined as $\|\mathbf{A}(t) - \langle \mathbf{A} \rangle\|_F / \sqrt{3}$, where $\langle \mathbf{A} \rangle$ is the mean HFC tensor over the 500 ns trajectory. Equivalent methyl-proton groups are averaged for visualisation only.

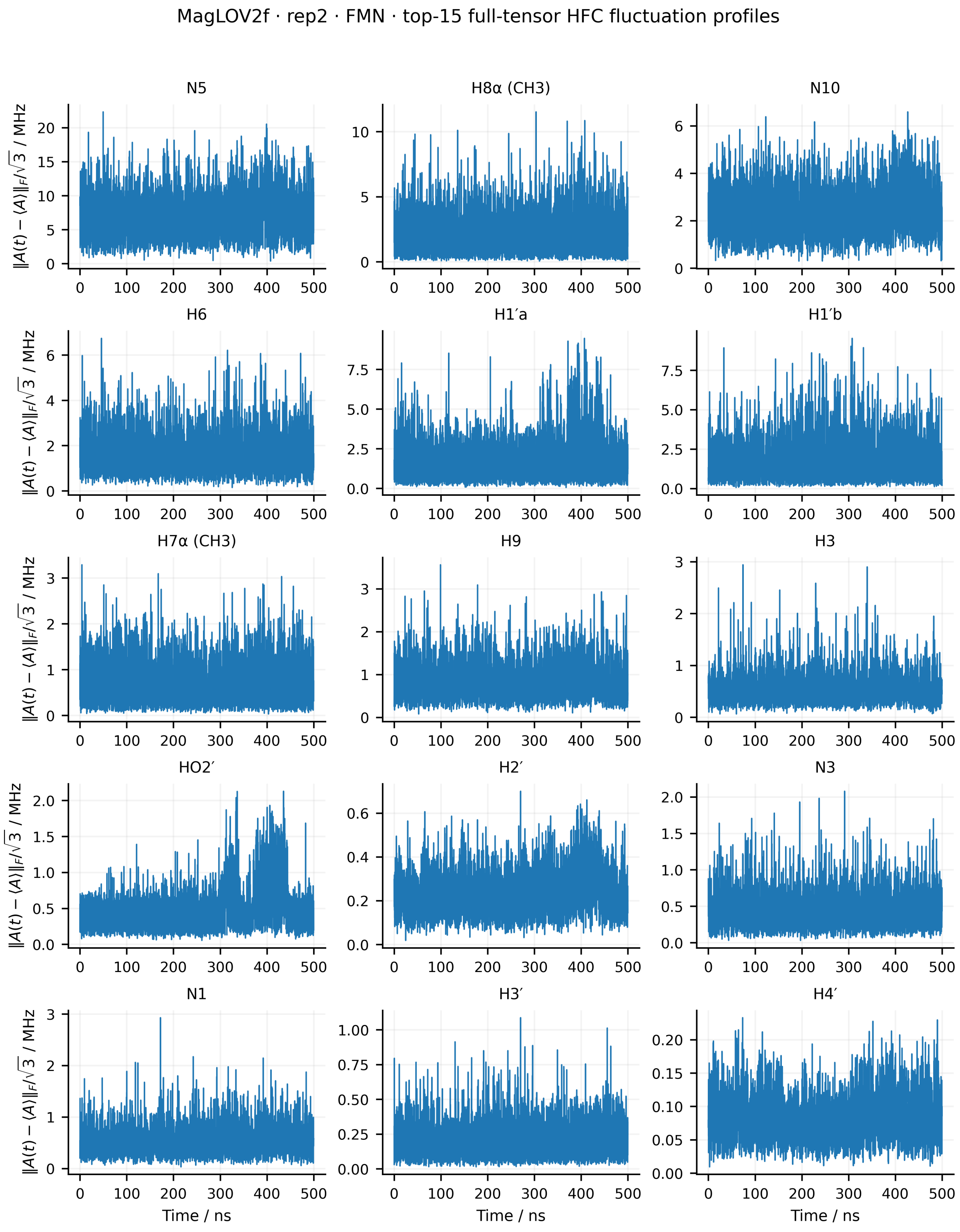


Figure S48: Full-tensor hyperfine-coupling fluctuations of the 15 dominant FMN HFC entities for replica 2 of MagLOV2f. The instantaneous fluctuation amplitude is defined as $\|\mathbf{A}(t)-\langle\mathbf{A}\rangle\|_{\mathrm{F}}/\sqrt{3}$, where $\langle\mathbf{A}\rangle$ is the mean HFC tensor over the 500 ns trajectory. Equivalent methyl-proton groups are averaged for visualisation only.

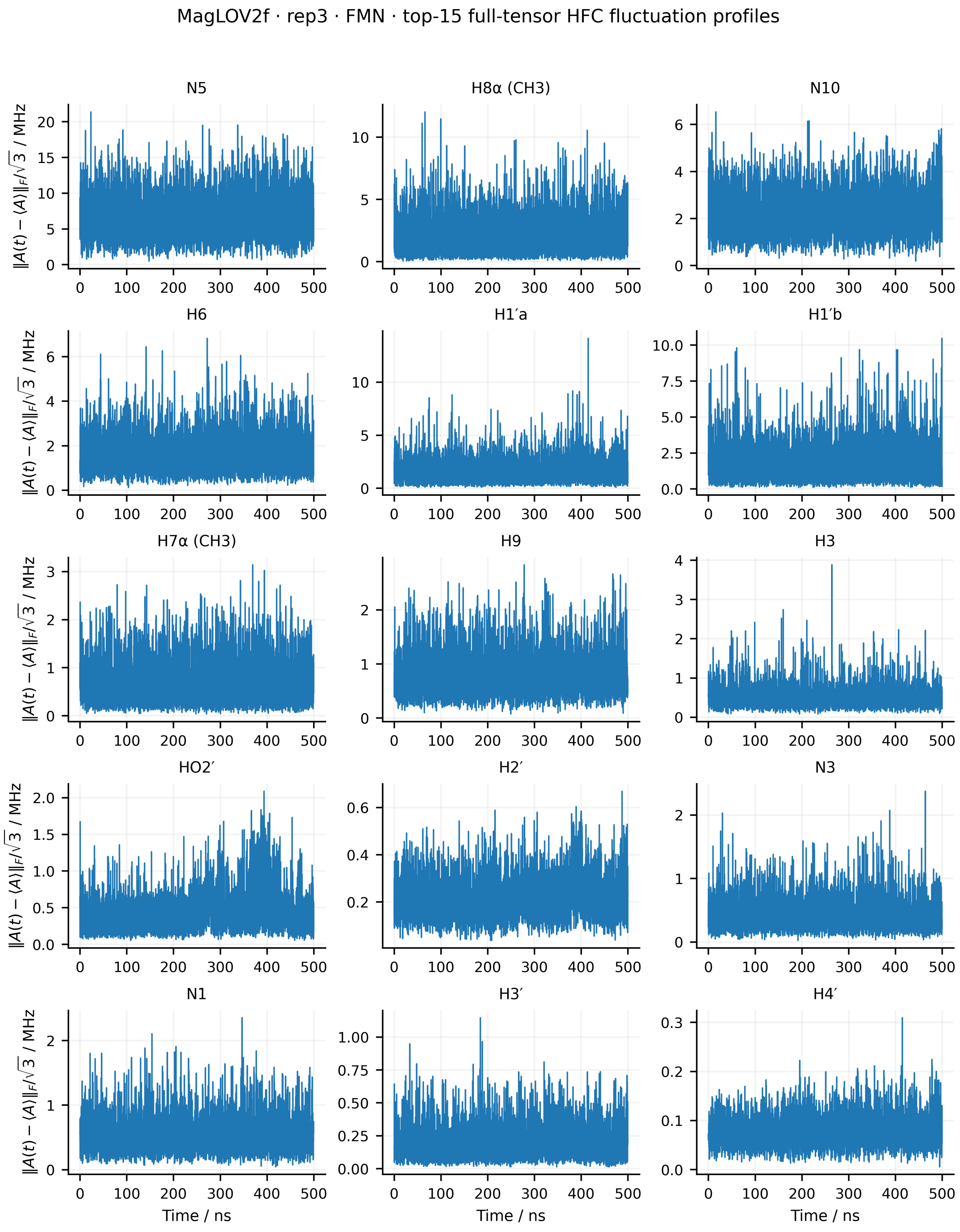


Figure S49: Full-tensor hyperfine-coupling fluctuations of the 15 dominant FMN HFC entities for replica 3 of MagLOV2f. The instantaneous fluctuation amplitude is defined as $\|\mathbf{A}(t)-\langle\mathbf{A}\rangle\|_{\mathrm{F}}/\sqrt{3}$, where $\langle\mathbf{A}\rangle$ is the mean HFC tensor over the 500 ns trajectory. Equivalent methyl-proton groups are averaged for visualisation only.

##### S8.2.2 Trp hyperfine coupling modulation

The corresponding analysis was performed for the $TrpH^{\bullet +}$ radical. In contrast to FMN, the Trp side chain is directly embedded in the mutation-dependent donor environment, such that changes in side-chain conformation and local protein interactions can strongly modulate its HFC tensors. The same physical Trp HFC entities are compared across all variants and replicas. Artificial hydrogen atoms introduced to cap the truncated quantum-chemical fragment were excluded from the analysis.

**Isotropic hyperfine fluctuations** The time-dependent isotropic HFCs of the 11 physical Trp HFC entities are shown below. As for FMN, $A_{\text{iso}}(t)$ was obtained from one third of the tensor trace, and the dashed horizontal line denotes the mean over the complete 500 ns trajectory. The profiles provide a direct view of both rapid HFC fluctuations and slower conformational changes that shift the local magnetic environment over extended trajectory intervals.

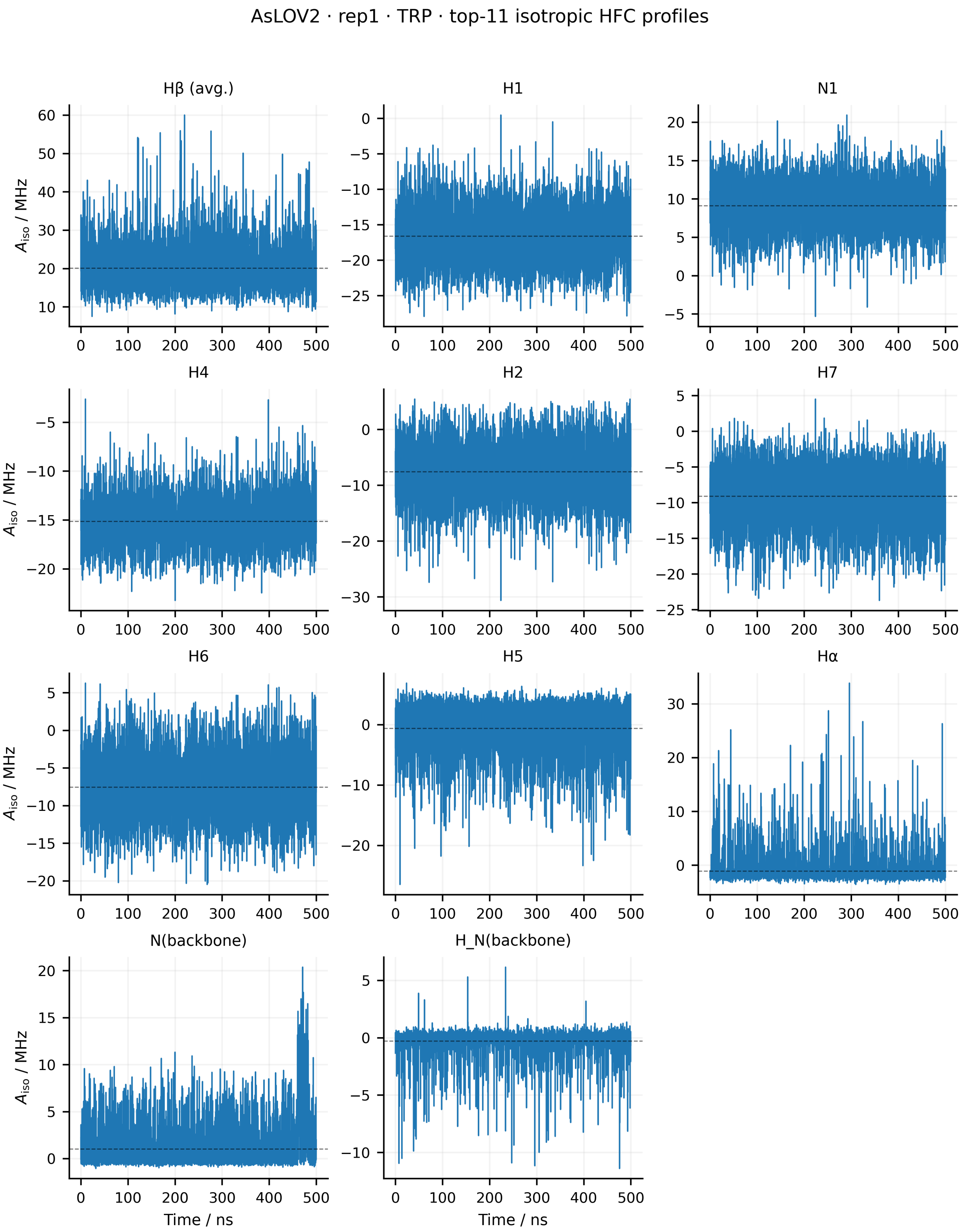


Figure S50: Time evolution of the isotropic hyperfine couplings of the 11 physical Trp HFC entities for replica 1 of *As*LOV2. The isotropic coupling was calculated as $A_{\mathrm{iso}}(t) = \mathrm{Tr}[\mathbf{A}(t)]/3$. Dashed horizontal lines indicate the mean value over the 500 ns trajectory. The two H$\beta$ couplings are averaged for visualisation, while the individual nuclear spins are retained in the relaxation analysis. Artificial capping hydrogens are excluded.

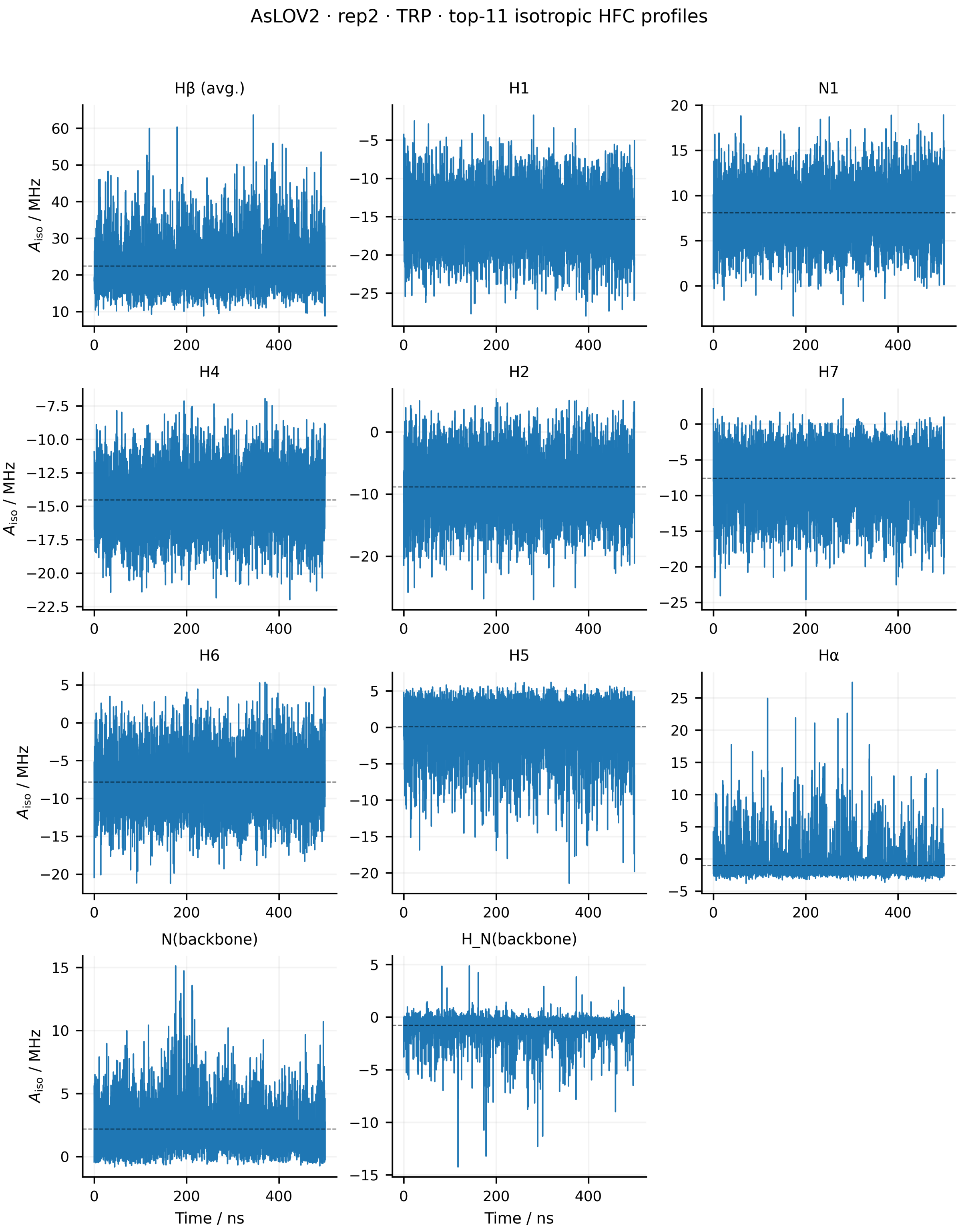


Figure S51: Time evolution of the isotropic hyperfine couplings of the 11 physical Trp HFC entities for replica 2 of *As*LOV2. The isotropic coupling was calculated as $A_{\mathrm{iso}}(t) = \mathrm{Tr}[\mathbf{A}(t)]/3$. Dashed horizontal lines indicate the mean value over the 500 ns trajectory. The two H$\beta$ couplings are averaged for visualisation, while the individual nuclear spins are retained in the relaxation analysis. Artificial capping hydrogens are excluded.

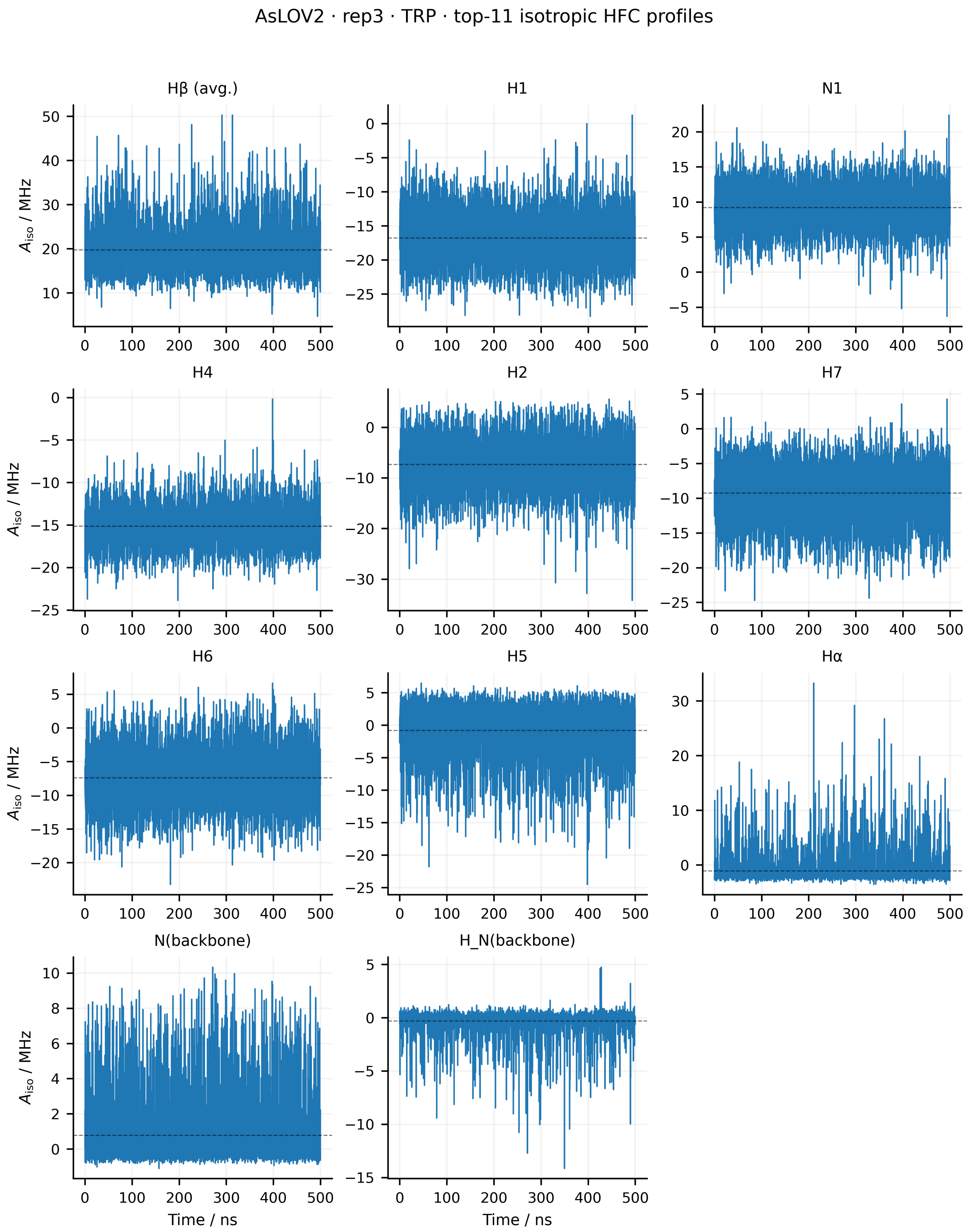


Figure S52: Time evolution of the isotropic hyperfine couplings of the 11 physical Trp HFC entities for replica 3 of *As*LOV2. The isotropic coupling was calculated as $A_{\rm iso}(t) = {\rm Tr}[\mathbf{A}(t)]/3$. Dashed horizontal lines indicate the mean value over the 500 ns trajectory. The two H$\beta$ couplings are averaged for visualisation, while the individual nuclear spins are retained in the relaxation analysis. Artificial capping hydrogens are excluded.

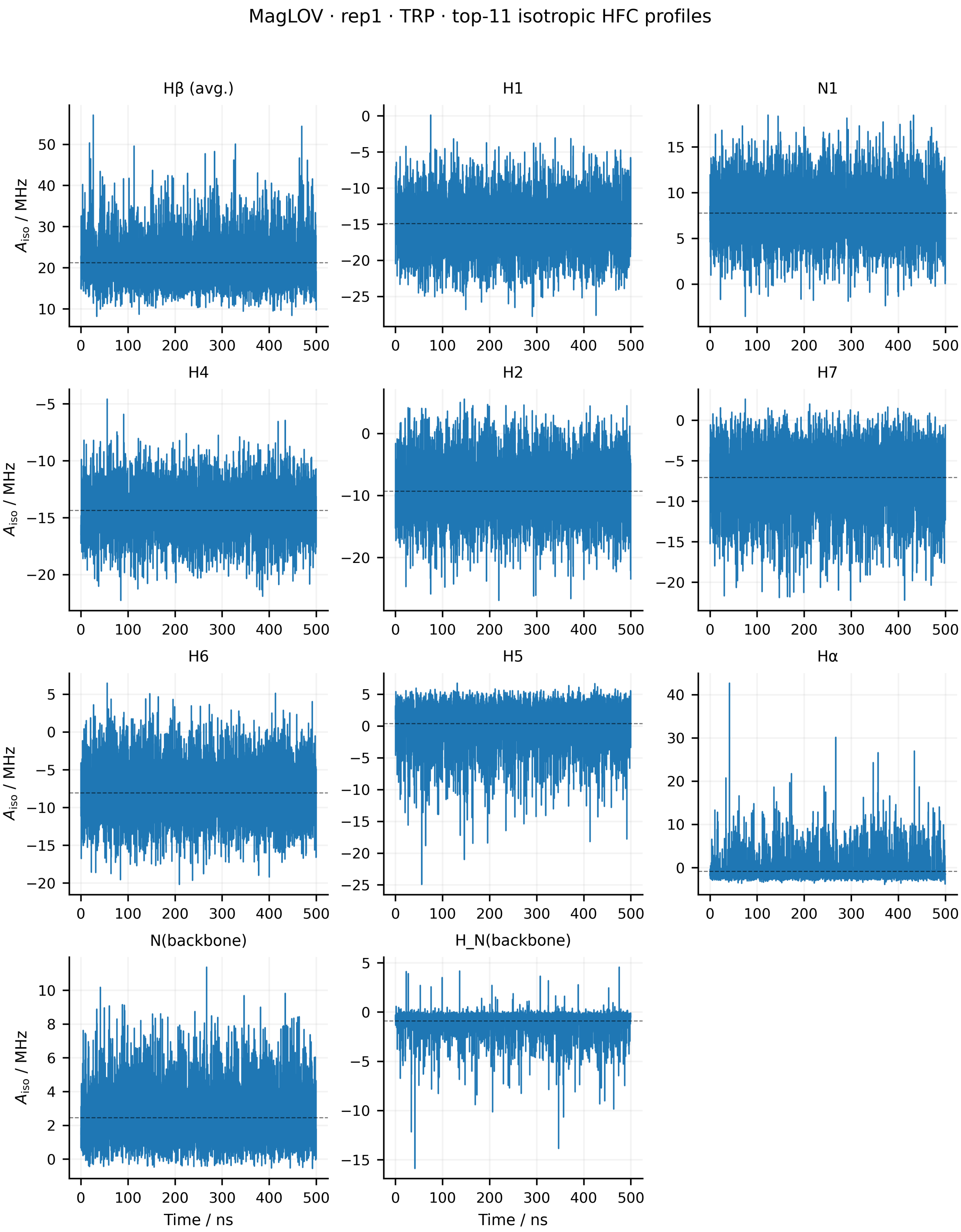


Figure S53: Time evolution of the isotropic hyperfine couplings of the 11 physical Trp HFC entities for replica 1 of MagLOV. The isotropic coupling was calculated as $A_{\text{iso}}(t) = \text{Tr}[\mathbf{A}(t)]/3$. Dashed horizontal lines indicate the mean value over the 500 ns trajectory. The two H$\beta$ couplings are averaged for visualisation, while the individual nuclear spins are retained in the relaxation analysis. Artificial capping hydrogens are excluded.

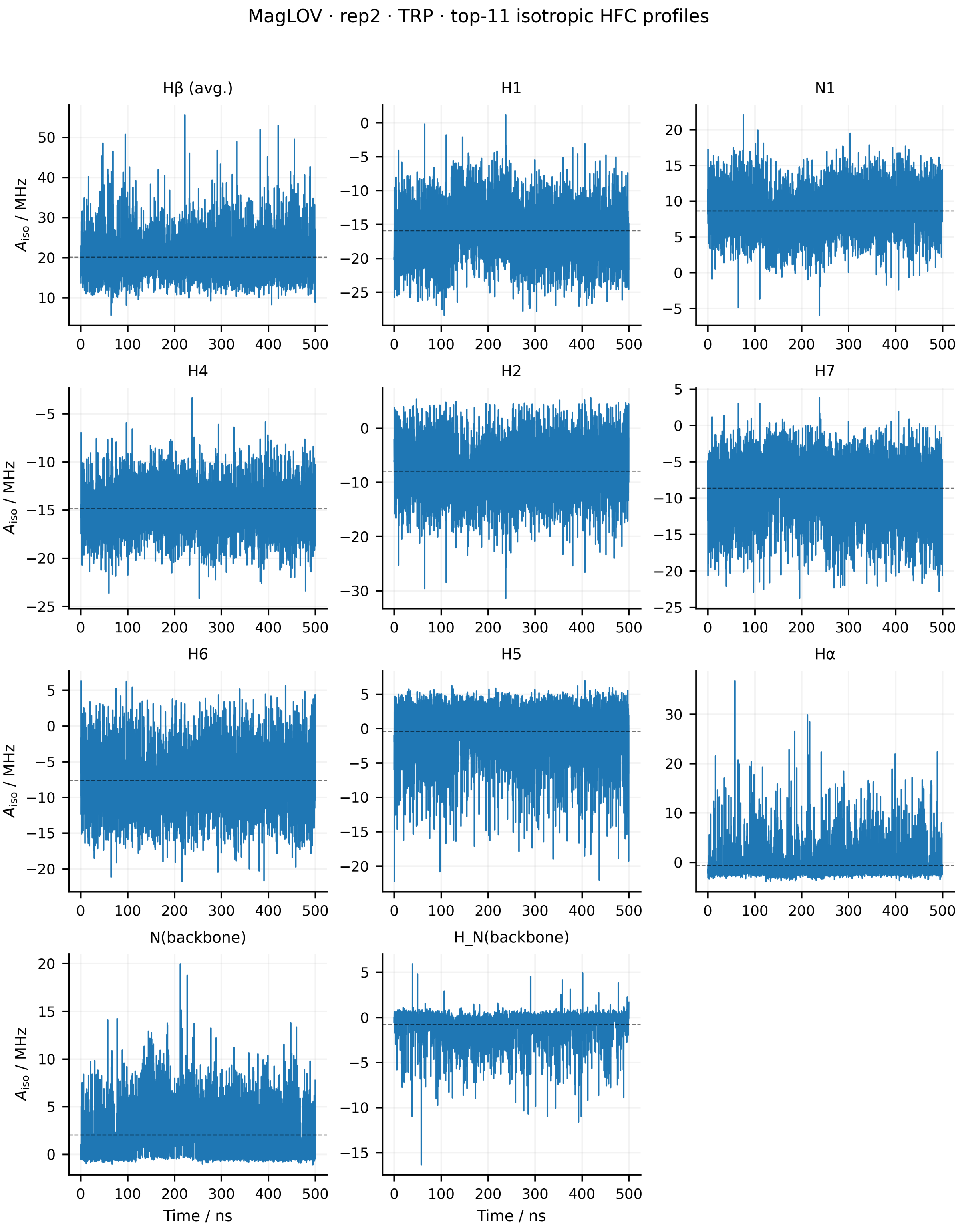


Figure S54: Time evolution of the isotropic hyperfine couplings of the 11 physical Trp HFC entities for replica 2 of MagLOV. The isotropic coupling was calculated as $A_{\text{iso}}(t) = \text{Tr}[\mathbf{A}(t)]/3$. Dashed horizontal lines indicate the mean value over the 500 ns trajectory. The two H$\beta$ couplings are averaged for visualisation, while the individual nuclear spins are retained in the relaxation analysis. Artificial capping hydrogens are excluded.

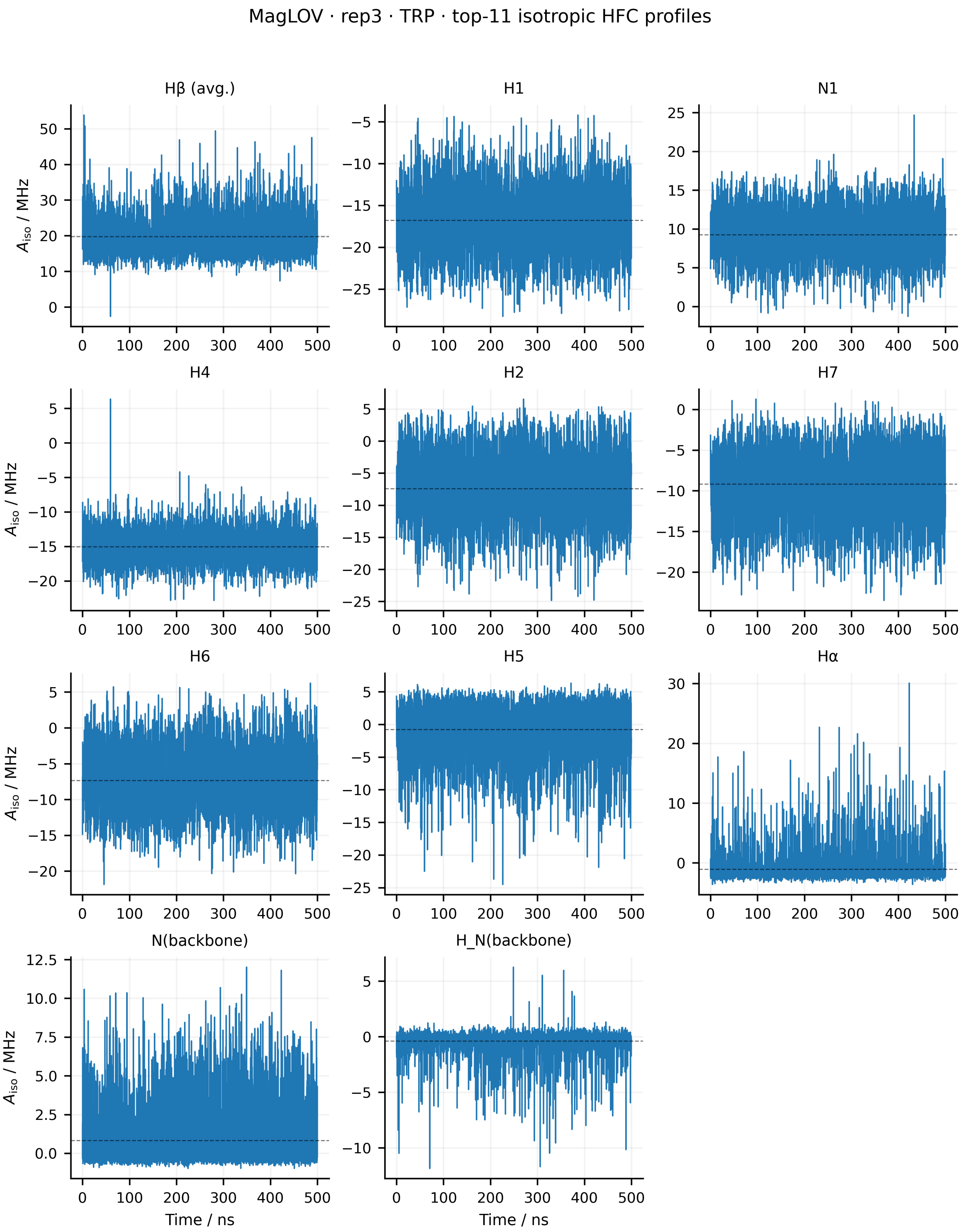


Figure S55: Time evolution of the isotropic hyperfine couplings of the 11 physical Trp HFC entities for replica 3 of MagLOV. The isotropic coupling was calculated as $A_{\text{iso}}(t) = \text{Tr}[\mathbf{A}(t)]/3$. Dashed horizontal lines indicate the mean value over the 500 ns trajectory. The two H$\beta$ couplings are averaged for visualisation, while the individual nuclear spins are retained in the relaxation analysis. Artificial capping hydrogens are excluded.

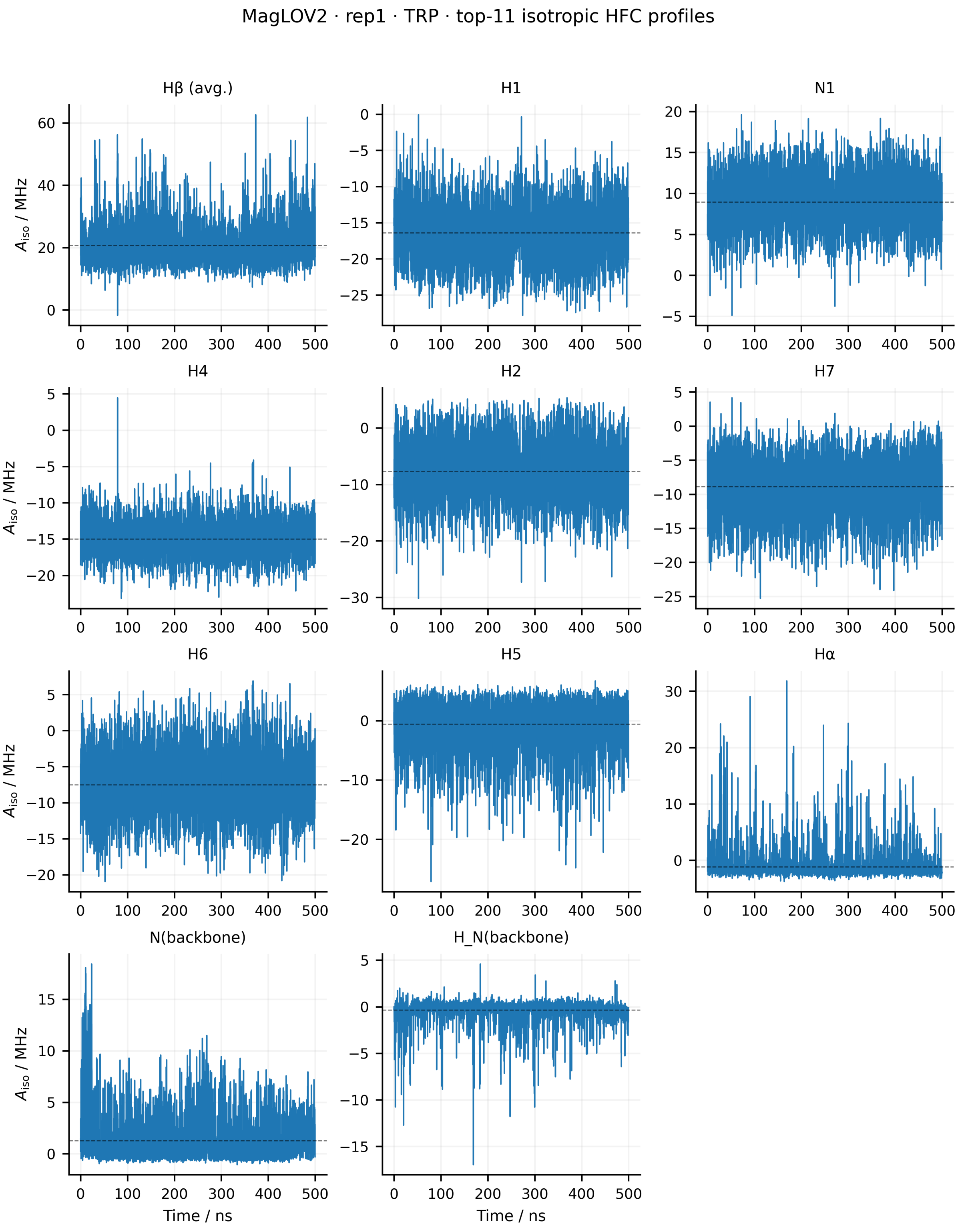


Figure S56: Time evolution of the isotropic hyperfine couplings of the 11 physical Trp HFC entities for replica 1 of MagLOV2. The isotropic coupling was calculated as $A_{\text{iso}}(t) = \text{Tr}[\mathbf{A}(t)]/3$. Dashed horizontal lines indicate the mean value over the 500 ns trajectory. The two H$\beta$ couplings are averaged for visualisation, while the individual nuclear spins are retained in the relaxation analysis. Artificial capping hydrogens are excluded.

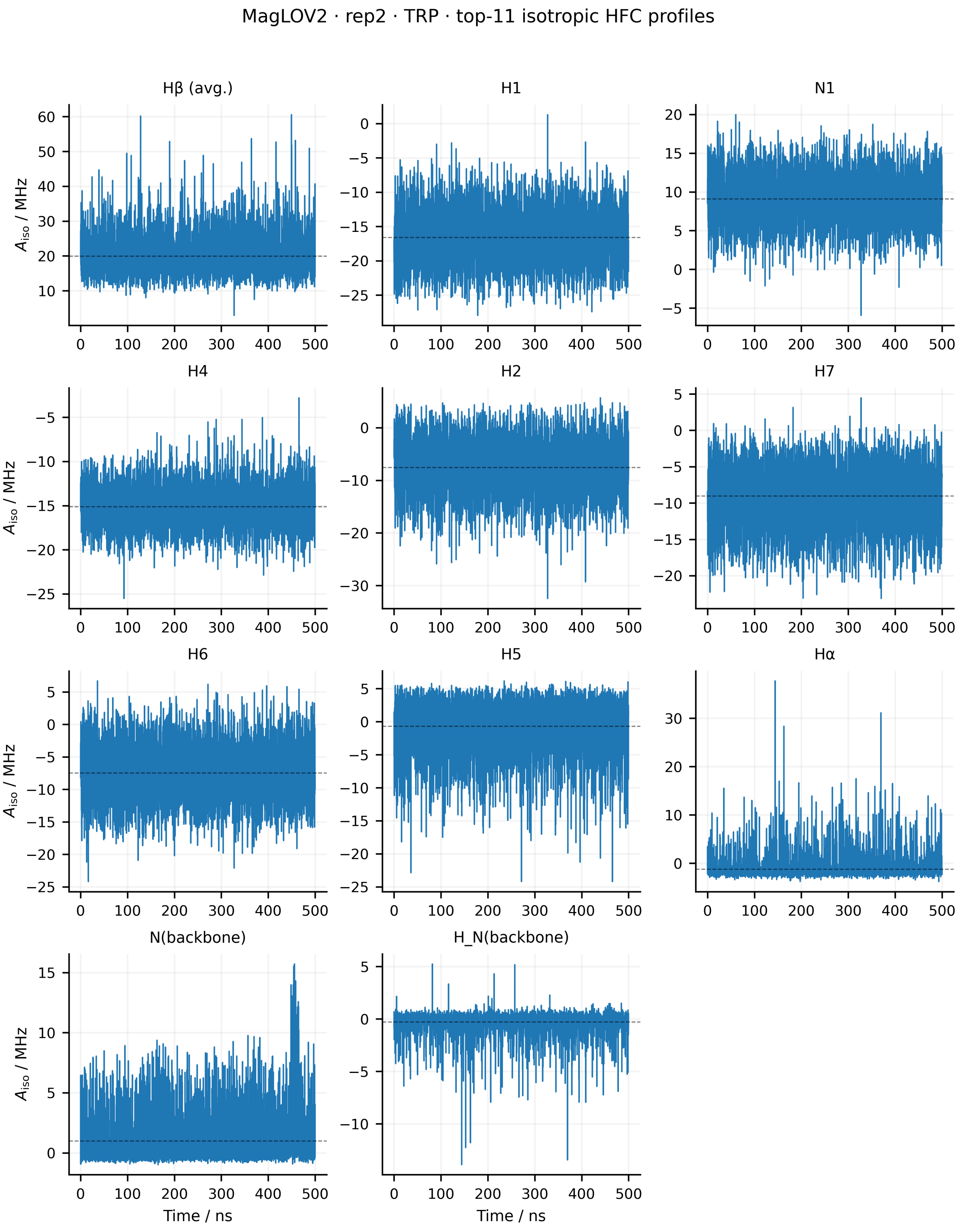


Figure S57: Time evolution of the isotropic hyperfine couplings of the 11 physical Trp HFC entities for replica 2 of MagLOV2. The isotropic coupling was calculated as $A_{\text{iso}}(t) = \text{Tr}[\mathbf{A}(t)]/3$. Dashed horizontal lines indicate the mean value over the 500 ns trajectory. The two H$\beta$ couplings are averaged for visualisation, while the individual nuclear spins are retained in the relaxation analysis. Artificial capping hydrogens are excluded.

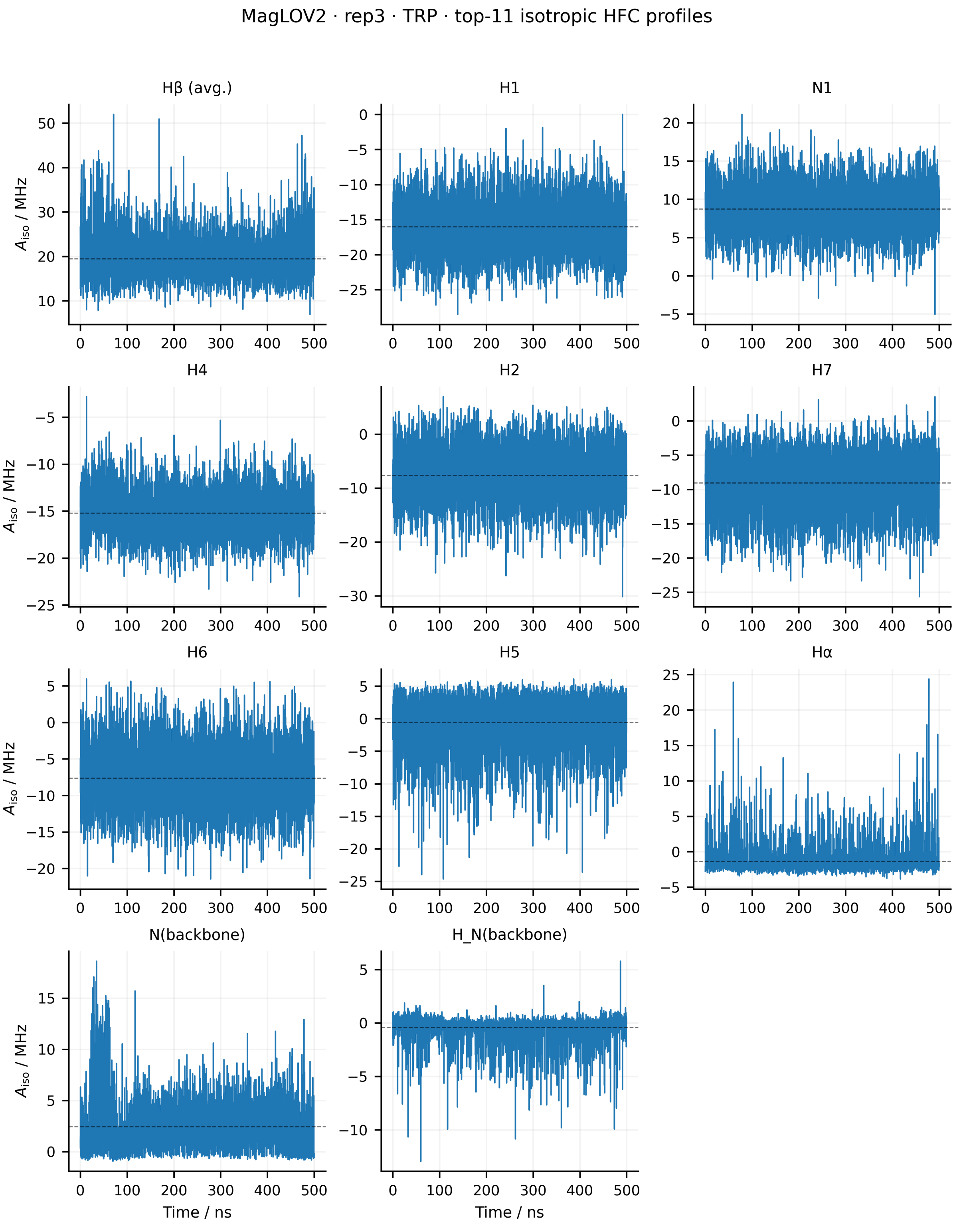


Figure S58: Time evolution of the isotropic hyperfine couplings of the 11 physical Trp HFC entities for replica 3 of MagLOV2. The isotropic coupling was calculated as $A_{\mathrm{iso}}(t) = \mathrm{Tr}[\mathbf{A}(t)]/3$. Dashed horizontal lines indicate the mean value over the 500 ns trajectory. The two H$\beta$ couplings are averaged for visualisation, while the individual nuclear spins are retained in the relaxation analysis. Artificial capping hydrogens are excluded.

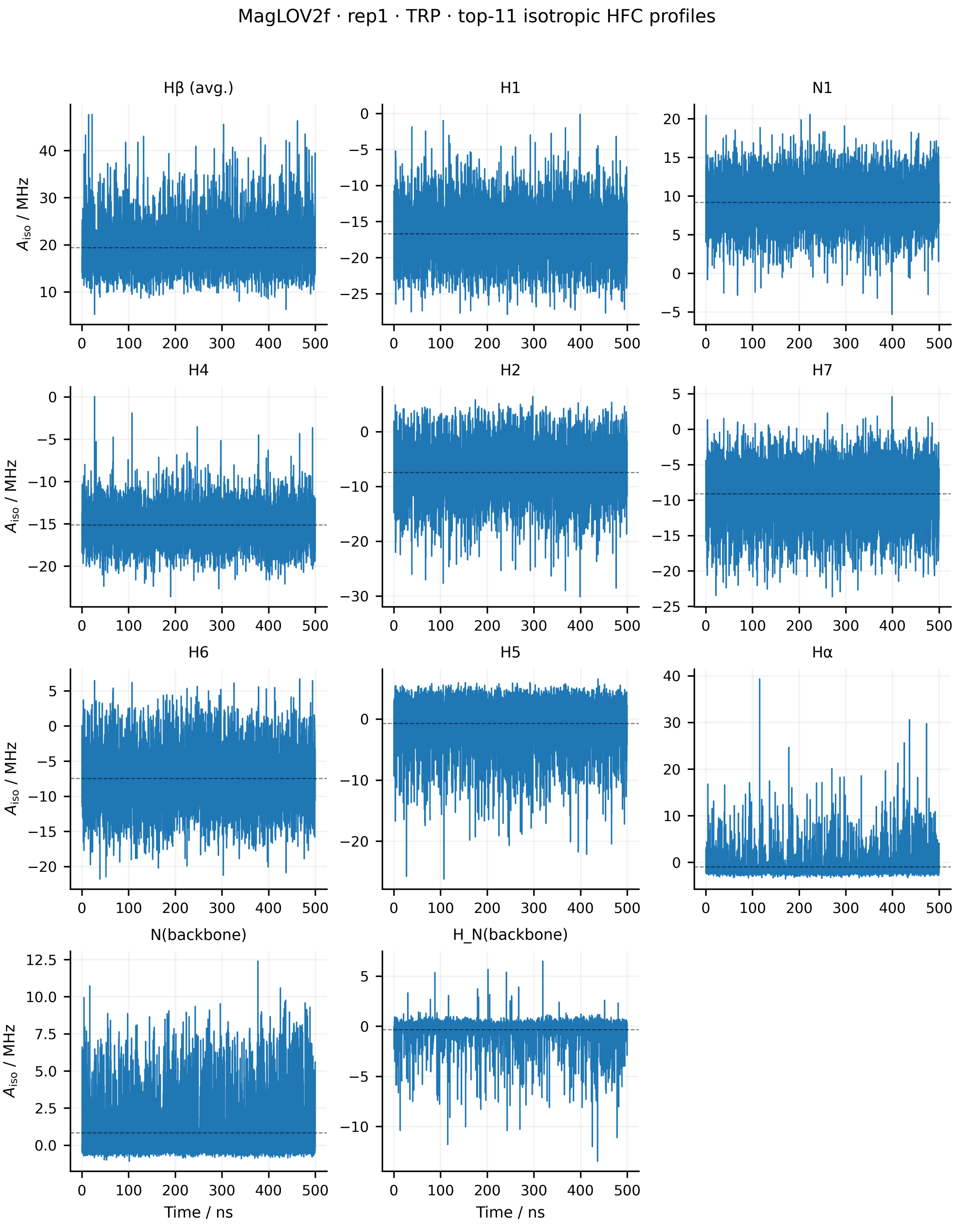


Figure S59: Time evolution of the isotropic hyperfine couplings of the 11 physical Trp HFC entities for replica 1 of MagLOV2f. The isotropic coupling was calculated as $A_{\text{iso}}(t) = \text{Tr}[\mathbf{A}(t)]/3$. Dashed horizontal lines indicate the mean value over the 500 ns trajectory. The two H$\beta$ couplings are averaged for visualisation, while the individual nuclear spins are retained in the relaxation analysis. Artificial capping hydrogens are excluded.

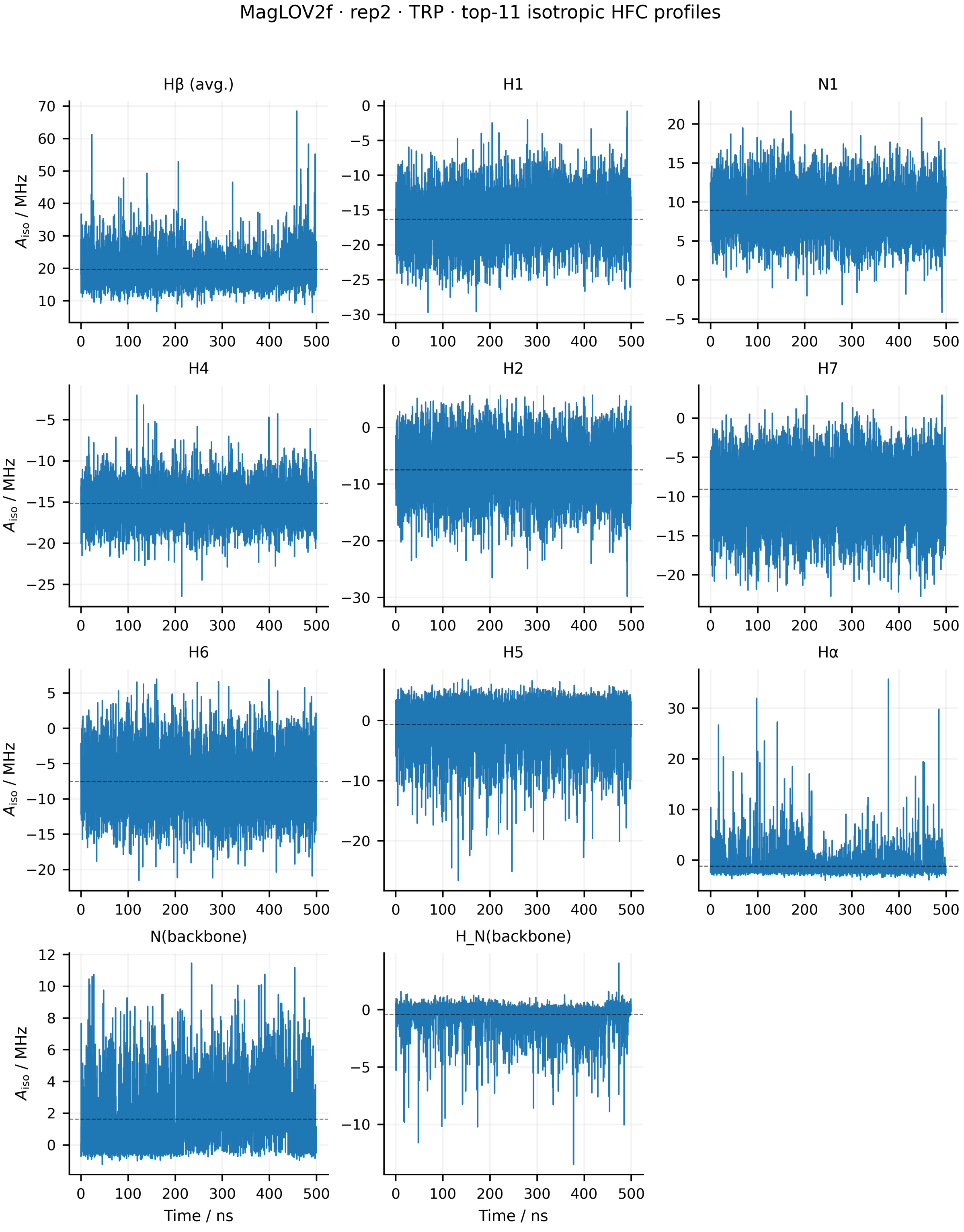


Figure S60: Time evolution of the isotropic hyperfine couplings of the 11 physical Trp HFC entities for replica 2 of MagLOV2f. The isotropic coupling was calculated as $A_{\mathrm{iso}}(t) = \mathrm{Tr}[\mathbf{A}(t)]/3$. Dashed horizontal lines indicate the mean value over the 500 ns trajectory. The two H$\beta$ couplings are averaged for visualisation, while the individual nuclear spins are retained in the relaxation analysis. Artificial capping hydrogens are excluded.

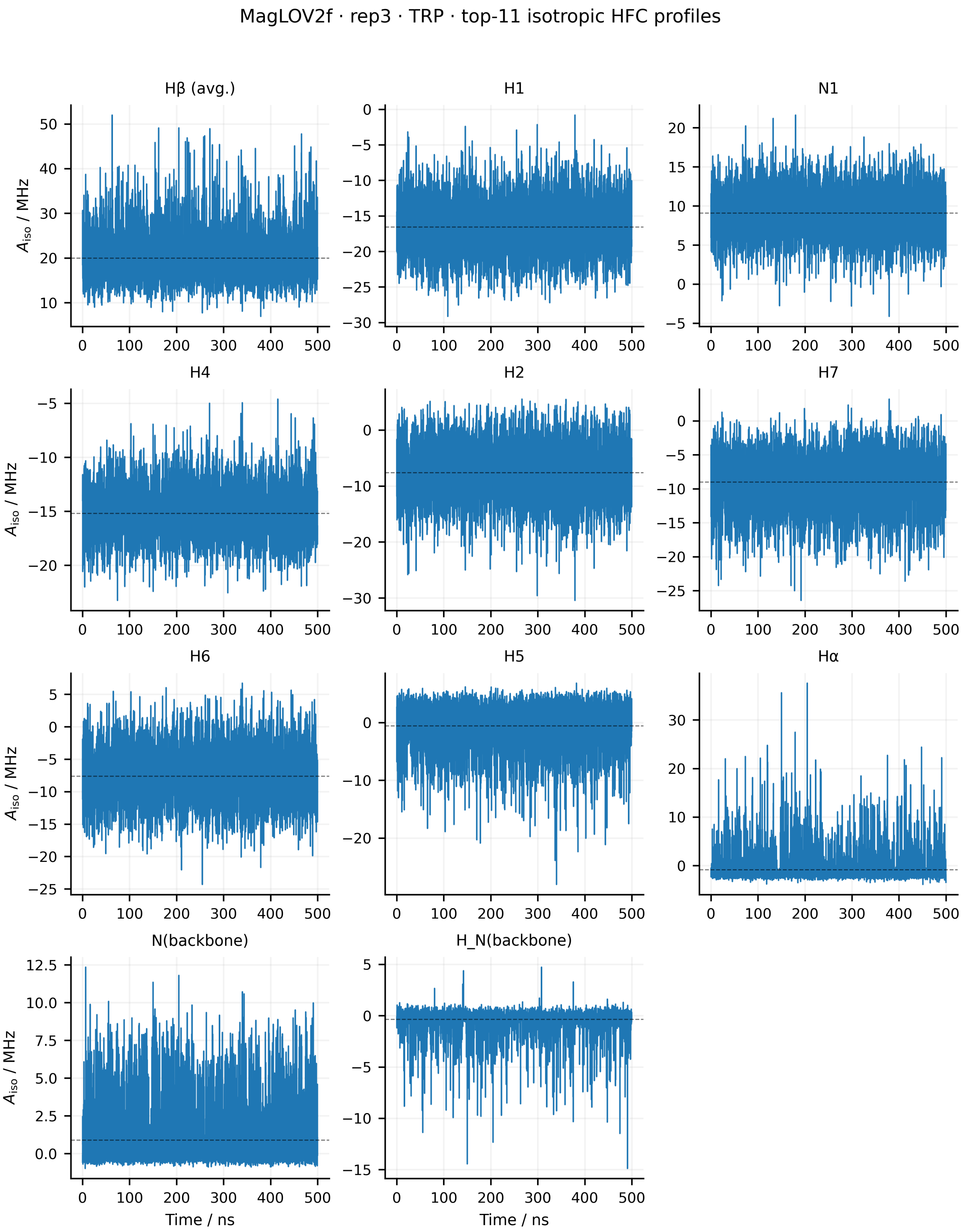


Figure S61: Time evolution of the isotropic hyperfine couplings of the 11 physical Trp HFC entities for replica 3 of MagLOV2f. The isotropic coupling was calculated as $A_{\mathrm{iso}}(t) = \mathrm{Tr}[\mathbf{A}(t)]/3$. Dashed horizontal lines indicate the mean value over the 500 ns trajectory. The two H$\beta$ couplings are averaged for visualisation, while the individual nuclear spins are retained in the relaxation analysis. Artificial capping hydrogens are excluded.

**Full hyperfine tensor fluctuations** The full Trp HFC-tensor fluctuations were evaluated using the same Frobenius-norm measure as for FMN. These trajectories retain fluctuations of all Cartesian tensor components and therefore contain the complete HFC modulation relevant to the random-field relaxation treatment. Comparison between replicas additionally reveals whether enhanced HFC modulation is persistent across independent trajectories or is associated with conformational substates sampled only in individual simulations.

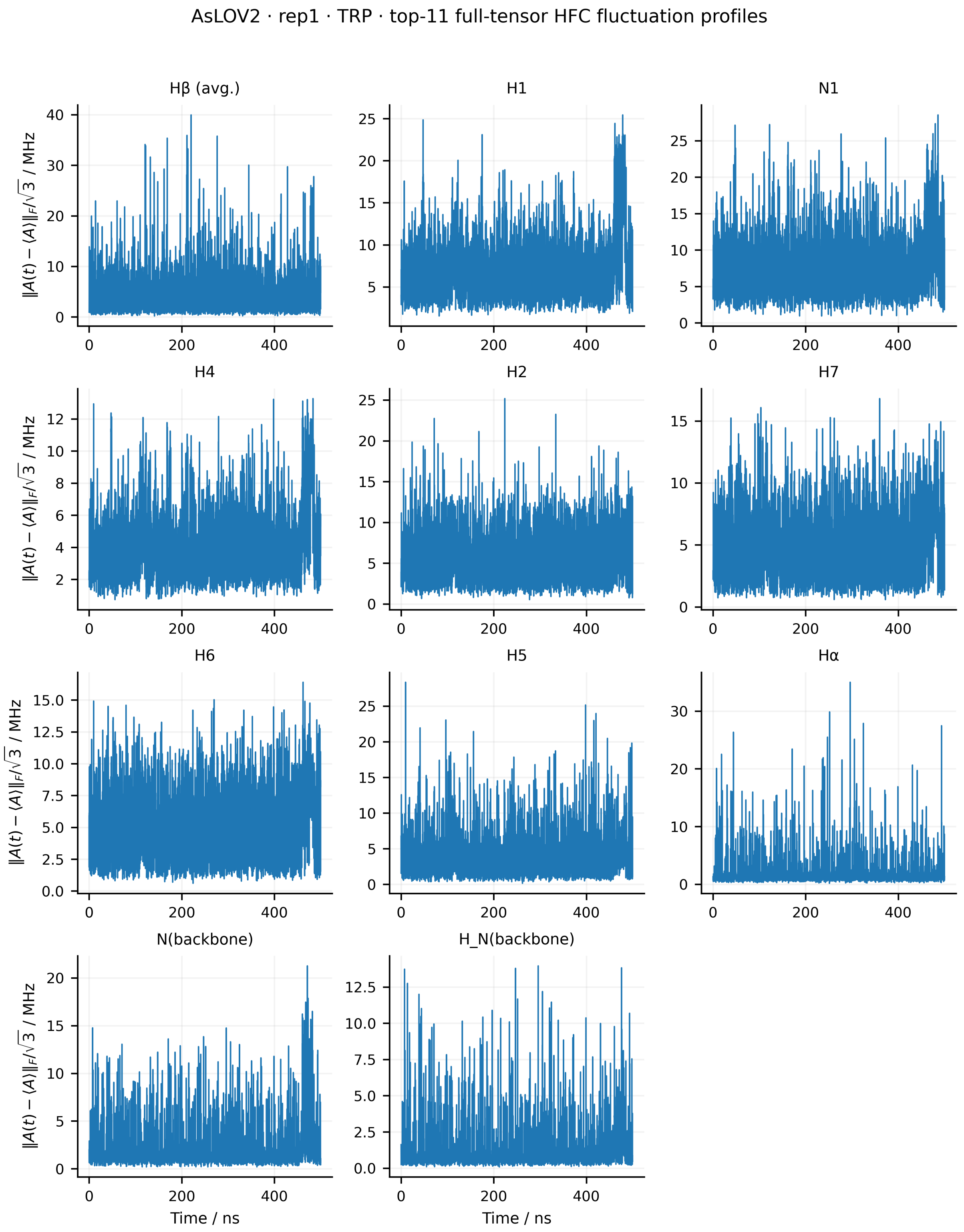


Figure S62: Full-tensor hyperfine-coupling fluctuations of the 11 physical Trp HFC entities for replica 1 of *As*LOV2. The instantaneous fluctuation amplitude is defined as $\|\mathbf{A}(t) - \langle\mathbf{A}\rangle\|_\mathrm{F}/\sqrt{3}$, where $\langle\mathbf{A}\rangle$ is the mean HFC tensor over the 500 ns trajectory. The two H$\beta$ couplings are averaged for visualisation only, and artificial capping hydrogens are excluded.

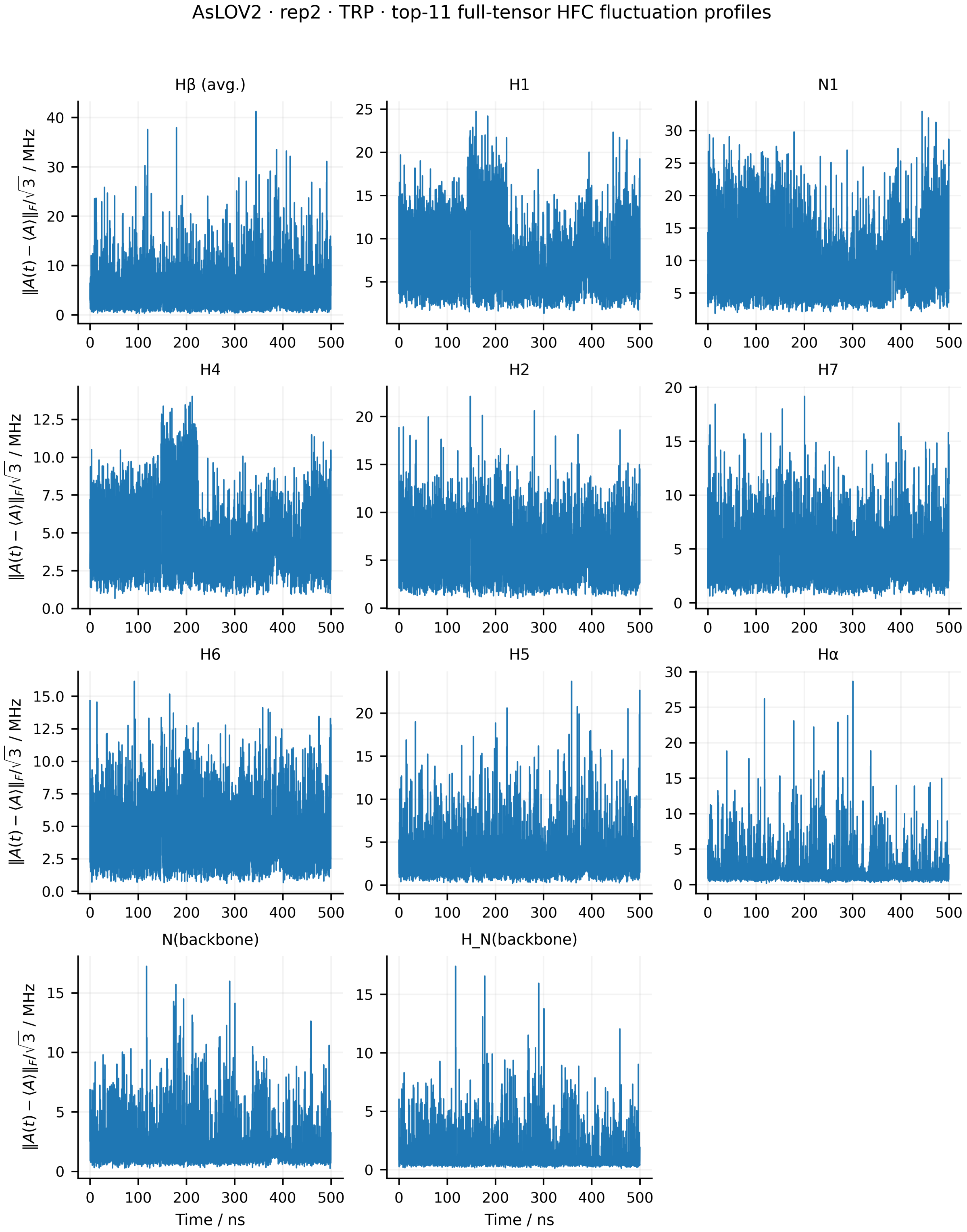


Figure S63: Full-tensor hyperfine-coupling fluctuations of the 11 physical Trp HFC entities for replica 2 of *As*LOV2. The instantaneous fluctuation amplitude is defined as $\|\mathbf{A}(t) - \langle\mathbf{A}\rangle\|_{\mathrm{F}}/\sqrt{3}$, where $\langle\mathbf{A}\rangle$ is the mean HFC tensor over the 500 ns trajectory. The two H$\beta$ couplings are averaged for visualisation only, and artificial capping hydrogens are excluded.

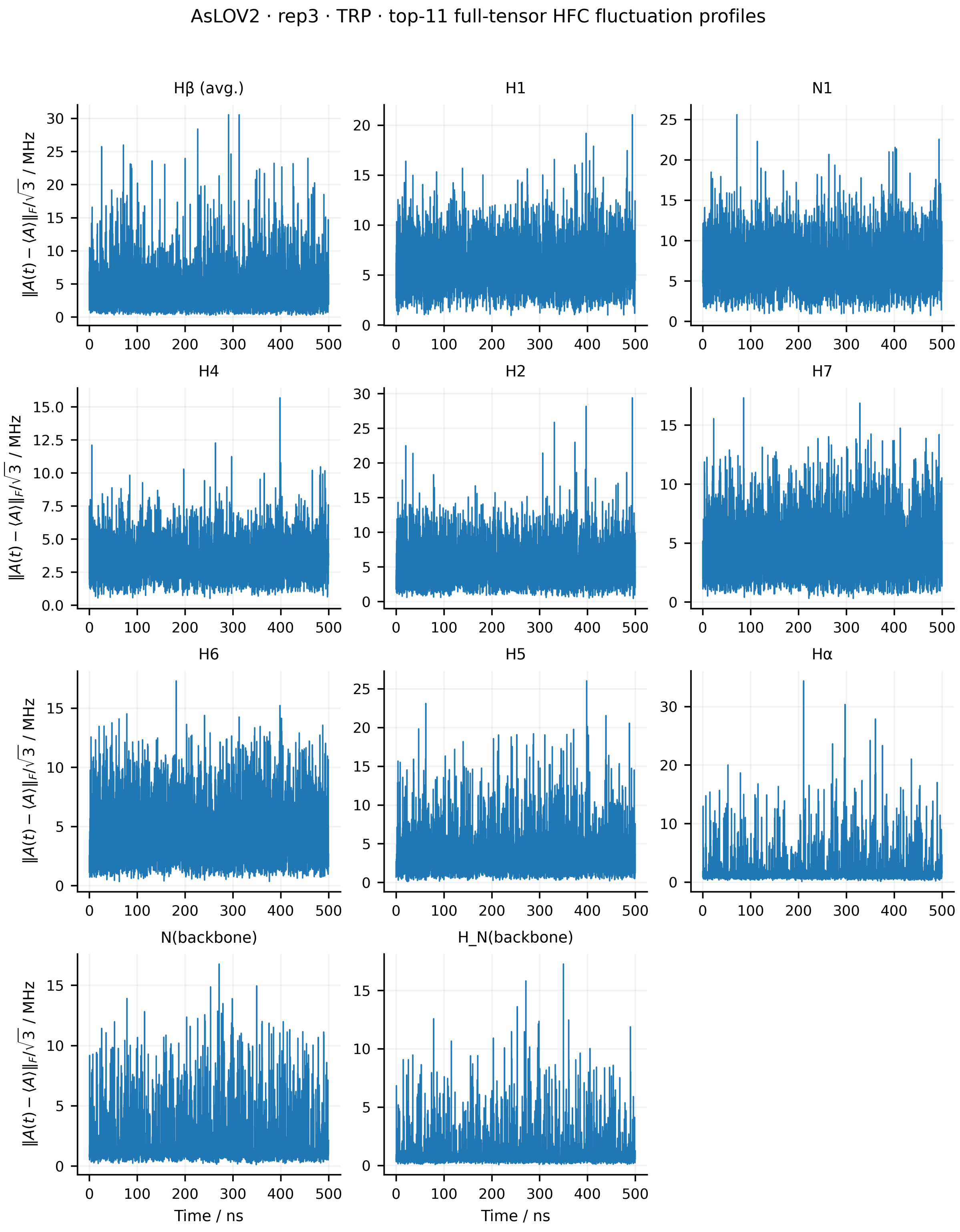


Figure S64: Full-tensor hyperfine-coupling fluctuations of the 11 physical Trp HFC entities for replica 3 of *As*LOV2. The instantaneous fluctuation amplitude is defined as $\|\mathbf{A}(t) - \langle \mathbf{A} \rangle\|_{\mathrm{F}}/\sqrt{3}$, where $\langle \mathbf{A} \rangle$ is the mean HFC tensor over the 500 ns trajectory. The two H$\beta$ couplings are averaged for visualisation only, and artificial capping hydrogens are excluded.

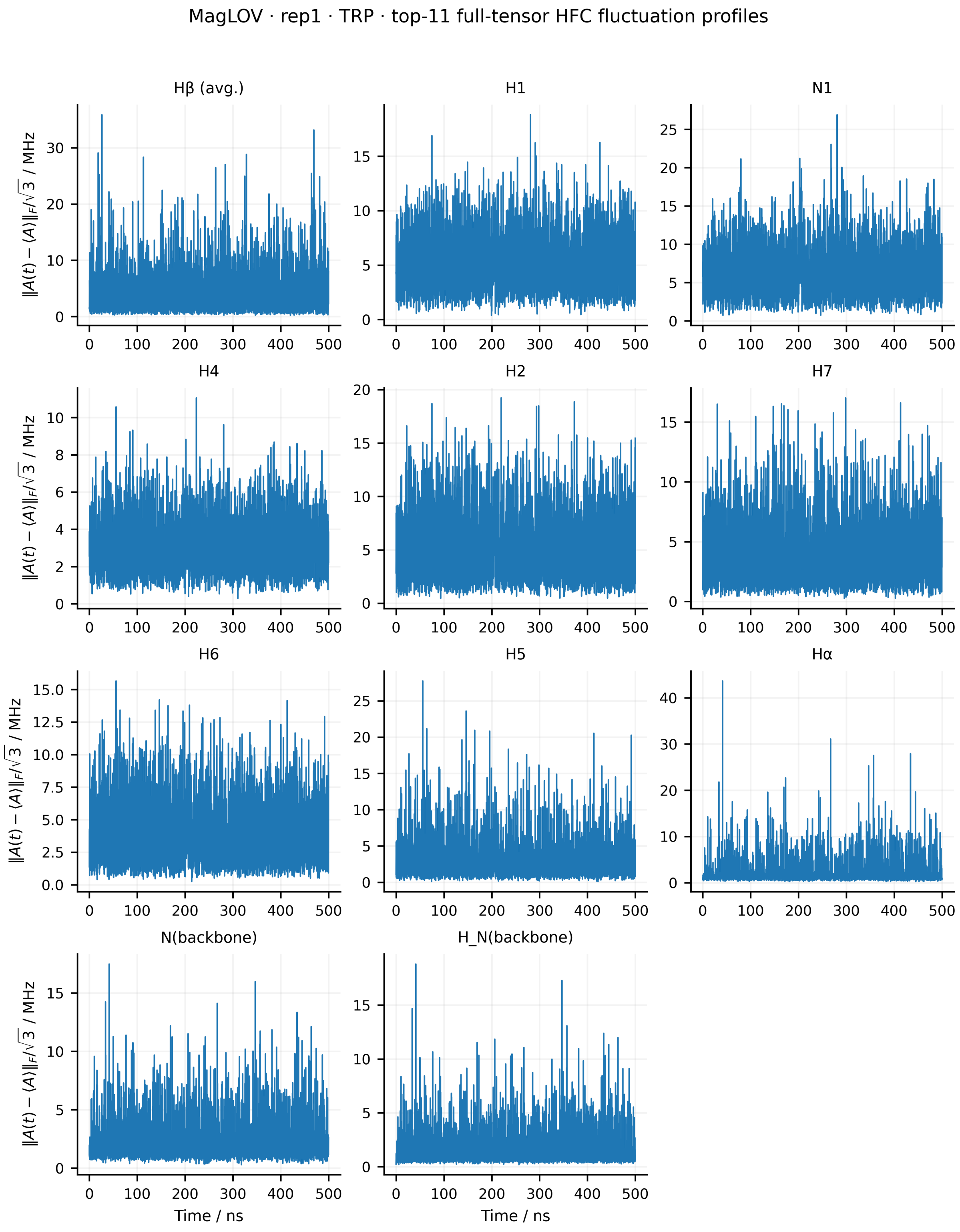


Figure S65: Full-tensor hyperfine-coupling fluctuations of the 11 physical Trp HFC entities for replica 1 of MagLOV. The instantaneous fluctuation amplitude is defined as $\|\mathbf{A}(t) - \langle\mathbf{A}\rangle\|_\mathrm{F}/\sqrt{3}$, where $\langle\mathbf{A}\rangle$ is the mean HFC tensor over the 500 ns trajectory. The two H$\beta$ couplings are averaged for visualisation only, and artificial capping hydrogens are excluded.

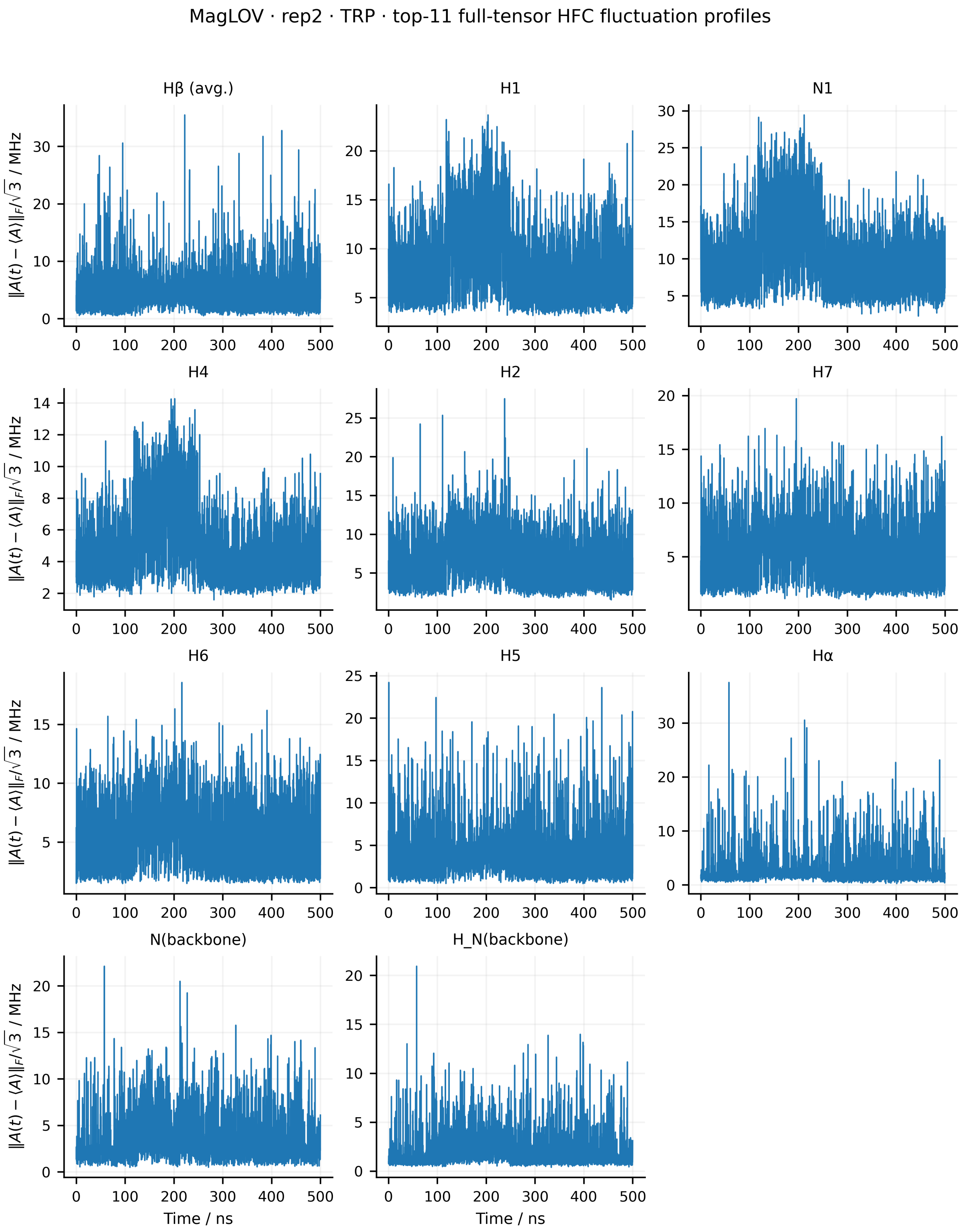


Figure S66: Full-tensor hyperfine-coupling fluctuations of the 11 physical Trp HFC entities for replica 2 of MagLOV. The instantaneous fluctuation amplitude is defined as $\|\mathbf{A}(t) - \langle\mathbf{A}\rangle\|_{\mathrm{F}}/\sqrt{3}$, where $\langle\mathbf{A}\rangle$ is the mean HFC tensor over the 500 ns trajectory. The two H$\beta$ couplings are averaged for visualisation only, and artificial capping hydrogens are excluded.

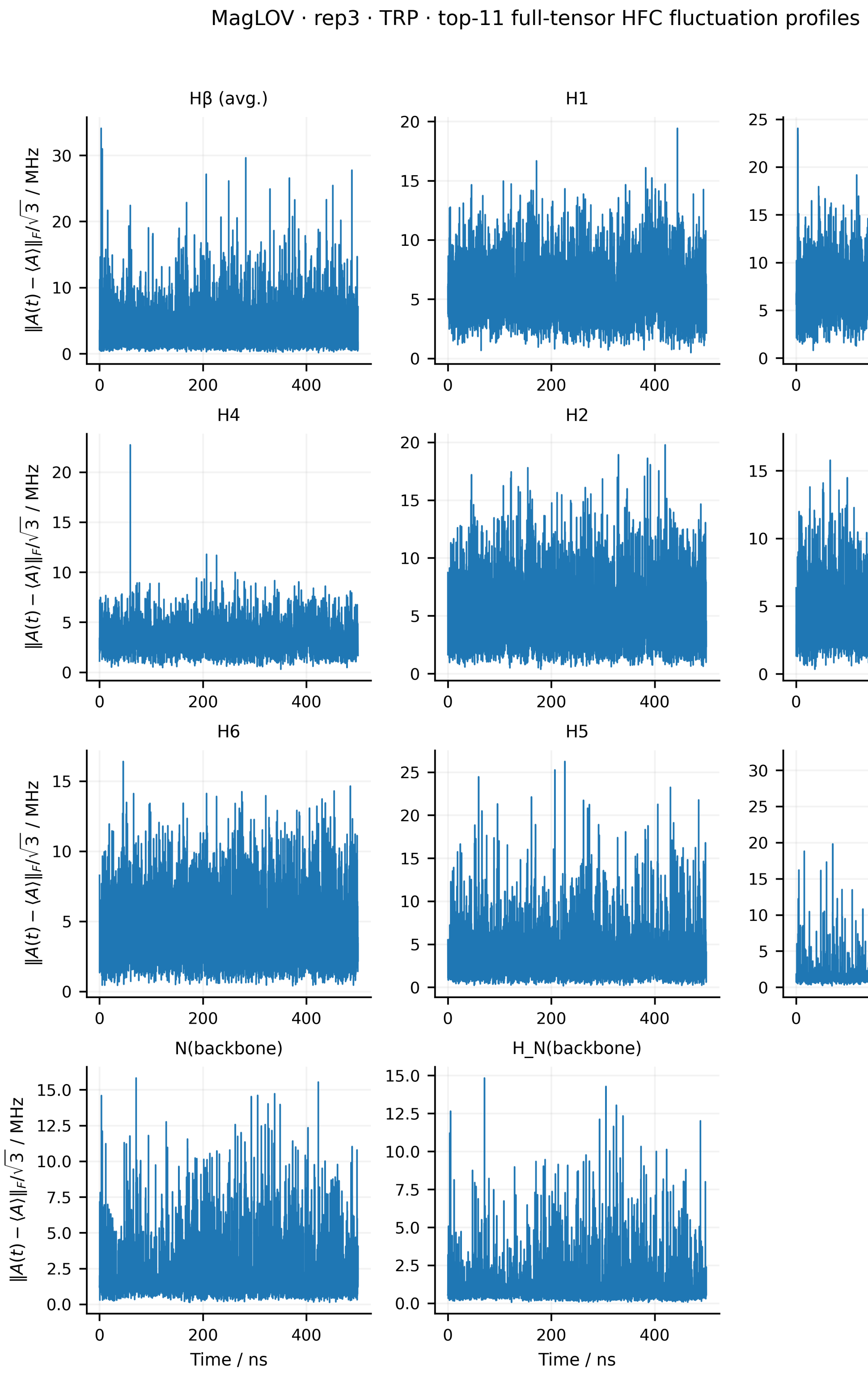


Figure S67: Full-tensor hyperfine-coupling fluctuations of the 11 physical Trp HFC entities for replica 3 of MagLOV. The instantaneous fluctuation amplitude is defined as $\|\mathbf{A}(t) - \langle \mathbf{A} \rangle\|_{\mathrm{F}}/\sqrt{3}$, where $\langle \mathbf{A} \rangle$ is the mean HFC tensor over the 500 ns trajectory. The two H$\beta$ couplings are averaged for visualisation only, and artificial capping hydrogens are excluded.

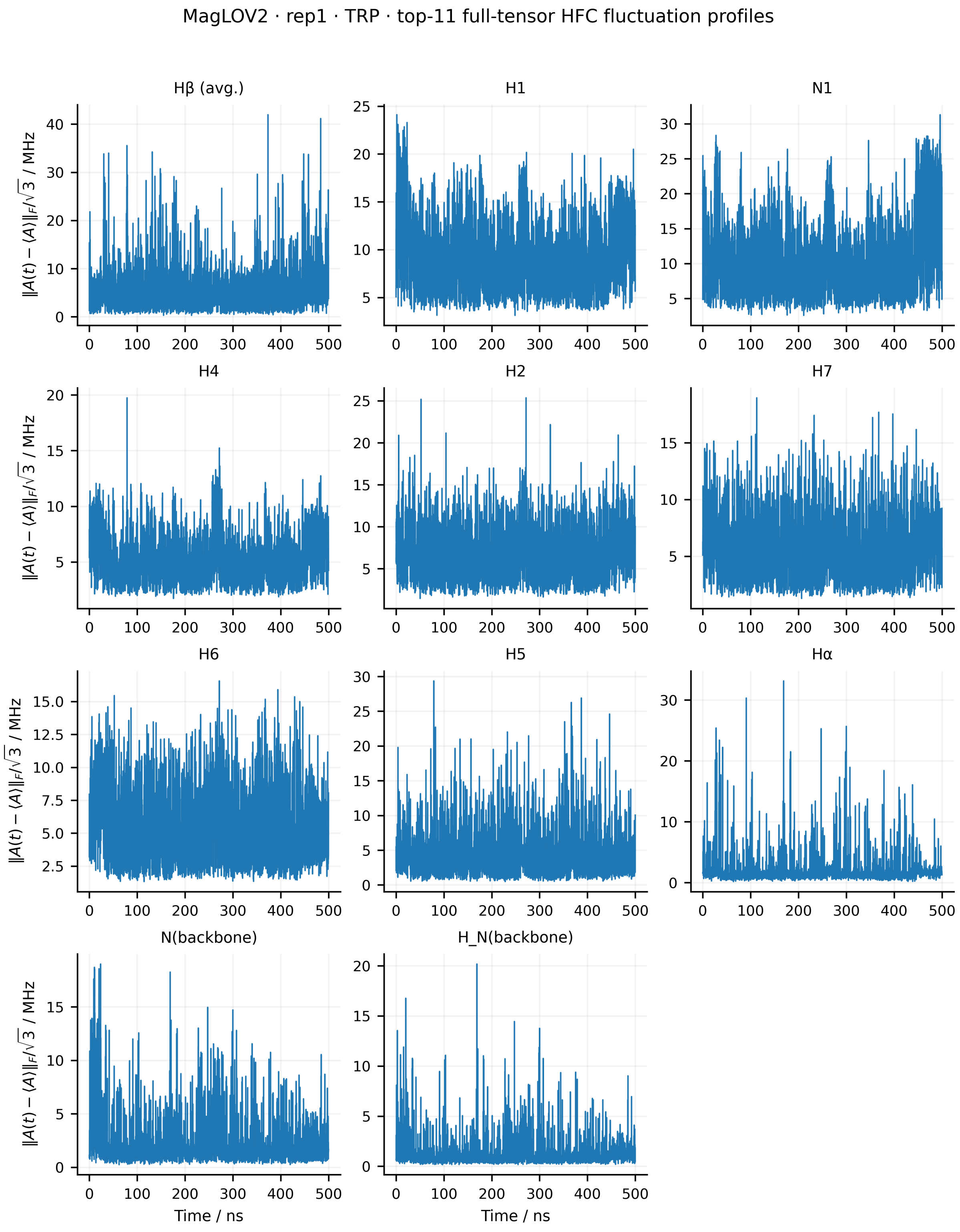


Figure S68: Full-tensor hyperfine-coupling fluctuations of the 11 physical Trp HFC entities for replica 1 of MagLOV2. The instantaneous fluctuation amplitude is defined as $\|\mathbf{A}(t) - \langle \mathbf{A} \rangle\|_{\mathrm{F}}/\sqrt{3}$, where $\langle \mathbf{A} \rangle$ is the mean HFC tensor over the 500 ns trajectory. The two H$\beta$ couplings are averaged for visualisation only, and artificial capping hydrogens are excluded.

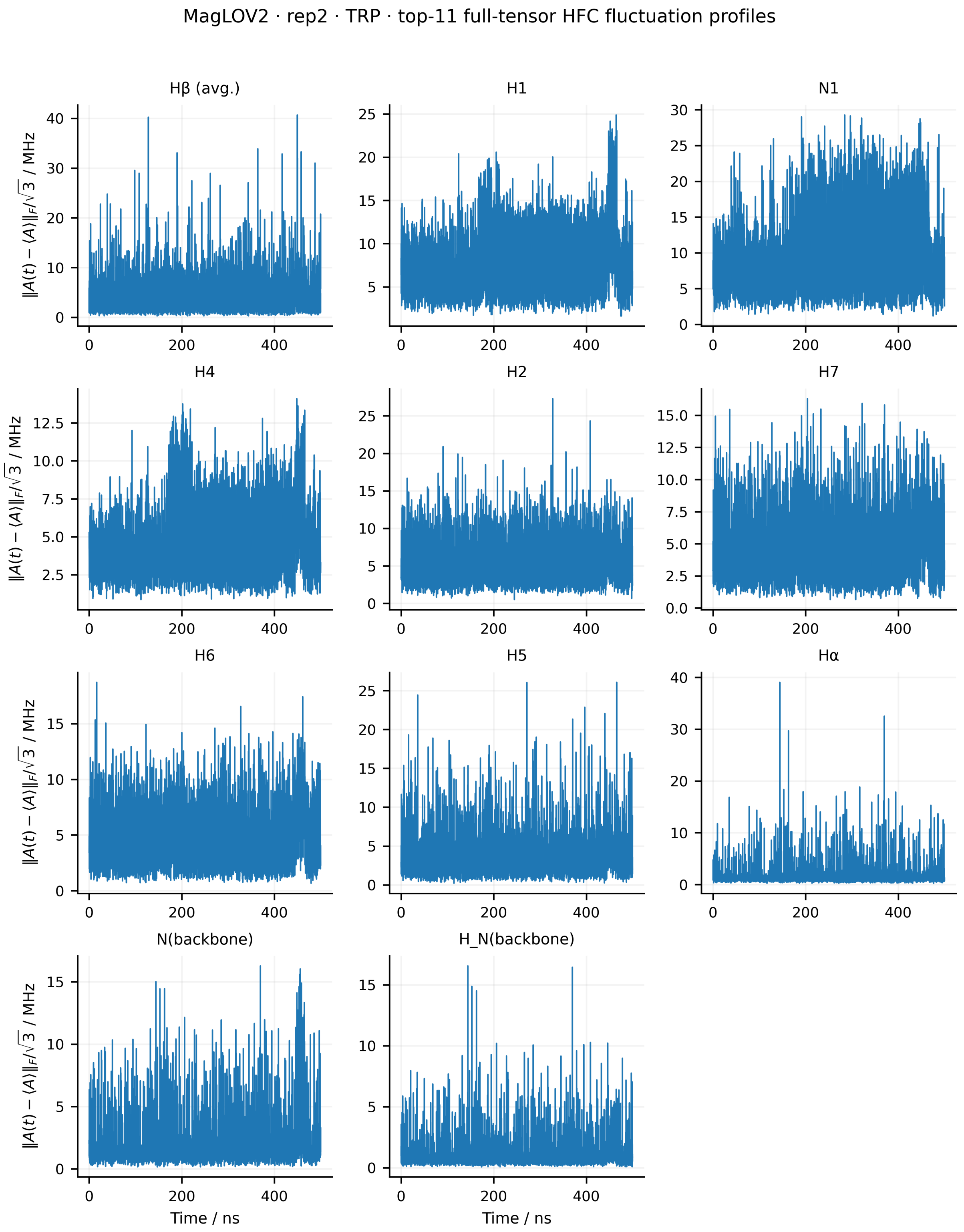


Figure S69: Full-tensor hyperfine-coupling fluctuations of the 11 physical Trp HFC entities for replica 2 of MagLOV2. The instantaneous fluctuation amplitude is defined as $\|\mathbf{A}(t) - \langle\mathbf{A}\rangle\|_{\mathrm{F}}/\sqrt{3}$, where $\langle\mathbf{A}\rangle$ is the mean HFC tensor over the 500 ns trajectory. The two H$\beta$ couplings are averaged for visualisation only, and artificial capping hydrogens are excluded.

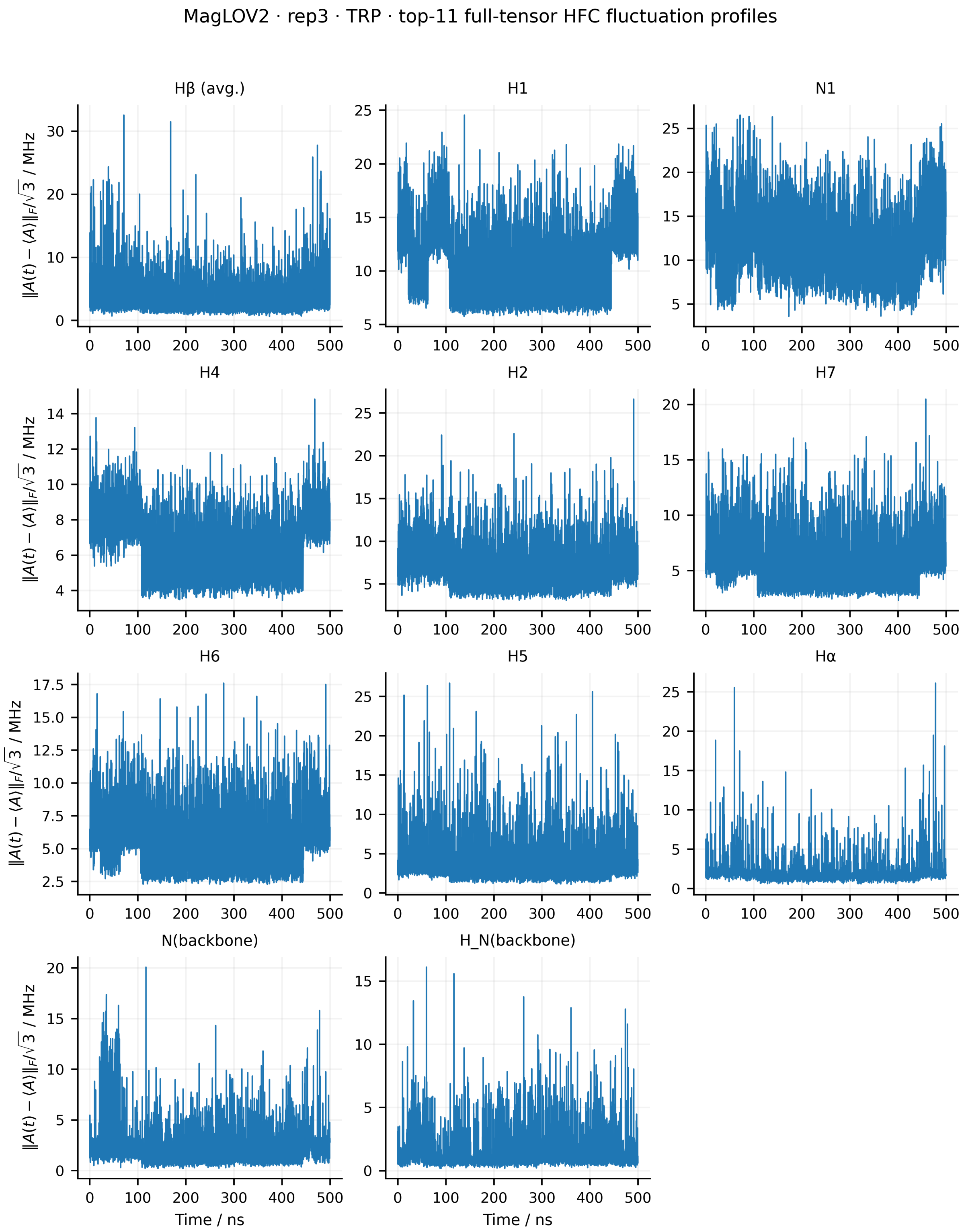


Figure S70: Full-tensor hyperfine-coupling fluctuations of the 11 physical Trp HFC entities for replica 3 of MagLOV2. The instantaneous fluctuation amplitude is defined as $\|\mathbf{A}(t) - \langle \mathbf{A} \rangle\|_F/\sqrt{3}$, where $\langle \mathbf{A} \rangle$ is the mean HFC tensor over the 500 ns trajectory. The two H$\beta$ couplings are averaged for visualisation only, and artificial capping hydrogens are excluded.

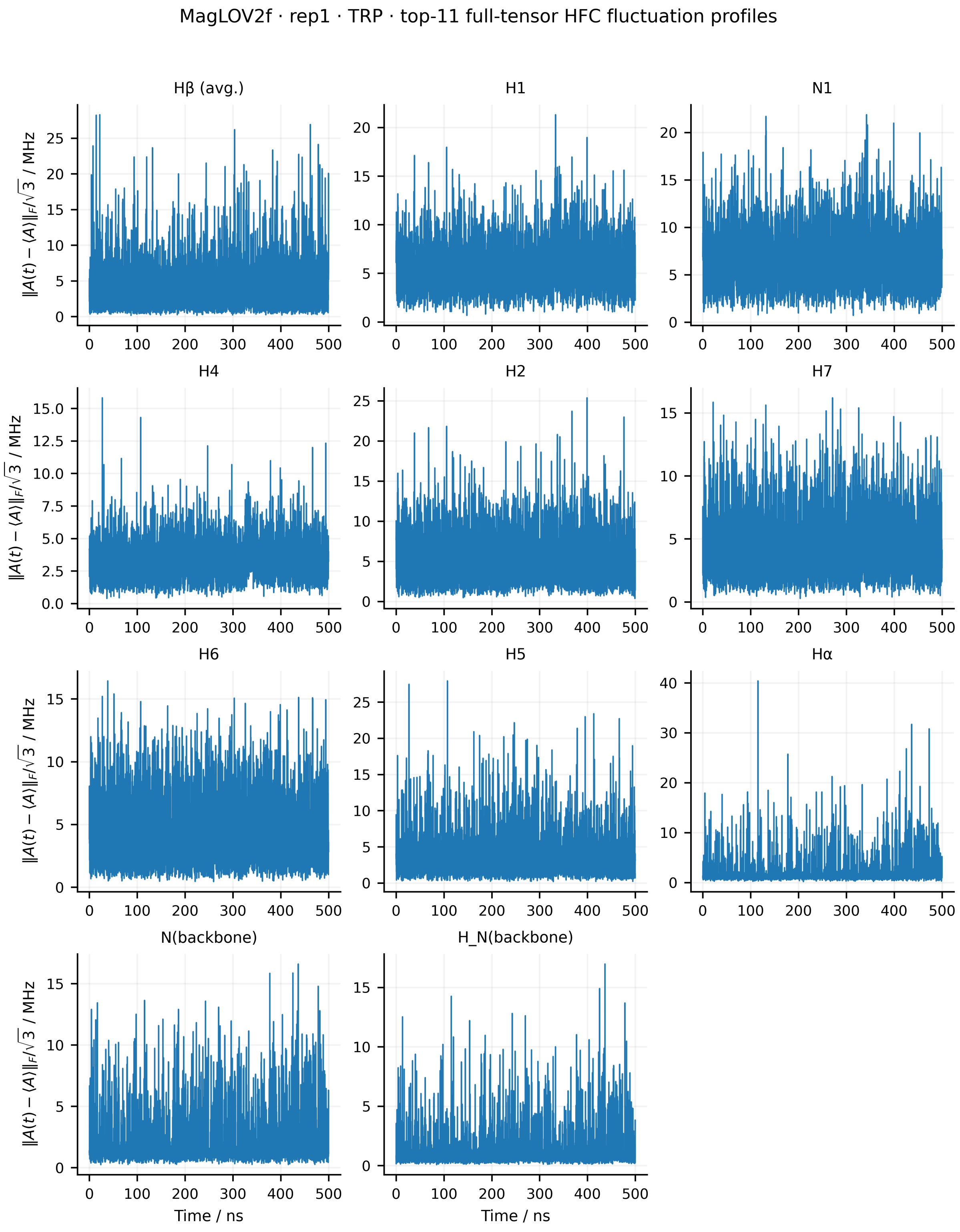


Figure S71: Full-tensor hyperfine-coupling fluctuations of the 11 physical Trp HFC entities for replica 1 of MagLOV2f. The instantaneous fluctuation amplitude is defined as $\|\mathbf{A}(t) - \langle\mathbf{A}\rangle\|_{\mathrm{F}}/\sqrt{3}$, where $\langle\mathbf{A}\rangle$ is the mean HFC tensor over the 500 ns trajectory. The two H$\beta$ couplings are averaged for visualisation only, and artificial capping hydrogens are excluded.

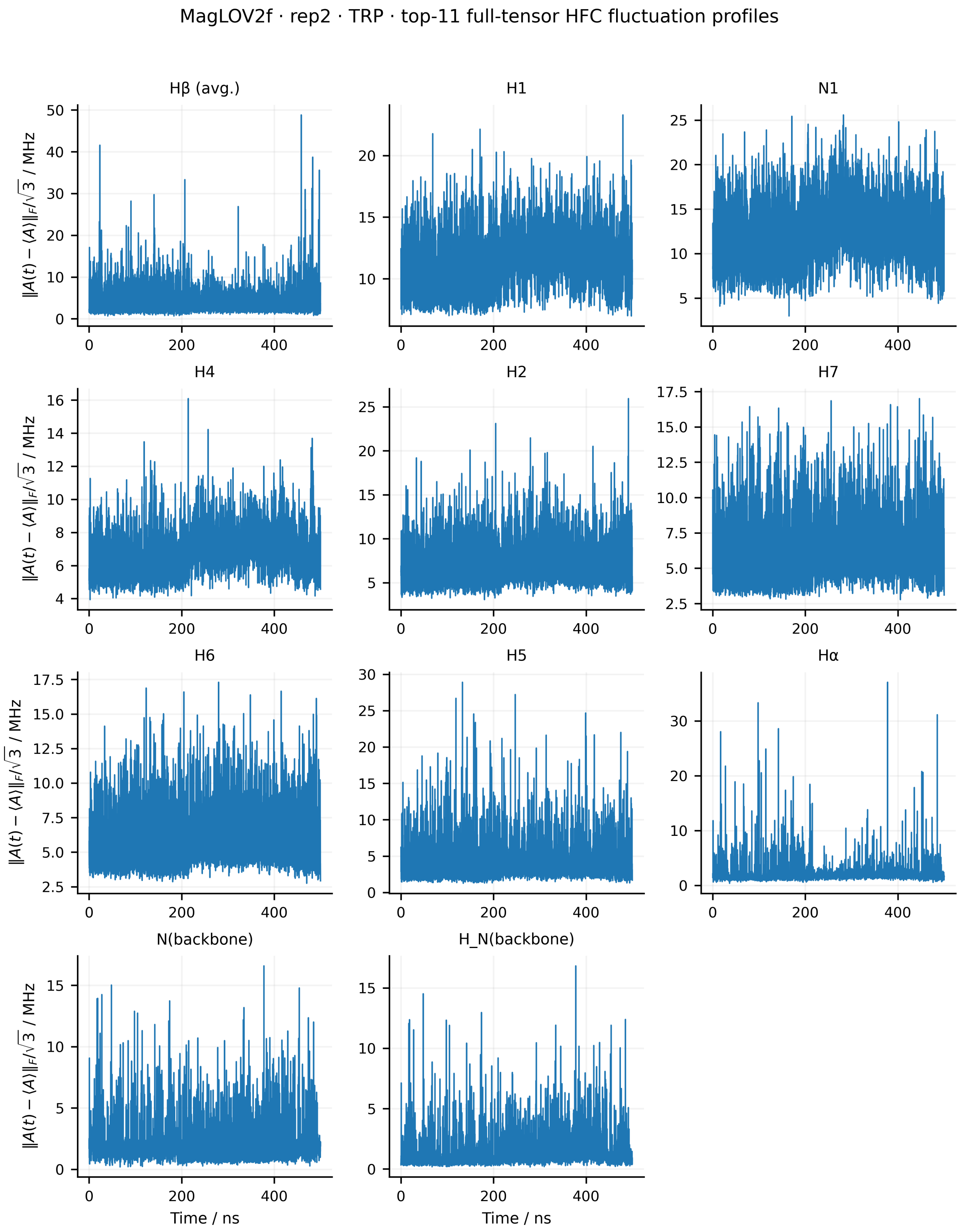


Figure S72: Full-tensor hyperfine-coupling fluctuations of the 11 physical Trp HFC entities for replica 2 of MagLOV2f. The instantaneous fluctuation amplitude is defined as $\|\mathbf{A}(t) - \langle\mathbf{A}\rangle\|_{\mathrm{F}}/\sqrt{3}$, where $\langle\mathbf{A}\rangle$ is the mean HFC tensor over the 500 ns trajectory. The two H$\beta$ couplings are averaged for visualisation only, and artificial capping hydrogens are excluded.

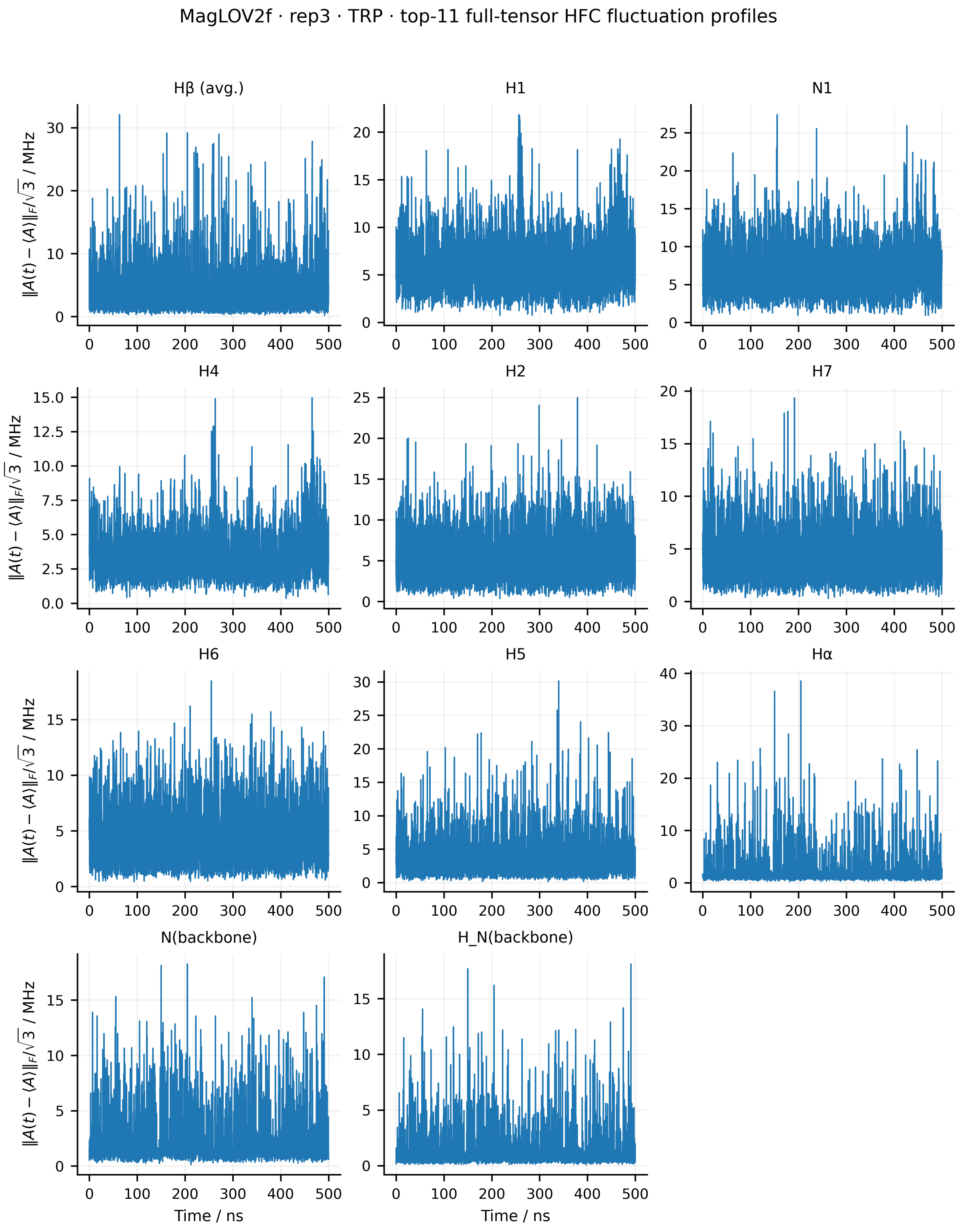


Figure S73: Full-tensor hyperfine-coupling fluctuations of the 11 physical Trp HFC entities for replica 3 of MagLOV2f. The instantaneous fluctuation amplitude is defined as $\|\mathbf{A}(t) - \langle \mathbf{A} \rangle\|_F/\sqrt{3}$, where $\langle \mathbf{A} \rangle$ is the mean HFC tensor over the 500 ns trajectory. The two H$\beta$ couplings are averaged for visualisation only, and artificial capping hydrogens are excluded.

### S9 Back electron transfer rate constants

For the native donor pathway considered here, the charge-separated radical pair is $FMN^{\bullet-}$ / $W89H^{\bullet+}$. Throughout this section, GS denotes the electronic ground state and RP the charge-separated radical-pair state. Back electron transfer (BET) is therefore the RP→GS reaction.

#### Energy-gap reaction coordinate

For a nuclear configuration $\mathbf{q}$, the vertical diabatic energy gap was defined with one sign convention throughout,

$$\Delta E(\mathbf{q}) = E_{\mathrm{RP}}(\mathbf{q}) - E_{\mathrm{GS}}(\mathbf{q}). \tag{S9}$$

Both electronic-state energies were evaluated on configurations sampled from each equilibrium ensemble. This gives two time series,

$$\Delta E_{\mathrm{GS}}(t) = E_{\mathrm{RP|GS}}(t) - E_{\mathrm{GS|GS}}(t), \tag{S10}$$

$$\Delta E_{\mathrm{RP}}(t) = E_{\mathrm{RP|RP}}(t) - E_{\mathrm{GS|RP}}(t), \tag{S11}$$

where $E_{A|B}$ denotes the energy of the electronic state $A$ at a geometry drawn from the ensemble of states $B$. The corresponding distributions are denoted $P_{\mathrm{GS}}(\Delta E)$ and $P_{\mathrm{RP}}(\Delta E)$.[4,5] This notation avoids assigning "reactant" and "product" labels to the two gap distributions, which can otherwise obscure the sign convention for the RP→GS reaction.

The classical force-field energies of different redox states can contain systematic state-dependent offsets. A subtractive multilayer correction, following the general ONIOM/QM–MM replacement logic,[6,7] was therefore applied:

$$E^{(s)}_{\mathrm{corr}}(t) = E^{(s)}_{\mathrm{MM,all}}(t) - \left[E^{(s)}_{\mathrm{MM,FMN}}(t) + E^{(s)}_{\mathrm{MM,Trp}}(t)\right] + \left[\overline{E}^{(s)}_{\mathrm{QC,FMN}} + \overline{E}^{(s)}_{\mathrm{QC,Trp}}\right], \tag{S12}$$

where $s \in \{\mathrm{GS}, \mathrm{RP}\}$. The quantum-chemical fragment terms are state-specific averages over 30 representative structures, using the electronic-structure protocol described in the main-text Methods. Consequently, this correction improves the relative state offset but does not introduce frame-resolved quantum-chemical fluctuations of the fragments. The absolute thermodynamic quantities obtained from this hybrid construction are therefore model-dependent; comparisons among the four variants are the principal use of the model.

Under the Gaussian linear-response approximation, the two energy-gap distributions generate parabolic diabatic free-energy surfaces.[4,5,8] We define

$$\mu_{\mathrm{GS}} = \langle \Delta E_{\mathrm{GS}} \rangle, \qquad \mu_{\mathrm{RP}} = \langle \Delta E_{\mathrm{RP}} \rangle, \tag{S13}$$

$$\sigma^2_{\mathrm{GS}} = \mathrm{Var}(\Delta E_{\mathrm{GS}}), \qquad \sigma^2_{\mathrm{RP}} = \mathrm{Var}(\Delta E_{\mathrm{RP}}), \tag{S14}$$

and use the common variance

$$\sigma^2 = \frac{1}{2}\left(\sigma^2_{\mathrm{GS}} + \sigma^2_{\mathrm{RP}}\right). \tag{S15}$$

The corresponding surfaces are

$$G_{\mathrm{GS}}(\Delta E) = \frac{k_{\mathrm{B}}T}{2\sigma^2}(\Delta E - \mu_{\mathrm{GS}})^2 + C_{\mathrm{GS}}, \tag{S16}$$

$$G_{\mathrm{RP}}(\Delta E) = \frac{k_{\mathrm{B}}T}{2\sigma^2}(\Delta E - \mu_{\mathrm{RP}})^2 + C_{\mathrm{RP}}. \tag{S17}$$

For the gap definition in Eq. (S9), the two diabatic surfaces are degenerate at $\Delta E = 0$. Their additive constants were therefore chosen so that $G_{\mathrm{GS}}(0) = G_{\mathrm{RP}}(0)$; a common constant was then subtracted so that the GS minimum is zero. The reorganisation energy obtained from the common-curvature parabolas is

$$\lambda_r = G_{\mathrm{GS}}(\mu_{\mathrm{RP}}) - G_{\mathrm{GS}}(\mu_{\mathrm{GS}}) = \frac{k_{\mathrm{B}}T}{2\sigma^2}\left(\mu_{\mathrm{GS}} - \mu_{\mathrm{RP}}\right)^2, \tag{S18}$$

and the recombination free energy is the difference between the minima,

$$\Delta G^{\circ}_{\mathrm{RP}\to\mathrm{GS}} = G_{\mathrm{GS}}(\mu_{\mathrm{GS}}) - G_{\mathrm{RP}}(\mu_{\mathrm{RP}}). \tag{S19}$$

For ideal linear response with exactly equal curvature, the same sign convention gives the useful moment relations

$$\Delta G^{\circ}_{\mathrm{RP}\to\mathrm{GS}} = -\frac{\mu_{\mathrm{GS}} + \mu_{\mathrm{RP}}}{2}, \qquad \lambda_{\mathrm{LR}} = \frac{\mu_{\mathrm{GS}} - \mu_{\mathrm{RP}}}{2}. \tag{S20}$$

The rate calculations reported here use $\lambda_r$ and $\Delta G^{\circ}_{\mathrm{RP}\to\mathrm{GS}}$ from the constructed parabolas, maintaining internal consistency with Fig. 5 of the main manuscript.

#### Marcus rate and geometry-dependent coupling

The nonadiabatic BET rate was evaluated using semiclassical Marcus theory,[8,9]

$$k_{\mathrm{BET}}(R,\alpha) = \frac{2\pi}{\hbar}|V(R,\alpha)|^2 \frac{1}{\sqrt{4\pi\lambda_r k_{\mathrm{B}}T}} \exp\left[-\frac{(\Delta G^{\circ}_{\mathrm{RP}\to\mathrm{GS}} + \lambda_r)^2}{4\lambda_r k_{\mathrm{B}}T}\right]. \tag{S21}$$

Here $V(R,\alpha)$ is an effective donor–acceptor electronic coupling. Direct diabatic coupling calculations would be preferable for absolute rate prediction, but they are considerably more demanding.[10–12] We therefore used the empirical Moser–Dutton protein electron-transfer ruler to set the baseline coupling scale,[13]

$$\log_{10}\left(\frac{k_{\mathrm{ref}}}{\mathrm{s}^{-1}}\right) = 13 - 0.6\,(R_{\mathrm{ref}} - 3.6) - 3.1\,\frac{(\Delta G^{\circ} + \lambda_r)^2}{\lambda_r}, \tag{S22}$$

with $R_{\mathrm{ref}}$ in Å and the energy terms in eV. The reference coupling is then

$$|V_{\mathrm{ref}}| = \sqrt{\frac{k_{\mathrm{ref}}\hbar}{2\pi F_{\mathrm{FC}}}}, \qquad F_{\mathrm{FC}} = \frac{1}{\sqrt{4\pi\lambda_r k_{\mathrm{B}}T}} \exp\left[-\frac{(\Delta G^{\circ} + \lambda_r)^2}{4\lambda_r k_{\mathrm{B}}T}\right]. \tag{S23}$$

Because aromatic $\pi$-orbital overlap is orientation dependent,[14] the sampled angle $\alpha$ between the undirected normals of the FMN and W89 aromatic planes was included phenomenologically as

$$V(R, \alpha) = V_{\text{ref}} \exp[-\beta(R - R_{\text{ref}})] \, |\cos \alpha|^m. \tag{S24}$$

The $|\cos \alpha|$ form captures the leading twist dependence expected for coupled conjugated $\pi$ systems,[15] and the absolute value makes the result invariant to the arbitrary sign of a plane normal. In the reported comparison, the distance attenuation already contained in the Moser–Dutton baseline was not applied a second time at the frame level ($\beta = 0$), and $m = 1$ was used; hence, the trajectory-level orientation factor contributes as $\cos^2 \alpha$ to the rate. The resulting frame-wise rates were averaged over the MD ensemble. This orientation factor is an empirical surrogate for an explicit diabatic coupling and is a principal source of uncertainty in the absolute $k_{\text{BET}}$ values.

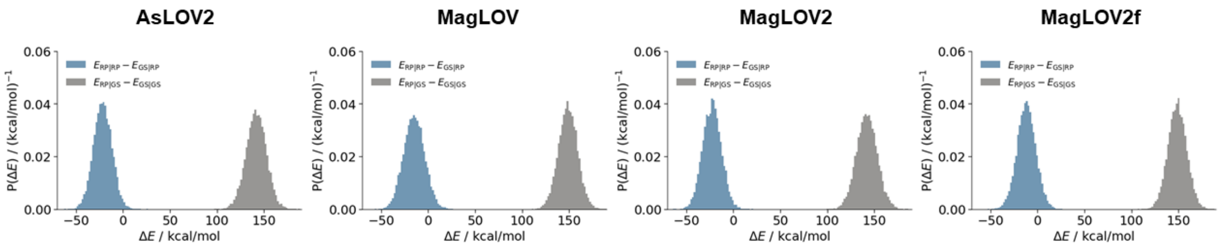


Figure S74: Energy-gap distributions $P(\Delta E)$ evaluated on the RP and GS nuclear ensembles of *As*LOV2, MagLOV, MagLOV2, and MagLOV2f. Blue distributions show $E_{\text{RP|RP}} - E_{\text{GS|RP}}$ and grey distributions show $E_{\text{RP|GS}} - E_{\text{GS|GS}}$, using the common sign convention of Eq. (S9). Their means and variances determine the comparative free-energy profiles used in the Marcus analysis.

The two state-conditioned gap distributions are well separated in every variant (Fig. S74), consistent with substantial solvent/protein reorganisation between the GS and RP ensembles. Their widths quantify thermal fluctuations within the corresponding classical ensembles, and their displacement controls the linear-response estimates of $\lambda_r$ and $\Delta G^\circ$. The separation itself should not be interpreted as independent evidence for a particular radical-pair lifetime; kinetic stability is determined by the resulting free-energy barrier together with the electronic coupling.

The four constructs retain the same qualitative two-ensemble structure, while the positions and widths of the distributions vary. These changes, combined with the mutation-dependent aromatic-plane-angle distributions (Fig. S18) and the closely similar mean COM separations (Fig. S23), result in the variant-dependent BET rates discussed in the main manuscript. Because both the hybrid energy correction and the orientation-dependent coupling are approximate, the rate ordering and relative shifts are interpreted more robustly than the absolute numerical rates.
